# Promoting Healthy Ageing through Choral Training: A 9-Month Multidomain Intervention (MultiMusic) Enhances Neural Processing Speed in Community-Dwelling Older Adults

**DOI:** 10.64898/2026.03.30.715236

**Authors:** M. Lippolis, A. Pantaleo, L. Mazzon, R. Diomede, M. Delussi, E. Seminerio, N. Quaranta, A. Pilotto, V. Solfrizzi, P. Vuust, E. Brattico

## Abstract

**Background:** Older adulthood is often accompanied by declines in auditory processing and cognitive functioning, increasing the risk of reduced autonomy and quality of life. Multidomain lifestyle interventions have shown potential to counteract these changes, and choir-based activities represent a promising approach by simultaneously engaging auditory, cognitive, physical, and social domains. However, evidence regarding their feasibility and neurophysiological impact in community-dwelling older adults, particularly those without formal musical training, remains scarce.

**Methods:** This 9-month quasi-experimental feasibility study involved 54 community-dwelling older adults (mean age = 72.9 years) with no formal musical background. Participants self-selected into a choir-based intervention group, an active control group engaging in non-musical leisure activities, or a passive control group; however, some participants in the control groups were selected from the waiting list for the choir. Assessments were conducted at baseline and follow-up and included measures of global cognition, cognitive reserve, psychological well-being (Flourishing Scale), multidimensional frailty (Selfy-MPI), music perception, pure-tone audiometry, and auditory evoked potentials recorded using a standardized clinical oddball paradigm.

**Results:** The choir-based intervention was feasible in a community setting. At the neurophysiological level, choir participation was associated with a bilateral, significant shortening of the N2–P3 inter-peak latency, indicating faster auditory–cortical processing. Additionally, through explorative analyses multidimensional frailty, as assessed by the Selfy-MPI, showed a significant reduction in individuals engaging in a higher number of activities, irrespective of group allocation. Similarly, psychological well-being revealed a decrease in flourishing scores in the passive control group relative to the choir group. No changes were observed in audiometric thresholds or music perception measures.

**Conclusion:** Choir-based multidomain participation is a feasible intervention for community-dwelling older adults without formal musical training and is associated with selective benefits in cognitive reserve, psychological well-being, auditory–cortical processing speed, and multidimensional frailty. These findings provide a foundation for a larger randomized controlled trial aimed at clarifying the cognitive, psychosocial, and neural mechanisms underlying choir-based interventions in ageing.

**Trial Registration:** The upcoming trial has been prospectively registered on ClinicalTrials.gov (ID: NCT06767410; registration date: January 9, 2025).

## Introduction

Ageing is a complex and multidimensional process characterized by biological, psychological, and social changes across the life span. In later adulthood, declines in processing speed, attentional control, and working memory often compromise everyday functioning and independence (Harada et al., 2013; Park & Festini, 2017). Importantly, cognitive ageing trajectories are highly heterogeneous: while some individuals preserve cognitive functioning into advanced age, others experience accelerated decline leading to mild cognitive impairment (MCI) or dementia. This variability highlights the need to identify protective factors and interventions capable of extending healthy cognitive ageing.

Among the most pervasive contributors to late-life vulnerability is sensory decline, particularly age-related hearing loss. Affecting nearly one third of adults over 65 and up to two thirds over 70 (Lin et al., 2011), hearing loss has been identified as a leading modifiable risk factor for dementia (Livingston et al., 2020). Beyond reduced audibility, degraded auditory input increases cognitive load on attentional and memory systems (Peelle & Wingfield, 2016) and promotes social withdrawal, thereby reducing engagement in cognitively stimulating activities (Mick et al., 2014). Neuroimaging evidence further links hearing loss to atrophy in auditory and prefrontal regions (Lin et al., 2014; Uchida et al., 2018), supporting the concept of a “cognitive ear,” whereby sensory degradation contributes silently to cognitive vulnerability in ageing (Sardone et al., 2019).

In response, growing attention has shifted toward non-pharmacological strategies to promote healthy and active ageing. Lifestyle-based approaches involving physical activity, cognitive stimulation, and social engagement are consistently associated with better brain health (Colcombe & Kramer, 2003; Hertzog et al., 2008; Kelly et al., 2014). These approaches have been formalized within multidomain intervention models, which acknowledge that age-related vulnerability emerges from the interaction of cognitive, sensory, physical, and psychosocial factors. Large-scale trials provide compelling evidence for this framework: the Finnish Geriatric Intervention Study to Prevent Cognitive Impairment and Disability (FINGER) demonstrated that multidomain programs can slow cognitive decline and reduce dementia risk, particularly in cognitively intact individuals or those at the MCI stage, when neuroplastic potential remains substantial (Ngandu et al., 2015; Rosenberg et al., 2020; Perus et al., 2022). This evidence has informed international initiatives, including WHO guidelines, which identify multidomain lifestyle interventions as central to dementia prevention (WHO, 2019).

This perspective aligns closely with the construct of multidimensional frailty, which conceptualizes vulnerability as the cumulative burden of deficits across multiple domains rather than a purely physical condition (Rockwood & Mitnitski, 2007; Pilotto et al., 2008, 2009). Multidomain interventions are therefore particularly well suited to address the complexity of ageing-related decline and to support cognitive reserve, defined as the capacity to maintain cognitive performance through flexible recruitment of neural resources in the face of brain ageing (Stern, 2002; 2012; Barulli & Stern, 2013).

Within this framework, music has emerged as a particularly promising form of enrichment. Music engagement simultaneously recruits auditory perception, attention, working memory, motor coordination, emotion regulation, and social interaction (Kraus & Chandrasekaran, 2010; Herholz & Zatorre, 2012), leading it to be described as “aerobics for the brain.” Converging evidence from developmental, cross-sectional, and longitudinal studies shows that musical training induces structural and functional brain adaptations, including changes in auditory and motor cortices and more efficient sound encoding (Gaser & Schlaug, 2003; Hyde et al., 2009; Habibi et al., 2016; Schlaug, 2015). A large meta-analysis further documented consistent differences in auditory, sensorimotor, interoceptive, and limbic regions between musicians and non-musicians (Criscuolo et al., 2022).

In older adults, music represents a feasible and socially rewarding activity that may overcome adherence limitations observed in other training programs (Davidson & Faulkner, 2010; Creech et al., 2013). Musical engagement has been associated not only with improved auditory skills but also with benefits in executive control, memory, and speech-in-noise perception (Bugos et al., 2007; Bialystok & DePape, 2009; Särkämö et al., 2014; Zendel & Alain, 2012; Slater et al., 2015). Crucially, randomized controlled trials (RCT) provide causal support for these associations: structured musical training has been shown to enhance cognitive and auditory functioning in healthy older adults (James et al., 2014; James et al., 2020), and long-term interventions in musically naïve participants lead to robust acquisition of musical skills with implications for cognitive and psychosocial outcomes (Losch et al., 2024). Together, these RCTs motivate the investigation of music-based interventions in later adulthood, consistent with models of experience-dependent learning and neuroplasticity proposed in the music neuroscience literature (Altenmüller & Schlaug, 2015).

Electrophysiological evidence further supports the role of music as a model of neural plasticity in ageing. EEG studies have shown that musically trained individuals exhibit more precise auditory encoding, enhanced temporal processing, and reduced age-related neural slowing (Parbery-Clark et al., 2011; Bidelman et al., 2015; Sanju & Kumar, 2016; Pentikäinen et al., 2022; Bonetti et al., 2024). Within auditory oddball paradigms, event-related potentials such as the N2 and P3 reflect successive stages of processing, from the detection of deviant stimuli to the allocation of attentional resources and updating of the stimulus context (Polich, 2007). Ageing is typically associated with delayed latencies and reduced amplitudes of these components, indicating slower and less efficient processing (Cheng et al., 2019; Shubhadarshan & Gaiwale, 2024). In contrast, musical engagement has been shown to modulate these electrophysiological markers, suggesting a preservation or enhancement of processing efficiency through experience-dependent plasticity (Brown et al., 2017; Okhrei et al., 2018; Hajimohammadi & Heidari, 2024).

Within the spectrum of music-based activities, choral singing represents a particularly ecologically valid intervention, integrating auditory–motor coordination, cognitive demands, respiratory training, emotional regulation, and structured social interaction. Community-based and neurophysiological studies report cognitive, emotional, and neural benefits of choir participation in older adults, including enhanced auditory encoding and structural connectivity (Pentikäinen et al., 2021, 2022, 2023; Moisseinen et al., 2024, 2025; Särkämö et al., 2025). Nevertheless, longitudinal studies examining choir participation within a multidomain intervention framework and integrating cognitive, audiometric, and neurophysiological outcomes remain scarce, limiting mechanistic understanding of how choir-based multidomain interventions influence ageing trajectories.

### Aims of the study

The protocol of this study is described in Lippolis et al. (2024). The present feasibility study had two overarching objectives. First, to evaluate the practicality of delivering a choir-centered multidomain intervention in community settings, focusing on recruitment and retention, participant adherence, and the implementation of a comprehensive measurement battery in preparation for a subsequent randomized controlled trial (RCT). Second, to explore the longitudinal effects of the intervention on cognitive and neurophysiological functioning in community-dwelling older adults. The program was designed to integrate musical, cognitive, physical, and social stimulation, consistent with evidence that multidomain approaches provide the strongest preventive benefits in ageing. By embedding choir participation as the central activity, we aimed to determine whether group singing can foster measurable improvements in cognition and auditory–cortical processing compared with both active and passive control conditions.

### Hypothesis

Based on prior evidence linking multidomain interventions and musical engagement to cognitive and neural plasticity, we hypothesized that choir-based training would yield selective longitudinal benefits, outperforming both engagement in alternative non-musical activities and the absence of structured activity. Specifically, we expected improvements in musical abilities, cognitive functioning and concomitant changes in auditory evoked potentials (AEP). Given previous findings of enhanced P2, N2, and P3 responses in musically trained individuals (e.g., Sanju & Kumar, 2016; Brown et al., 2017; Okhrei et al., 2018), we anticipated that choir participation would attenuate age-related slowing in these components (i.e., increased amplitudes and shortened latencies), whereas control groups would show stability or decline.

Moreover, besides absolute peak latencies and amplitudes, we predicted changes in the interpeak measures, as these indices capture the efficiency of information transfer between successive processing stages. Previous research has shown that musical training enhances temporal precision and neural synchrony across auditory–cognitive pathways, reflected in faster or more reliable latencies and reduced temporal variability in auditory evoked responses (Zendel & Alain, 2014; Kraus & Strait, 2015; Liang et al., 2016; Meha-Bettison et al., 2018; Sorati & Behne, 2019). Such evidence makes interpeak dynamics a plausible and sensitive marker of training-induced plasticity.

More broadly, we predicted that embedding choir participation within a multidomain framework would provide synergistic cognitive, motor, and social stimulation, translating into measurable gains in cognitive reserve. At the psychosocial level, we further hypothesized that choir-based participation would be associated with improvements in psychological well-being, as indexed by higher flourishing, and that higher engagement in structured activities would be inversely associated with multidimensional frailty.

In addition to the outcome-related hypothesis, this study was designed as a feasibility investigation to inform a subsequent RCT. Indicators of feasibility included successful recruitment, adherence to choir activities, retention across the 9-months study period and tolerability of the assessment protocol, in line with established recommendations for feasibility studies (e.g., Orsmond & Cohn, 2015; Eldridge et al., 2016). These indicators were evaluated descriptively through participation rates, attrition patterns and completion of the whole assessment.

## Methods

This feasibility study was conducted in preparation for a larger randomized controlled trial (RCT), which will replicate the design with an expanded sample size. The upcoming trial has already been prospectively registered on ClinicalTrials.gov (ID: NCT06767410, registration date: January 9, 2025). Ethical approval was granted by the Ethics Committee of the Department of Education, Psychology, and Communication at the University of Bari (reference number: ET-23-27). To safeguard participants’ identity, each subject was assigned a unique anonymized ID. Given the broad range of assessments required for a comprehensive evaluation, the study was carried out within an interdisciplinary framework, involving collaboration between the Department of Education, Psychology, and Communication, the Department of Pharmacology, and the Otolaryngology and Geriatrics Units of the Polyclinic consortium hospital of Bari.

### Design

The study was conducted in Bari, Southern Italy, and follows a quasi-experimental pre–post design with three arms: a choir-based multi-domain intervention group *“MultiMusic”*, an active control group, and a passive control group (**Figure 1**). Participants were assessed at baseline (T0) and after 9 months (T1). Recruitments and assessments began in November 2023 through local Universities of the Third Age (U3A) and district leisure centers for older adults in both metropolitan and suburban areas of Bari and ended in April 2025. Some participants in the control groups were selected from the waiting list for the choir, rather than being randomly allocated. Although this design may entail baseline imbalances and limits causal inference, it enables the evaluation of the intervention under ecologically valid, community-based conditions. Baseline variables were assessed and statistically controlled to mitigate potential confounding effects.

**Figure 1.**
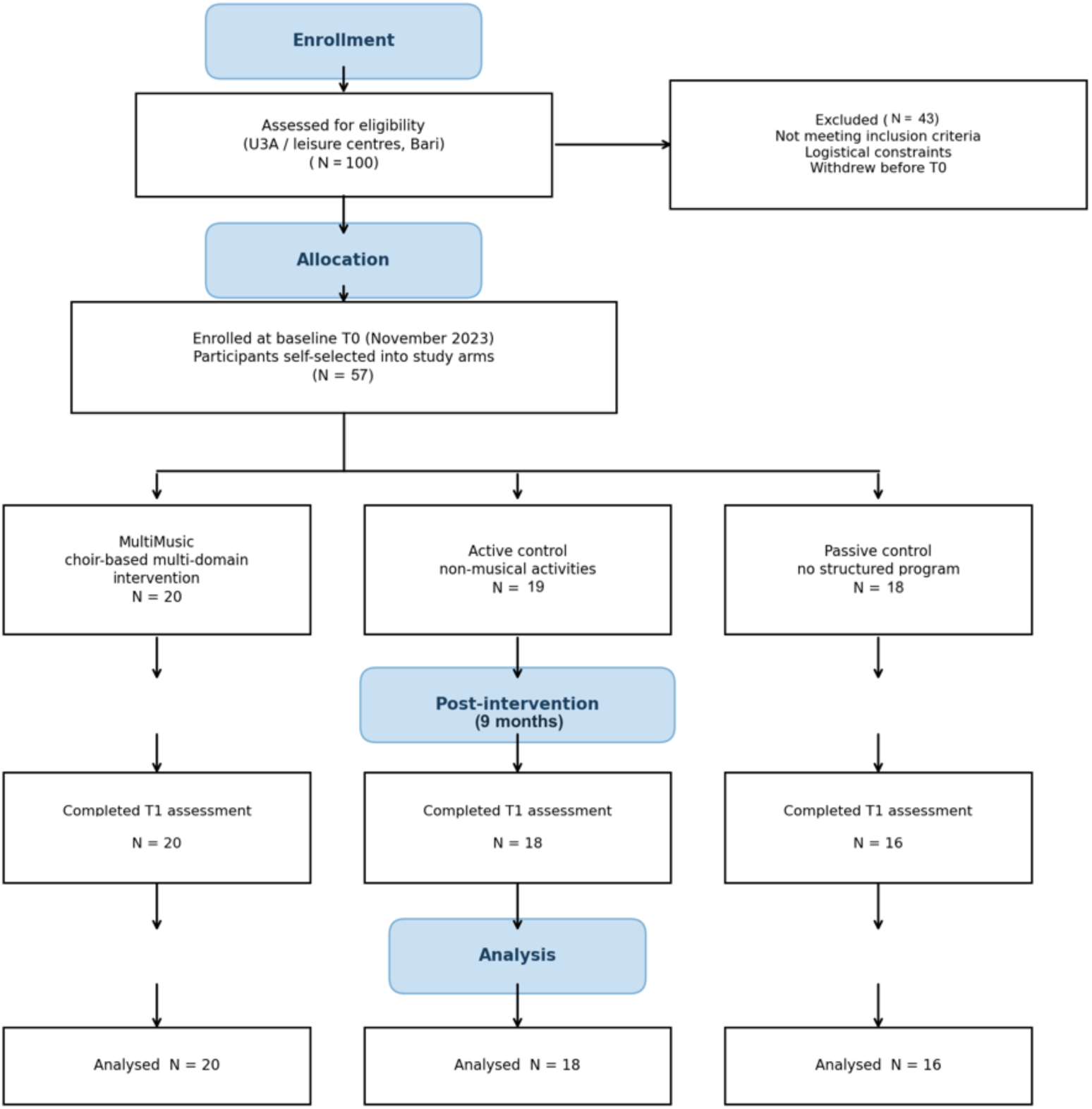
Study design and assessment protocol. As part of this feasibility study, participants self-selected in one of three conditions: a choir-based multi-domain intervention (MultiMusic) group, an active control group (engaged in non-musical activities), or a passive control group (no structured activity). Assessments were conducted at baseline (T0) and after 9 months (T1), covering cognitive, audiometric, and neurophysiological domains.

### Participants

#### Eligibility

Participants were recruited from community-based settings in Bari (Southern Italy) and were eligible for inclusion if they were aged 65 years or older, lived independently in the community, had no extensive prior musical training, and did not present severe cognitive or functional impairments that could interfere with participation in group-based activities or completion of the assessment protocol. Additional eligibility criteria included normal hearing or symmetrical hearing loss, defined as an interaural threshold difference of ≤15 dB at any tested frequency; mild to moderate–severe hearing loss, as determined by the four-frequency pure-tone average (PTA; 500, 1000, 2000, and 4000 Hz) calculated across both ears; and the absence of audiological conditions other than presbycusis.

#### Descriptives

Tables 1a and 1b contain the general sample descriptives. About 100 participants were initially enrolled in the study. However, nearly half of them discontinued participation due to logistical constraints, i.e., they resided in a municipality located farther from the assessment site and could not reliably travel to the hospital-based evaluations. Among those without major logistical barriers, 59 participants remained eligible to continue the study; of these, three withdrew due to health-related reasons and two were excluded from the dataset following the detection of otosclerosis. Hence, the final sample comprised 54 participants (*M*_age_: 72.91; SD = 5.40; 77.7% females) who completed both baseline (T0) and follow-up (T1) assessments, representing those living closer to the assessment center or experiencing fewer logistical barriers to attendance.

**Table 1a.**
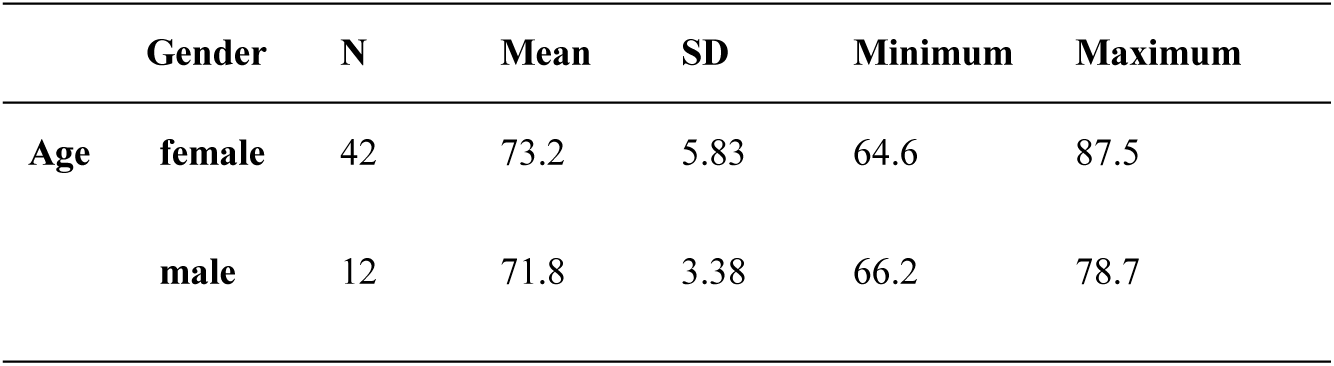
General sample descriptives for age and gender.

**Table 1b.** General sample descriptives for cognitive functioning (MoCa), cognitive reserve (CRI-q) and fluid intelligence (MIQ) at T0.

|  | Mean | SD | Minimum | Maximum |
| --- | --- | --- | --- | --- |
| **MoCa** | 21.74 | 3.70 | 14.00 | 30.00 |
| **CRI-q** | 107.20 | 23.2 | 70.00 | 163.00 |
| **MIQ** | -2.60 | 1.26 | -4.00 | 1.39 |

Participants demonstrated a broad range of cognitive functioning on the MoCA (M = 21.74, SD = 3.70), spanning values typically associated with both normal ageing and mild cognitive impairment. Importantly, Bosco et al. (2017) have shown that Italian older adults generally obtain lower MoCA scores compared to the original validation sample by Nasreddine et al. (2005), with optimal cut-off scores being reduced when distinguishing between normal cognition, probable cognitive impairment, and Alzheimer’s disease. Thus, the mean level observed in our sample should not be interpreted as uniformly pathological, but rather as reflecting the expected variability within an Italian elderly population.

### Intervention

Participants self-selected into three groups. The group participating in the choir-based *MultiMusic* intervention (N = 20; *M*_age_ = 72.1; SD = 4.67) was engaged for two hours a week in choral training starting from November 2023. For the sake of clarity and brevity, this group is hereafter referred simply as Choir. At the beginning of each meeting, the teacher started with body percussion exercises carried out playfully and designed to strengthen body awareness, memory, motor coordination and concentration. Subsequently, respiration exercises and explanations of the functioning of organs impaired in the phonation process were proposed, in order to raise awareness on the use of vocal and respiratory apparatus. This was followed by progressive vocalizations, adapted to the textures of the various vocal sections (basses, tenors, altos, sopranos and treble voices) and, finally, the repertoire is studied, starting with participants’ cultural background, i.e. music from popular culture, then gradually enriched it with more complex, multi-voice and more articulate pieces. Songs were also performed with the participation of soloists and also in the form of duets and trios, in dialogue with the choir, to stimulate creativity and improvisation. Moments of very simple musical and rhythmic reading exercises are also provided, being participants totally lacking musical instruction.

As here we deal with a multidomain intervention, participants were also engaged in various other non-musical activities targeting complementary functional domains. A comprehensive overview of all activities, including their frequency and intensity, is provided in Table 2a, while Table 2b summarizes the distribution and combination of activities across participants. Activities can be grouped into the following macro-categories:

1. Physical activities: this category includes gymnastics, group dancing, sports (e.g., swimming), postural exercise (pilates or body harmony).
2. Musical activities: choir and body percussion.
3. Manual activities: horticulture, handicraft, tailoring, cooking.
4. Arts/Intellectual activities: theater, mental gymnastics, reading, learning programs (such as history, languages etc.).

**Table 2a.**
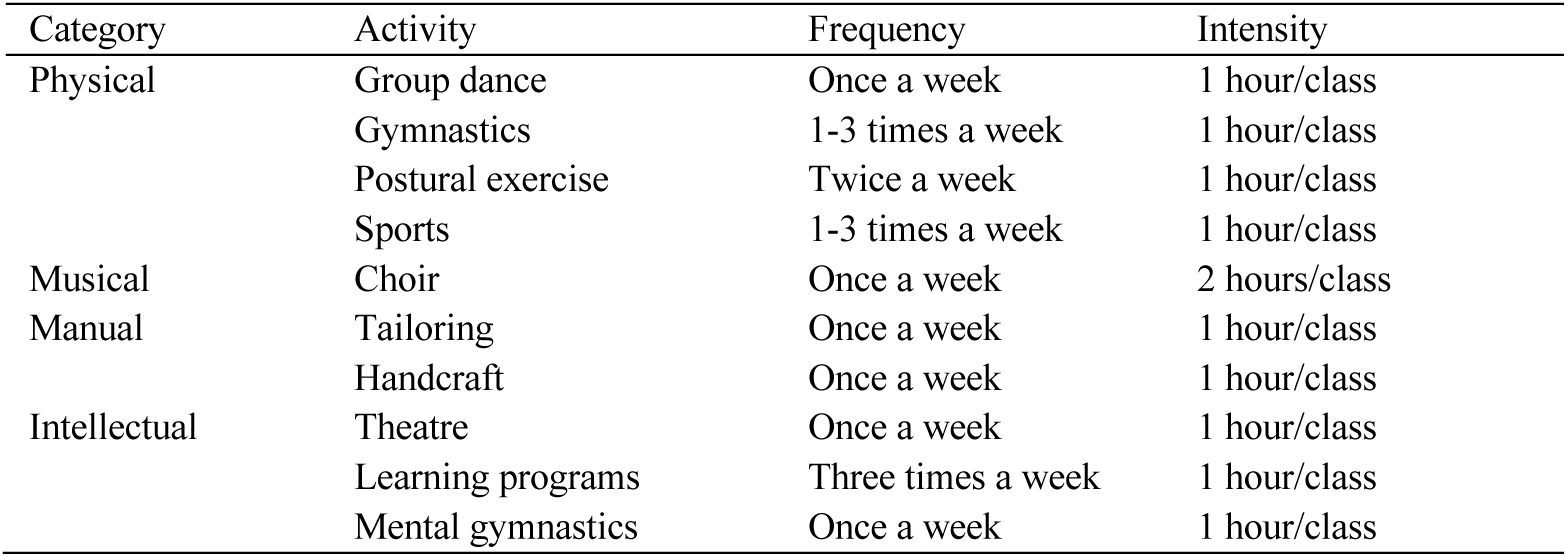
List of activities carried out by the participants.

**Table 2b.** Distribution of activities among participants.

|  | Activities | N° Participants |
| --- | --- | --- |
| Multidomain with choir | Musical activities + Physical activities | 7 |
|  | Musical activities + Intellectual activities | 1 |
|  | Musical activities + Manual activities | 1 |
|  | Musical activities + Physical activities + Intellectual Activities | 4 |
|  | Musical activities + Physical activities + Manual activities | 5 |
|  | Musical activities + Intellectual activities + Manual activities | 0 |
|  | Musical activities + Physical activities + Intellectual activities + Manual activities | 2 |
| Programs without choir | Physical activities | 3 |
|  | Intellectual activities | 10 |
|  | Manual activities | 1 |
|  | Physical activities + Intellectual activities | 3 |
|  | Physical activities + Manual activities | 0 |
|  | Intellectual activities + Manual activities | 0 |
|  | Physical activities + Intellectual activities + Manual activities | 1 |
| Sporadic non-musical activity / No activity | - | 16 |
| Total |  | 54 |

Participants in the active control group (N = 18; *M*_age_ = 74.4; SD = 6.89) engaged in a comparable schedule of non-musical activities, drawn from the same pool of physical, manual, and intellectual programs (e.g., gymnastics, dancing, handicraft, theater etc.), combined according to individual preference and availability. The passive control group (N = 16; *M*_age_ = 71.0; SD = 3.93) did not take part in any structured program.

### Procedures

Participants underwent a comprehensive set of cognitive, musical, audiometric, and neurophysiological assessments. As part of the protocol, participants also provided saliva samples for Brain-Derived Neurotrophic Factor (BDNF) measurement as specified in the registered RCT protocol; ClinicalTrials.gov Identifier: NCT06767410 (for details, see Lippolis et al., 2024). However, these data are not included in the present analyses due to ongoing collection and processing. Data collection took place in district leisure centers seniors and at the Otolaryngology Unit of the Bari University Hospital.

Behavioral and cognitive testing sessions were conducted in the presence of trained researchers and psychologists. Each session lasted approximately 90 minutes per participant and was administered either individually or in small groups (often in pairs). Computerized and paper-based tests were alternated to optimize time and minimize fatigue. Because many participants were not accustomed to technology, researchers provided detailed instructions before computerized tasks, ensuring participants could complete them independently.

Audiometric, neurophysiological, and biological measurements were conducted at the Otolaryngology Unit. Pure-tone audiometry was performed in a sound-attenuated booth, while neurophysiological recordings and BDNF sampling were carried out in dedicated clinical testing rooms. Each session lasted about 40 minutes per participant. Attendance at extra-musical activities and the individual intensity of choir practice were also monitored throughout the intervention using attendance records collected during scheduled activities.

## Materials

### Primary outcomes

#### a) Cognitive tests

Montreal Cognitive Assessment (MoCA; Nasreddine et al., 2005) was chosen as a reliable and common tool to evaluate general cognitive functioning in elderly population, also longitudinally (Cooley et al., 2015; Malek-Ahmadi et al., 2018; Luo et al., 2022; Aiello et al, 2024). MoCa evaluated multiple domains, including attention, task switching, visuoconstructive abilities, naming, memory, WM, calculation, language, fluency, abstraction, and orientation. Subtests included word recall, figure drawing, and object naming.

Matrix Reasoning (MIQ; Condon & Revelle, 2014). The MIQ is a computerized adaptive measure of fluid intelligence, structured as a series of 3×3 geometric patterns with one item missing. Participants must identify the correct piece to complete the matrix, in a task design reminiscent of Raven’s Progressive Matrices. Its adaptive algorithm adjusts task difficulty in real time according to participant performance, thereby enhancing both efficiency and precision. To date, the MIQ has been successfully employed in developmental studies with children, showing strong sensitivity to inter-individual differences in abstract reasoning (Lippolis et al., 2022; Müllensiefen, Elvers, & Frieler, 2022). In the present feasibility study, we extend its application to older adults, aiming to test its validity and responsiveness in the context of age-related changes in fluid intelligence, where adaptive tools may offer advantages in reducing fatigue and measurement error.

#### b) Audiometric Measures

Otoscopy was performed in all participants to examine the external auditory canal and tympanic membrane and to rule out any potential pathological conditions. Pure-tone and speech audiometry were performed using the Madsen® Astera2 audiometer (GN Otometrics), in accordance with European (IEC 60645-1) and international (ISO 389-1) standards, within a sound-treated booth.

### Pure-Tone Audiometry

Air conduction (AC) thresholds were measured using insert earphones at frequencies ranging from 250 to 8000 Hz. Bone conduction (BC) thresholds were assessed using a bone vibrator placed on the mastoid for frequencies between 500 and 4000 Hz. The pure-tone average (PTA) was calculated as the mean threshold at 500, 1000, 2000, and 4000 Hz for each ear, reflecting sensitivity in the speech-relevant frequency range.

### Speech Audiometry

Speech audiometry was conducted following standard clinical procedures. Speech Reception Threshold (SRT) was measured separately for each ear using standardized lists of disyllabic, phonetically balanced words and was defined as the lowest presentation level (dB HL) at which the participant correctly repeated 50% of the presented items and was used as an estimate of the minimum intensity required for reliable speech perception (Hamid and Brookler 2006). Speech intelligibility was assessed using the Word Recognition Score (WRS), defined as the percentage of correctly repeated monosyllabic words presented at suprathreshold intensity levels (Hamid 2006). Standardized monosyllabic, phonetically balanced word lists were presented at a fixed level above each participant’s SRT, typically around 40 dB sensation level. Participants were asked to repeat each word they perceived aloud, and their responses were rated as correct or incorrect to obtain a percentage score. WRS offers an indicator of speech recognition performance beyond thresholds in guaranteed audibility and adds value to threshold-based measures by defining speech comprehension beyond pure-tone sensitivity.

### Speech-in-noise

Matrix Sentence Test (speech-in-noise; Kollmeier & Wesselkamp,1997; Puglisi et al., 2015). It consists of syntactically fixed, but semantically unpredictable five-word sentences generated from a 50-word matrix (10 names, 10 verbs, 10 numerals, 10 adjectives, 10 nouns), e.g., “Andrea eats many useful chairs.The purpose of the Matrix Test is to find the signal/noise ratio (SNR) that allows the patient to understand 50% of the words (SRT, Speech Recognition Threshold). The test uses an adaptive procedure that adjusts the speech level while keeping the background noise fixed, enabling accurate estimation of the SRT with a precision of approximately 1 dB. Normative data in normal-hearing listeners report mean SRT of –6.7 ± 0.7 dB SNR (open-set) and –7.4 ± 0.7 dB SNR (closed-set). Puglisi et al., 2015). In the present study, speech intelligibility in noise was measured using the Oldenburg Measurement Applications software (HörTech GmbH, Oldenburg, Germany). Speech and noise stimuli were presented in free-field conditions through a loudspeaker positioned at 0° azimuth with respect to the listener, with a background noise fixed at 65 dB SPL; All audio stimuli were digitally reproduced from the software’s internal library.

#### c) Neurophysiological Measures

Auditory Brainstem Response (ABR) evaluates hearing function and auditory pathway integrity by recording waves generated along the brainstem. Wave I originates from the auditory nerve and represents spiral ganglion cell activity; Wave II is generated by globular cells of the cochlear nucleus; Wave III reflects activity from spherical cells of the cochlear nucleus; and Waves IV and V arise from central brainstem structures, including the lateral lemniscus, inferior colliculus, and medial superior olive (Starr & Hamilton, 1976; Melcher & Kiang, 1996).

ABR was recorded using the Interacoustics Eclipse EP25 system. Four disposable surface electrodes were applied: reference electrodes were placed on the right (M1) and left (M2) mastoids, the active electrode on the upper forehead (Fz), and the ground electrode on the lower forehead, slightly right of the midline (Gnd). Acoustic stimuli were delivered monaurally through earphones (Ear Tone ABR). A click stimulus at 90 dB nHL was presented at a rate of 27.7 clicks per second. The recording window was set to 10 milliseconds. Each participant underwent two trials per ear, with an average of 2,000 responses recorded for each. High-pass and low-pass filters were set at 100 Hz and 3,000 Hz, respectively. Recordings were conducted while participants lay supine on a stretcher and were instructed to relax and remain as still as possible. The absolute latencies of waves I, III, and V were measured, as well as the interpeak latencies I–III, I–V, and III–V.

P300 Auditory evoked potential (AEP): P300 AEPwere recorded monaurally using the same electrodes montage and for ABR, with the exception that the active electrode was moved to the vertex (Cz), to optimize cortical potential detection. Interelectrode impedances were maintained below 5 kΩ. The ER-3A insert earphones were used to provide auditory stimuli. An oddball paradigm was employed, using a 1KHz tone burst as standard stimulus and 2KHz tone burst as target stimulus, with a 20% target probability and 80% standard probability. During the test, each stimulus window lasted 700 milliseconds and from 1 Hz to 20Hz. Participants were instructed to listen to a series of identical auditory stimuli, interspersed randomly with a rare auditory stimulus, and to maintain a mental count of how many rare stimuli they had heard. Participants maintained a supine position throughout the procedure, remaining awake with their eyes open and were instructed to minimize movement, blinking, and swallowing.

Oddball-elicited AEP were examined focusing on the P2, N2, and P3 components. The P2 component (150–220 ms) was defined as an early positive deflection indexing sensory–attentional encoding. The N2 component (200–350 ms) was identified as a negative peak associated with novelty detection and cognitive control, whereas the P3 component (300–600 ms) was characterized as a large positive deflection reflecting attentional resource allocation and context updating.

For each component, absolute peak latencies (in milliseconds) were extracted as the time from stimulus onset to the maximum or minimum peak. Component amplitudes (in microvolts) were quantified relative to baseline or, when appropriate, to the preceding trough. In addition to component-specific measures, inter-peak temporal intervals between successive components (e.g., P2–N2 and N2–P3) were calculated to index processing speed and the efficiency of cortical information transmission across successive stages of auditory–cognitive processing. Inter-peak latencies (IPLs) were considered alongside absolute peak latencies to capture not only isolated responses but also their temporal relationships. The P300 was prioritized in the analyses given its established sensitivity to cognitive ageing, attentional modulation, and auditory training effects, while ABR served as a physiological control.

#### d) Multidimensional frailty

Multidimensional frailty was assessed using the Selfy-MPI (Pilotto et al., 2019) via the Portable-MPI app, a digital self-administered version of the Multidimensional Prognostic Index designed to capture frailty as a cumulative vulnerability across multiple domains. The instrument integrates information on functional autonomy in activities of daily living and instrumental activities of daily living (ADLs/IADLs), mobility, cognitive functioning, nutritional status, comorbidity burden, medication use/polypharmacy, and living situation/social context. Responses across domains are combined to compute a single composite frailty score, providing an overall index of multidimensional frailty, with higher scores indicating greater vulnerability.

#### e) Socio-psychological well-being

The Flourishing Scale (Diener et al., 2010) is an 8-item self-report questionnaire designed to capture key aspects of eudaimonic well-being. The scale evaluates multiple domains of positive psychological functioning, including purpose and meaning in life, quality of interpersonal relationships, self-esteem, competence, optimism, and engagement with life. Participants rated their agreement with each statement (e.g., *“I lead a purposeful and meaningful life”*) on a 7-point Likert scale ranging from 1 (*strongly disagree*) to 7 (*strongly agree*). Item scores were summed to yield a total flourishing score ranging from 8 to 56, with higher values indicating greater psychological flourishing and overall well-being. In the present study, the total score was used as a continuous measure to assess longitudinal changes in psychological well-being across groups.

### Secondary outcomes

#### a) Musical abilities

The following LongGold computerized, adaptive perceptual tasks were employed for this study:

Melody Discrimination Test (MDT). The MDT is an adaptive three-alternative forced-choice (3-AFC) paradigm designed to measure melodic discrimination (Harrison et al., 2017). On each trial, participants hear three versions of the same melody: two are identical in interval structure (lures), while one contains a single altered note (the “odd” version). The task is to identify the odd melody by comparing the three versions and selecting the one that does not belong to the most similar pair. Task difficulty is manipulated by varying the length of the melodies, thereby increasing cognitive load. In the present study, the MDT included 18 items.

Mistuning Perception Test (MPT). The MPT follows a two-alternative forced-choice (2-AFC) format where each trial presents two renditions of the same pop-music excerpt, one in tune and the other deliberately pitch-shifted (Larrouy-Maestri et al., 2019). The stimuli consist of short segments (6–12 s) with a vocal line accompanied by instrumentation. The “out-of-tune” versions are produced by applying a constant pitch shift to the voice, while the accompaniment remains unaltered. Participants are required to decide which of the two excerpts contains the mistuned vocal line.

Rhythm Ability Test (RAT). The RAT is an adaptive assessment of rhythmic perception and reproduction (MacGregor et al., 2022). Participants listen to rhythmic sequences consisting of high- and low-pitched tones and must select the image that best represents the heard pattern. Visual representations display rows of colored squares, with high tones at the top row and low tones at the bottom. The test increases in complexity across trials, with sequences expanding from 4 to 16 elements (quarter, eighth, and sixteenth notes).

Emotion Discrimination Test (EDT). The EDT is an adaptive task using a two-alternative forced-choice (2-AFC) format to evaluate recognition of musical emotions (MacGregor & Müllensiefen, 2019). For each trial, two variations of the same melody are presented, and participants must choose which one conveys a target emotion (anger, happiness, sadness, or tenderness). Item difficulty varies according to differences in acoustic and structural features across excerpts. For example, anger is typically conveyed by loud amplitude, fast tempo, and high roughness; happiness by fast tempo and high amplitude but lower roughness; sadness by slow tempo and low intensity; and tenderness by acoustic qualities similar to sadness but with subtle expressive cues.

#### b) Physical activity

Physical activity levels were assessed using the Global Physical Activity Questionnaire (GPAQ), developed by Armstrong and Bull (2006) for the World Health Organization (WHO). The GPAQ is a standardized self-report instrument designed to capture habitual physical activity across multiple domains of daily life, allowing a comprehensive characterization of overall activity patterns in adult and older populations. Specifically, the questionnaire assesses physical activity performed in work-related contexts, transport-related activity (i.e., walking or cycling to and from places), and recreational physical activity, as well as sedentary behavior.

### Other measures

Cognitive Reserve Index Questionnaire (CRI-q; Nucci, Mapelli & Mondini, 2012). The CRI-q is one of the most used tools for quantifying cognitive reserve (Nogueira et al., 2022), defined as the reserve of mental resources accrued throughout life. It collects information across three domains, i.e., education, occupational attainment, and social/intellectual engagement, to generate a composite index. The instrument has been applied in numerous ageing studies across populations, including Italian population, to examine protective factors against cognitive decline, showing valid associations with performance on memory, attention, and executive tasks (Caffo et al., 2016; Maiovis et al., 2016; Ozakbas et al., 2021; Cao et al., 2022; Marselli et al., 2024).

Although cognitive reserve was initially listed as a primary outcome in the trial registration, in the final analytic plan it was treated as a baseline control variable rather than a longitudinal outcome. This decision was made on theoretical grounds, because CRI-q reflects accumulated lifetime cognitive reserve rather than a short-term intervention-sensitive endpoint.

### Statistical Analysis

All analyses were conducted in R (version 4.3., R Core Team, 2025) using the packages *lme4* 1.1-35, *lmerTest* 3.1-3, *emmeans* 1.9-5, *cocor* 1.4-2, *car* 3.1-2, and *performance* 0.10-4. Baseline normality and variance homogeneity were assessed with Shapiro–Wilk and Levene’s tests. Full output and results can be found in the Supplementary Information.

#### Baseline analyses

A cluster analysis of baseline variables was first conducted to inform domain definition. Specifically, a hierarchical cluster analysis was performed using a correlation-based dissimilarity measure. The distance between subjects was defined as 1−∣r∣, where r is the Pearson correlation coefficient computed across the set of variables. This distance metric groups subjects who exhibit similar profiles across variables, regardless of the direction of the correlation. Complete linkage was used as the agglomeration method, and only subjects with complete data on all clustering variables were included (listwise deletion; n = 95).

Audiometric measures clustered with perfect purity (7/7 variables), confirming their coherence as a single domain. Neurophysiological measures formed three clusters, separating absolute peak latencies (P2, N2, P3 bilaterally), inter-peak latencies (IPLs; P2–N2 and N2–P3), and inter-peak amplitudes (IPAs; P2–N2 and N2–P3), while cognitive and musical tasks clustered together, reflecting their strong empirical association.

Based on this structure and theoretical considerations, four operational domains were defined for subsequent analyses: (i) cognitive (MoCA, MIQ, CRI-q), (ii) musical (MDT, MPT, EDT, RAT), (iii) audiometric (pure-tone and speech thresholds, intelligibility, and speech reception in noise), and (iv) neurophysiological, subdivided into latency (absolute latencies and IPLs) and IPAs. Although cognitive and musical variables clustered together, they were analyzed separately given their distinct conceptual targets. For neurophysiology, a parsimonious latency–amplitude subdivision was adopted to avoid peak-specific fragmentation driven by methodological covariance rather than functional distinctions.

For conceptual and statistical coherence, Selfy-MPI and Flourishing scores were excluded from the cluster-based domain structure. As global, integrative outcome constructs indexing multidimensional frailty and psychological well-being, respectively, they were treated as downstream outcomes and analyzed separately using targeted longitudinal models consistent with the multidomain framework.

Baseline group differences across domains were assessed using one-way ANOVAs (Choir, Active Control, Passive Control). Pairwise baseline and T1 correlations within and across domains were computed using Pearson’s or Spearman’s coefficients, as appropriate. Audiometric scores were directionally aligned so that higher values consistently reflected better performance prior to computing composites (PTA and SRT scores multiplied by −1; intelligibility unchanged). For bilateral audiometric and neurophysiological measures, differences between correlation coefficients were tested using Zou’s method (2007). Multiple comparisons were controlled within and across domains using the Benjamini–Hochberg FDR procedure (α = 0.05).

#### Longitudinal analyses

Longitudinal effects were examined using hierarchical linear mixed-effects models (maximum likelihood estimation), with model building guided by step-up comparisons based on likelihood-ratio tests and information criteria (AIC, BIC). Dependent variables included domain-specific outcomes: cognitive outcomes (MoCA and MIQ), musical tasks (MDT, MPT, EDT, RAT), audiometric measures (SRT, PTA, intelligibility, Matrix Test), neurophysiological AEP measures (P3 latency and amplitude), multidimensional frailty (Selfy-MPI), and psychological well-being (Flourishing Scale).

Domain-specific baseline covariates were derived at T0 using principal component analysis (PCA) conducted separately within each domain. For the cognitive domain, the PCA included MoCA, MIQ, and CRI-q, with CRI-q contributing only to the derivation of the baseline cognitive component and not treated as a longitudinal outcome. Variables were z-standardized, PCA was fitted on complete cases, and component scores were computed for all participants, with projection used in cases of partial missingness. Leading PCs were retained based on explained variance and interpretability and entered as baseline covariates. The domain-level PC scores were derived independently from the hierarchical cluster analysis described above: whereas clustering operated on the original observed variables to identify subject subgroups, PCA operated on domain-specific variable sets to reduce collinearity and produce summary scores.

Models included fixed effects of Time, Group, and, when applicable, Side (right/left), with participant-specific random intercepts. Activity intensity and baseline ability were included as covariates to control for individual differences and additional training exposure. Fixed effects and interaction terms (Time × Group, Time × Side, Group × Side, and Time × Group × Side) were entered hierarchically and retained only when supported by a significant likelihood-ratio test (p < .05) and a ≥2-point reduction in both AIC and BIC. When significant Time × Group interactions were observed, baseline outcome values were additionally included to adjust for initial between-group differences.

For Selfy-MPI and Flourishing Scale an additional mixed model was set up. Besides testing Group as the main between-subject factor, the activity level (i.e., number of structured activities) was tested in a parallel model to further assess the role of engagement. For descriptive purposes, participants were categorized into four activity-count groups (0, 1, 2, ≥3 activities). For inferential analyses, activity engagement was dichotomized into Low (0–1 activities) and High (≥2 activities), reflecting a distinction between minimal and cumulative multidomain engagement. Estimated marginal means and post-hoc contrasts were obtained using Kenward–Roger degrees of freedom, with Bonferroni correction for multiple comparisons.

## Results

### Baseline results

At T0, Levene’s test of homogeneity of variance showed no significant heteroscedasticity for the measures tested. Seven measures survived the Benjamini–Hozchberg correction applied to the one-way ANOVA. The Choir scored lower than both control groups on CRI-q (pFDR < .001), MoCA (pFDR = .019), and MIQ (pFDR = .029), indicating poorer cognitive reserve and global cognition. In the musical domain, the Choir showed lower rhythmic aptitude on RAT (pFDR = .018). Electrophysiologically, the Choir showed longer left-hemisphere P2 latency (pFDR < .05; higher latency = worse) and higher N2–P3 IPL values bilaterally (right: pFDR < .001; left: pFDR < .01). No audiometric measures survived FDR correction (Figure 2).

**Figure 2.**
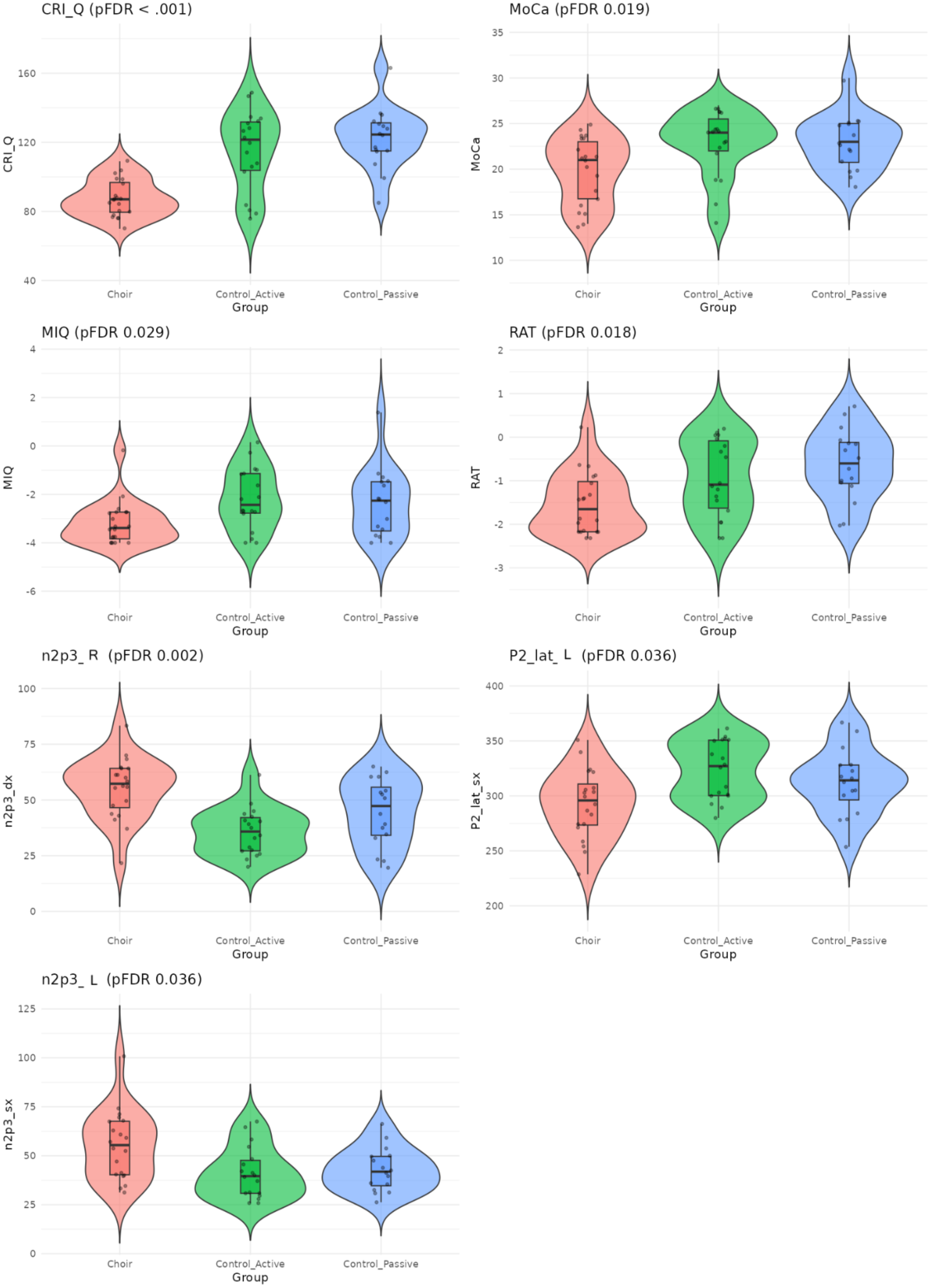
Baseline group comparisons (T0) on cognitive, musical, and neurophysiological measures that survived Benjamini–Hochberg FDR correction. Violin plots display the distribution, median, and interquartile range for each group (Choir, Active Control, Passive Control). These results highlight baseline disadvantages in the Choir group across cognitive reserve, global cognition, musical rhythm aptitude, and specific electrophysiological latency indices.

Descriptive statistics indicated no significant baseline differences between groups in Selfy-MPI scores (Kruskal–Wallis test, *p* = 0.072), confirming comparable levels of multidimensional frailty at study entry. Flourishing Scale’s baseline scores did not differ significantly among the three groups (Kruskal–Wallis test, *p* = 0.407).

### Correlation results

Shapiro–Wilk tests indicated non-normality for multiple variables, so we used Spearman correlations whenever at least one variable in a pair violated normality (p < .05). Widespread correlations were observed among tests within the same domains. Regarding inter-domain correlations, musical tests related to audiometric measures primarily via MDT, with significant positive links to PTA left, PTA right, SRT left, and SRT right. A positive trend just above the FDR threshold was also observed for Matrix, as well as for Intelligibility thresholds of both sides. EDT and RAT did not yield FDR-significant associations with any of the audiometric measures.

Significant correlations also emerged between cognitive and musical measures, including CRI-q with RAT, MoCA with MDT, MIQ with RAT, MoCA with MPT, MoCA with RAT and MIQ with MDT. In contrast, associations between cognitive and audiometric measures showed positive trends but did not remain significant after FDR correction in this analysis. MIQ correlated with the right N2 neurophysiological amplitude index, while several neurophysiological amplitude measures were associated with neurophysiological latency indices. As for the Zou’s test, the largest asymmetry was observed for N2 amplitude with MIQ, where the right-side correlation exceeded the left and the Zou 95% CI excluded zero (significant), whereas other sizeable differences did not reach significance.

At T1, intra-domain coherence was largely preserved relative to T0, with audiometric variables remaining the most strongly intercorrelated (rs = 0.49–0.96) and cognitive and musical domains showing moderate internal associations. However, the inter-domain pattern shifted notably: the musical–audiometric links observed at T0 (MDT with PTA and SRT) did not survive FDR correction at T1, nor did the cognitive–musical associations (e.g., MoCA/MIQ with MDT/RAT). Instead, MIQ emerged as a new cross-domain bridge between cognitive and audiometric measures (rs = 0.40–0.45). Neurophysiological amplitude and IPL sub-domains remained internally correlated, but again showed no significant cross-domain links after FDR correction. Unlike T0, no significant bilateral asymmetries were detected at T1 (Zou’s method, all p > 0.40).

Flourishing scores showed no significant correlations with cognitive, musical, audiometric, or neurophysiological measures after within-domain FDR correction at T0 nor T1. While at baseline correlations between outcome measures did not survive correction, at T1, higher frailty was associated with lower cognitive functioning (MoCA, MIQ, CRI-q) and with slower auditory–cortical processing indexed by P2–N2 inter-peak latency.

### Longitudinal results - Groups

Significant longitudinal effects were found only for the Neurophysiological (latency) domains. Longitudinal trends were also found for MIQ, Selfy-MPI and Flourishing Scale.

The linear mixed-effects model revealed a significant Time × Group interaction for the N2–P3 inter-peak latency (F(2, 377.56) = 33.42, p < .001). Post hoc comparisons (Kenward–Roger correction, Bonferroni-adjusted) showed that only the choir group exhibited a significant reduction from T0 to T1 (−21.12 ms, SE = 1.84, t = −11.50, p < .001). In contrast, no significant longitudinal change was observed in the active control group (−0.92 ms, SE = 1.98, p = 1.000) or in the passive control group (−3.92 ms, SE = 2.05, p = .172). At baseline (T0), groups did not significantly differ. At T1, however, the choir group showed significantly shorter N2–P3 inter-peak latencies compared to both the active (−14.72 ms, p < .001) and passive control groups (−13.43 ms, p < .001), while the two control groups did not differ from each other (p = 1.000) (Figure 3a, 3b).

**Figure 3.**
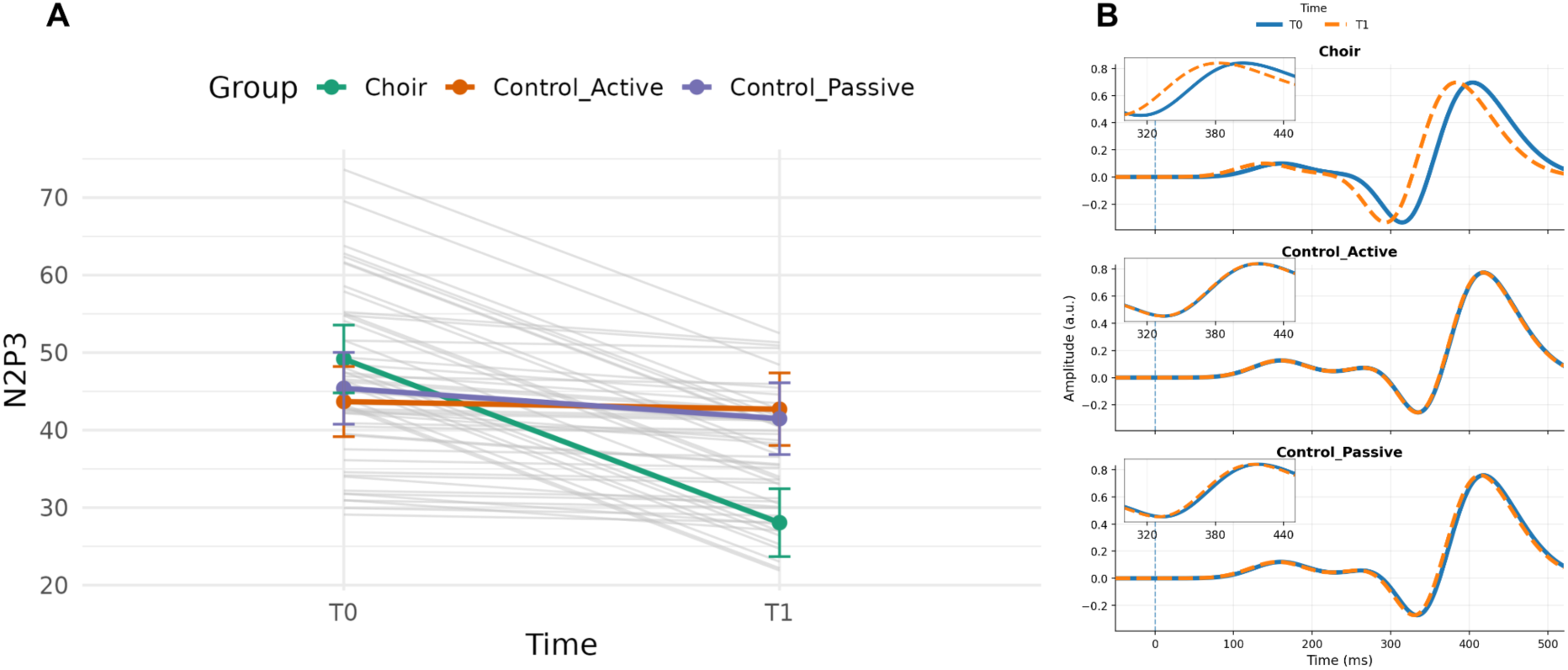
Longitudinal changes in N2–P3 inter-peak latency (IPL). (A) Spaghetti plot of individual fitted values derived from the linear mixed-effects model. Thin grey lines represent subject-specific trajectories, while colored points and lines indicate estimated marginal means (± SE) at T0 and T1. A significant reduction in N2–P3 IPL was observed only in the Choir group (p < .001), with no longitudinal change in the active or passive control groups. (B) Group-averaged auditory evoked potentials (AEPs) illustrating the N2–P3 IPL by group and hemisphere. The shorter IPL at T1 reflects a faster N2→P3 transition in the Choir group.

For P2 latency, the linear mixed-effects model revealed no significant effects of Time or Group, and no evidence of differential longitudinal changes across groups (all ps > .54). For N2 latency, a significant Time × Group interaction emerged (F(2, 159) = 3.84, p = .023). However, post hoc within-group comparisons did not show significant longitudinal changes in any group (all ps ≥ .066). No significant interactions were found for P3 latency and P2-N2 IPL.

For MIQ, the interaction Time x Group was significant, F(2, 108) = 4.06, p = .020. Post hoc comparisons indicated a marginal improvement in the Choir group (Δ = +0.42, p = .056), whereas no significant changes were observed in the control groups. No significant interaction effects were found for MoCa.

In the mixed-effects models examining longitudinal changes in Selfy-MPI and Flourishing, the Time × Group interaction was not significant, indicating that overall changes across groups did not differ statistically. However, exploratory within-group contrasts were conducted to characterize longitudinal trends. These contrasts showed a modest reduction in frailty in the Choir group (Δ = −0.036, p = .049), whereas no significant changes were observed in the Active (p = .682) or Passive control groups (p = .124) (Figure 4). For Flourishing Scale, pairwise comparisons showed between-group differences at T1, where the difference between the Choir and Passive Control groups approached significance (Δ = 3.56, p = .054). Contrasts are summarized in Table 3.

**Figure 4.**
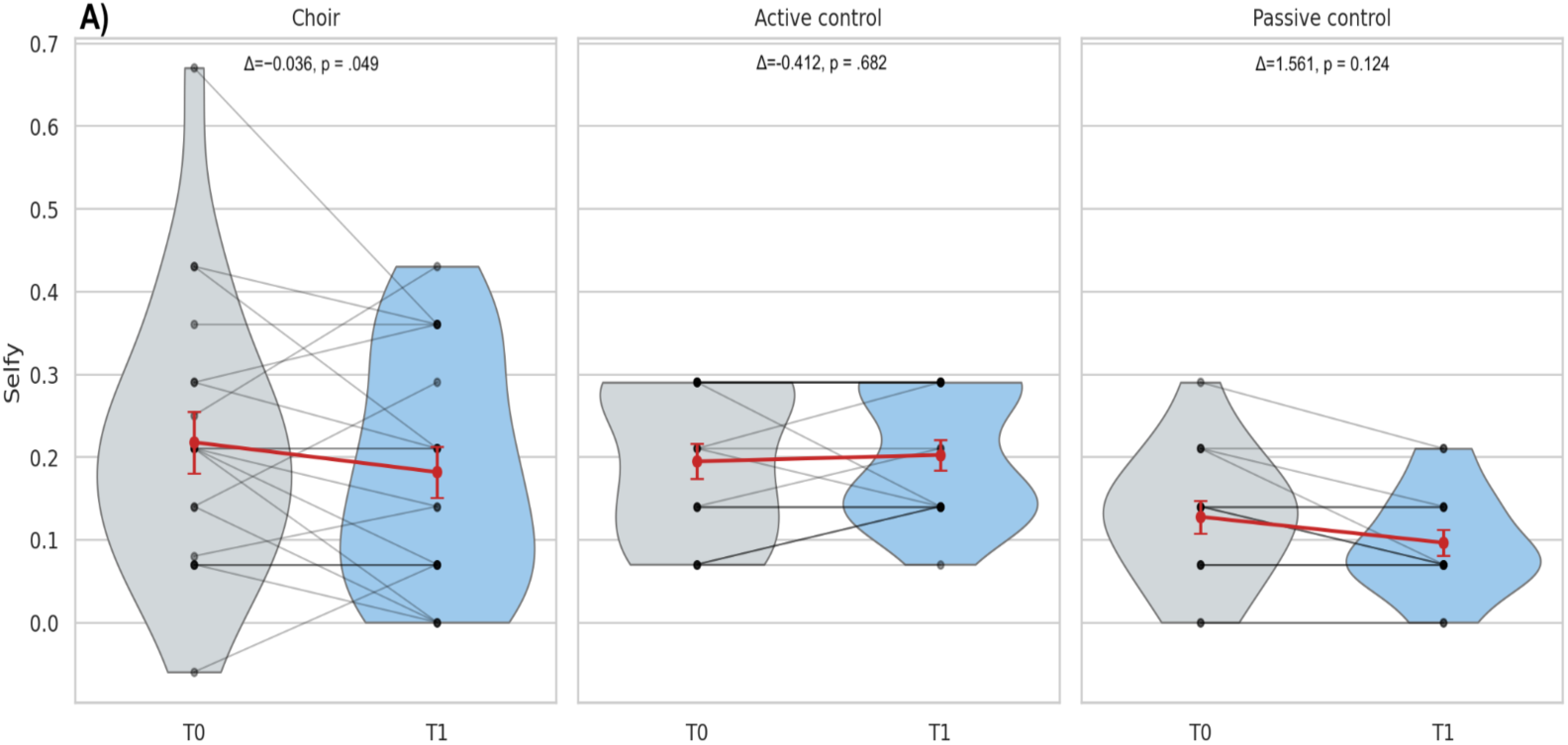
Multidimensional frailty (Selfy-MPI) at baseline and follow-up by study group. Violin plots show the distribution of Selfy-MPI scores at T0 and T1 in the Choir, Active control, and Passive control groups. Individual participant trajectories are overlaid as connecting lines between repeated measurements, and red points with error bars represent group means ± SE. Exploratory within-group contrasts showed a significant reduction in frailty in the Choir group (Δ=−0.036, p = .049), with no significant change in the Active control (p = .682) or Passive control groups (p = .124).

**Table 3.** Post-hoc Group contrasts N2-P3 IPL, MIQ, Selfy-MPI, and Flourishing.

| Outcome | Contrast type | Comparison | $\Delta$ | SE | df | t | p |
| --- | --- | --- | --- | --- | --- | --- | --- |
| N2-P3 IPL | Within-group | Choir, T1 - T0 | -21.12 | 1.84 | 377 | -11.50 | < .001 |
| N2-P3 IPL | Within-group | Control_Active, T1-T0 | -0.918 | 1.98 | 387 | 0.465 | 1.0000 |
| N2-P3 IPL | Within-group | Control_Passive, T1-T0 | -3.917 | 2.05 | 377 | -1.907 | 0.1717 |
| MIQ | Within-group | Choir, T1 - T0 | 0.424 | 0.217 | 58.3 | 1.952 | .056 |
| MIQ | Within-group | Control_Active, T1-T0 | -0.377 | 0.229 | 58.3 | -1.644 | 0.1056 |
| MIQ | Within-group | Control_Passive, T1-T0 | 0.357 | 0.243 | 58.3 | 1.470 | 0.1468 |
| Selfy-MPI | Within-group | Choir, T1 - T0 | -0.036 | 0.0179 | 58.3 | 2.011 | .049* |
| Selfy-MPI | Within-group | Control_Active, T1 - T0 | -0.0077 | 0.0189 | 58.3 | -0.412 | 0.6817 |
| Selfy-MPI | Within-group | Control_Passive, T1 - T0 | 0.03125 | 0.0200 | 58.3 | 1.561 | 0.1239 |

Note. $\Delta$ values represent estimated post hoc contrasts from linear mixed-effects models. Within-group change scores are expressed as T1 - T0 where applicable. Positive values for between-group differences indicate higher scores in the first group listed. Bonferroni-adjusted p values are reported where applicable. Asterisks indicate significance level: \*\*\*: $p < 0.001$ ; \*\*: $0.001 \leq p < 0.01$ ; \*: $0.01 \leq p < 0.05$ ; .: $0.05 \leq p < 0.1$ ; (no symbol): $p \geq 0.1$ .

### Longitudinal results – Level of engagement

The analyses based on activity engagement indicated that a significant reduction in frailty was observed only among participants with high levels of activity engagement (Δ = −0.036, p = .034), whereas no change emerged in the low-engagement group (p = .641) (Figure 5a).

**Figure 5a.**
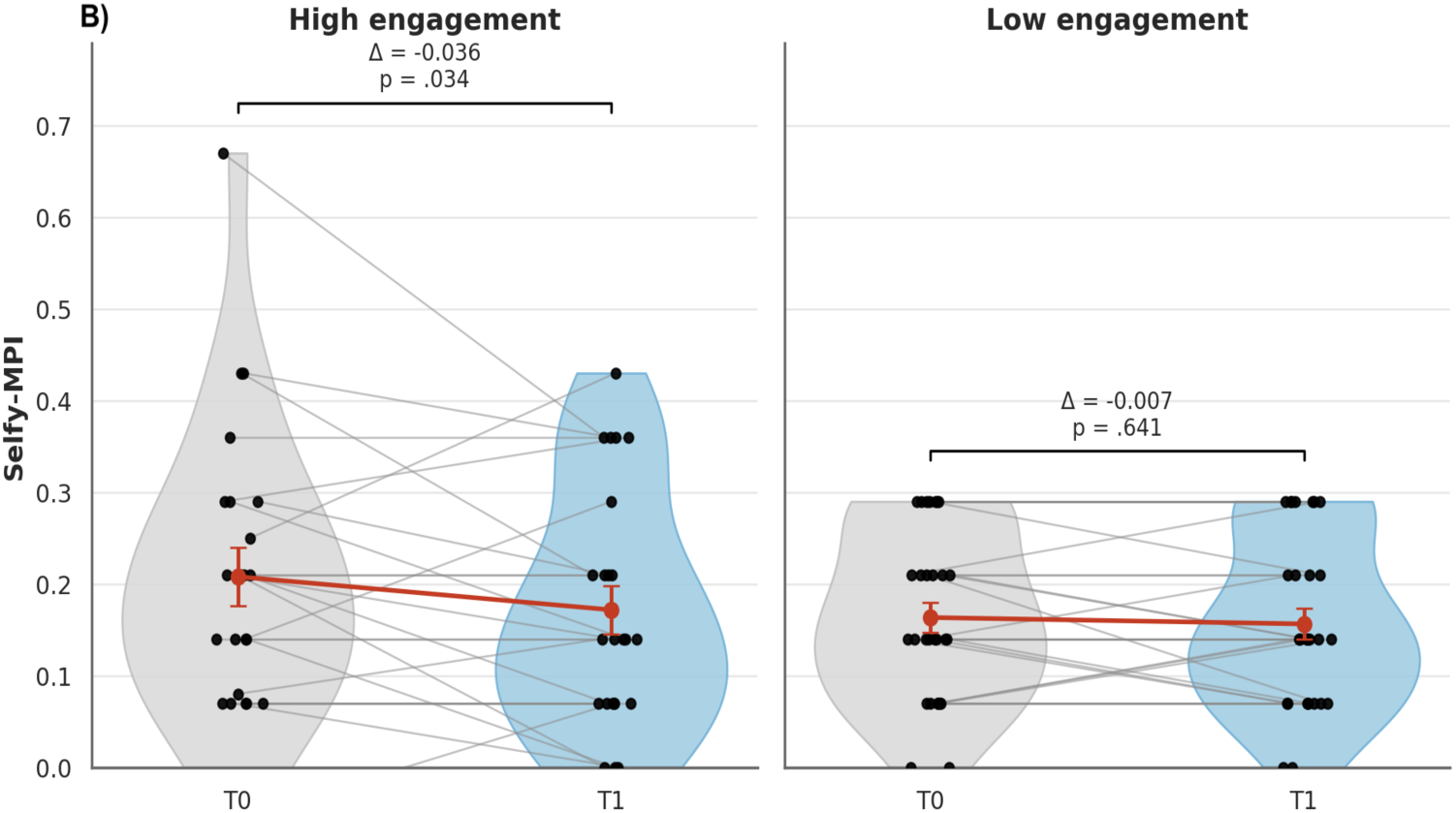
Selfy-MPI scores at baseline (T0) and post-intervention (T1) by activity engagement level. Violins depict score distributions, black dots indicate individual participants, grey lines connect repeated measures, and red points with error bars represent estimated means ± standard error. Post hoc comparisons showed a significant decrease in the High engagement group, whereas no significant change emerged in the Low engagement group.

Between-group comparisons at T1 revealed significantly higher flourishing scores in the high engagement group compared to the low engagement group (Δ = 3.16, p = .008) (Figure 5b). Contrasts are summarized in Table 4.

**Figure 5b.**
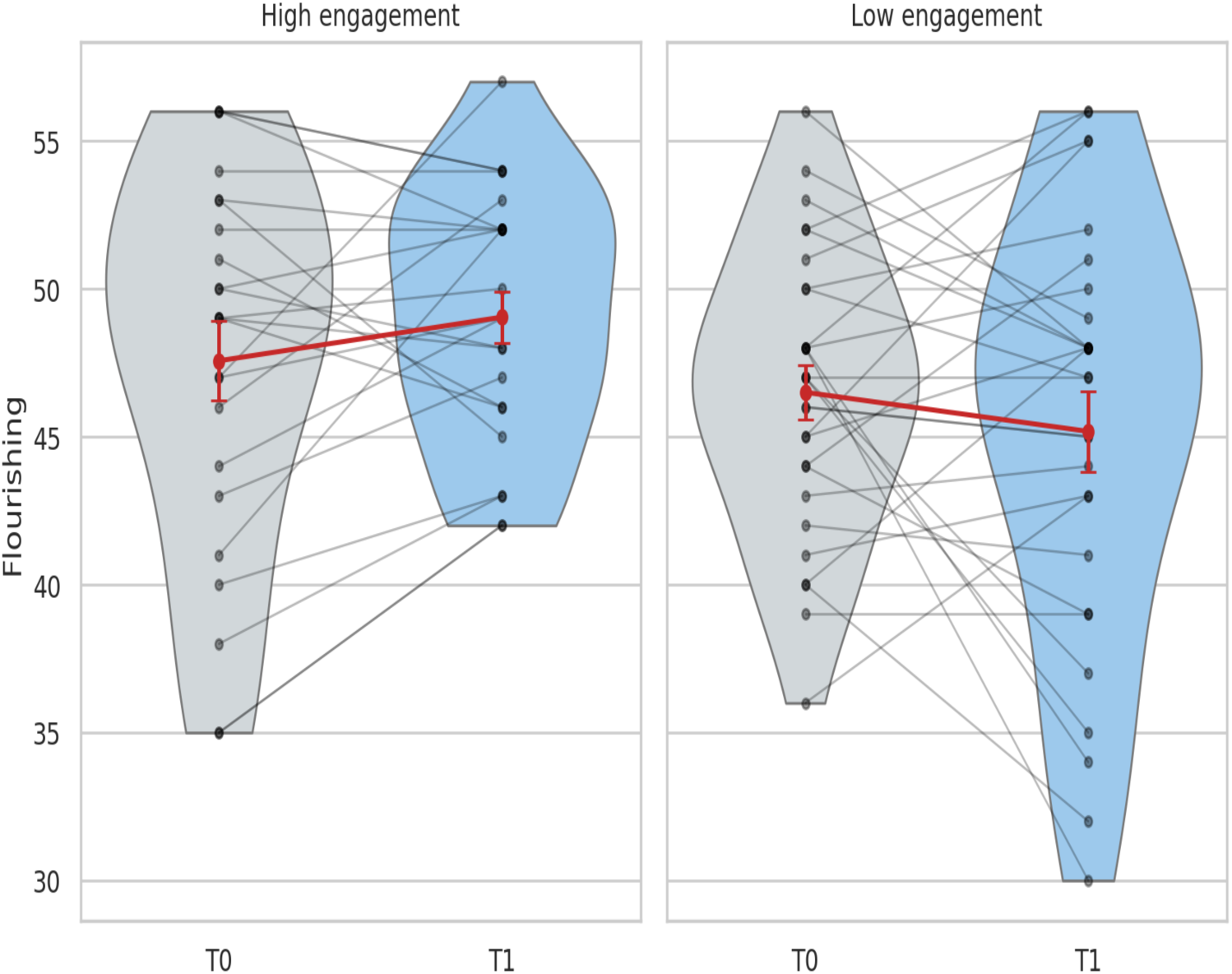
Psychological well-being (Flourishing Scale) at baseline and follow-up by activity engagement level. Violin plots show the distribution of Flourishing Scale scores at T0 and T1 in participants with high versus low activity engagement. Individual participant trajectories are overlaid as connecting lines between repeated measurements, and red points with error bars represent group means ± SE. participants with high activity engagement showed significantly higher flourishing scores than those with low engagement at T1 (Δ=3.16, p = .008), whereas no difference was observed at baseline (p = .814).

**Table 4.** Post-hoc Engagement contrasts N2-P3 IPL, MIQ, Selfy-MPI, and Flourishing.

| Outcome | Contrast type | Comparison | $\Delta$ | SE | df | t | p |
| --- | --- | --- | --- | --- | --- | --- | --- |
| Selfy-MPI | Within-group | High engagement, T1 - T0 | -0.0362 | 0.0167 | 57.2 | -2.169 | .034* |
| Selfy-MPI | Within-group | Low engagement, T1 - T0 | -0.0070 | 0.0149 | 57.2 | -0.468 | .641 |
| Flourishing Scale | Between-group | High - Low engagement T0 | 0.274 | 1.16 | 107 | 0.236 | 0.8142 |
| Flourishing Scale | Between-group | High - Low engagement T1 | 3.160 | 1.16 | 107 | 2.720 | .008** |

Note. $\Delta$ values represent estimated post hoc contrasts from linear mixed-effects models. Within-group and between-group change scores are expressed as T1 - T0 where applicable. Positive values for between-group differences indicate higher scores in the first group listed. Bonferroni-adjusted p values are reported where applicable. Asterisks indicate significance level: \*\*\*, $p < 0.001$ ; \*\*, $0.001 \leq p < 0.01$ ; \*, $0.01 \leq p < 0.05$ ; ., $0.05 \leq p < 0.1$ ; (no symbol): $p \geq 0.1$ .

## Discussion

This feasibility study tested a choir-centered, community-delivered multidomain program (“MultiMusic”) of 9 months in healthy older adults and examined longitudinal change in cognition, multi-dimensional frailty, musical ability, psychological well-being, audiometric measures and auditory–cortical evoked potentials (AEP). The strongest and most consistent longitudinal change was found in N2-P3 interpeak latency (IPL) only for the Choir group.

### Feasibility Outcomes

In line with the predefined feasibility objectives, the present study provides encouraging evidence regarding the feasibility of implementing a choir-based multidomain intervention in community-dwelling older adults. Recruitment was successful, indicating substantial interest and willingness to engage in structured musical and leisure activities within this population. Adherence to choir participation was high among those enrolled, supporting the acceptability of choir singing as a sustained group-based activity over an extended period. Importantly, retention across the 9-month study period was strongly influenced by logistical rather than motivational factors: participants residing closer to the assessment facilities showed high completion rates, whereas attrition primarily reflected difficulties associated with travel to hospital-based assessments.

This pattern suggests that dropout was not driven by lack of engagement with the intervention itself, but rather by structural barriers. Finally, the tolerability of the cognitive, audiometric, and neurophysiological assessments was generally good among participants who completed the post session testing, supporting the practicality of the multimodal assessment battery. Together, these findings indicate that the core elements of the intervention (i.e., recruitment, adherence, and acceptability) met feasibility benchmarks, while also highlighting specific logistical constraints that should be addressed in a future randomized controlled trial (RCT).

### Baseline characteristics and correlations

Baseline analyses indicated that choir participants exhibited a less favorable cognitive and neurophysiological profile than controls, with lower CRI-q, MoCA, and MIQ scores and longer bilateral N2–P3 inter-peak latencies, despite comparable peripheral hearing, pointing to central cognitive–neural rather than sensory differences. This finding could be interpreted as reflecting baseline differences linked to the socio-cultural characteristics of the choir group, recruited within the Libertà district of Bari, an area characterized by socioeconomic disadvantage. On the contrary, control groups originated from more advantaged districts within the same metropolitan area. Correlation analyses showed coherent within- and cross-domain clustering: melodic discrimination (MDT) correlated with audiometric thresholds, consistent with links between pitch processing and fine-grained auditory resolution (Parbery-Clark et al., 2011; Grassi et al., 2017), whereas weaker EDT and RAT associations suggest greater reliance on higher-order cognition.

In fact, musical and cognitive measures were tightly coupled: cognitive reserve and fluid reasoning related to rhythmic aptitude, and global cognition to multiple music tasks, supporting the role of executive resources in musical ability (Criscuolo et al., 2019). Fluid reasoning also correlated with right-hemisphere N2 amplitude, and amplitude–latency couplings indicated integrated auditory–cognitive cascades (Polich, 2007; Alain et al., 2018), with lateralization patterns consistent with prior ageing research (Divenyi & Haupt, 1997; Alain et al., 2014; Bidelman & Alain, 2015).

### Neurophysiological evidence

Neurophysiological data were acquired using a standardized clinical AEP setup (Cz reference, low impedances <5 kΩ, ER-3A insert earphones, auditory oddball paradigm). Although less spatially dense than high-density EEG, such configurations are optimized for reliability and reproducibility and are widely used in both research and clinical contexts. Comparable protocols have established normative P300 values in older adults (Cóser et al., 2010; Didoné et al., 2016), validated cortical AEPs for objective auditory assessment (Van Dun et al., 2015), and demonstrated high test–retest stability even in biometric applications (Seha & Hatzinakos, 2018), supporting the robustness of the present approach.

The N2-P3 IPL has been shown to be particularly sensitive to cognitive status in older adults, including MCI populations (Papaliagkas et al., 2008, 2011), and indexes the temporal coupling between successive processing stages, i.e., from novelty detection and conflict monitoring (N2) to higher-order cognitive evaluation and context updating (P3; Polich, 2007; Sur & Sinha, 2009; Picton, 2011; Luck, 2014). Age-related slowing of this cascade is well documented (Gaál et al., 2007; Kropotov et al., 2016; Tarawneh et al., 2021), making IPLs particularly informative markers of ageing-related neural inefficiency.

Although direct evidence on training-induced modulation of N2–P3 IPL remains limited, converging work indicates that auditory–cognitive training enhances neural timing and synchrony at both cortical and subcortical levels in older adults (Anderson & Kraus, 2013; Skoe & Kraus, 2024; Fisher et al., 2025). Importantly, N2–P3 IPL captures the efficiency of information transfer between perceptual–attentional and cognitive updating stages, whereas P3 latency reflects the cumulative duration of earlier sensory and cognitive processes (Pihlaja et al., 2023). The selective IPL shortening observed here therefore suggests an acceleration of higher-order cognitive integration rather than global sensory speeding.

N2 and P3 are generated by partially distinct but interacting networks, including medial frontal (ACC/pre-SMA) and posterior parietal–temporal regions, respectively (Halgren et al., 1998; Folstein & Van Petten, 2008). Efficient communication between these regions depends on fronto-parietal connectivity and theta-band synchrony supporting cognitive control and memory updating (Sauseng et al., 2010; Cavanagh & Frank, 2014). Ageing is associated with reduced conduction velocity and weakened theta coherence across these networks (Cummins & Finnigan, 2007; Andrews-Hanna et al., 2007). The choir-specific N2–P3 IPL reduction, together with behavioral gains, may therefore reflect strengthened fronto-parietal coupling and improved temporal coordination within auditory–cognitive circuits.

Absolute peak latencies did not show a consistent training-related modulation. P2 and P3 latencies were unaffected, and although N2 latency yielded a significant Time × Group interaction, post hoc comparisons revealed no significant within-group longitudinal changes, with a pattern not consistent with the hypothesized facilitation. Likewise, the earlier P2–N2 inter-peak interval did not change significantly. Given that the N2–P2 interval primarily reflects early cortical discrimination and perceptual encoding within auditory regions (Näätänen & Picton, 1987; Crowley & Colrain, 2004), and that P2 modulation has been more closely associated with stimulus-specific auditory learning than higher-order executive integration (Tremblay et al., 2001; Alain et al., 2010), the absence of effects at these stages suggests that the intervention did not globally accelerate early sensory processing. Rather, plasticity appears to have selectively targeted later integrative stages, as indexed by the N2–P3 interval.

The lack of robust single-peak modulation is consistent with prior reports highlighting variability in P2, N2, and P3 effects depending on task demands, recording parameters, and heterogeneity in ageing cohorts (Luck, 2014; Alain et al., 2014; Kropotov et al., 2016). This variability parallels the broader literature on music-related auditory plasticity, where cortical ERP changes are frequently observed but differ in magnitude and direction (Sanju & Kumar, 2016; Brown et al., 2013; Okhrei et al., 2018), and where large-scale evidence suggests limited generalization to early subcortical encoding (Whiteford et al., 2025).

Consistent with this interpretation, no longitudinal changes were observed in pure-tone thresholds or speech-in-noise performance. Given the slow rate of age-related threshold decline (≈1 dB/year from age 60; Lee et al., 2005), the absence of peripheral changes is expected. Prior work similarly shows that short-to medium-term musical interventions preferentially induce central neurophysiological modulation, without concomitant improvements in audiometric thresholds or speech-in-noise scores (Grassi et al., 2017; Fleming et al., 2019; Hennessy et al., 2022; Feng et al., 2020). When behavioral gains in challenging listening conditions do emerge, they are more consistently mediated by central mechanisms such as auditory working memory, attentional control, and cortical recruitment (Zendel et al., 2019; Guo et al., 2021; Neves et al., 2023).

Overall, the choir-specific shortening of N2–P3 IPL in the absence of peripheral hearing changes identifies this interval as a sensitive marker of central auditory–cognitive plasticity. This finding aligns with evidence that musical engagement sharpens neural timing and mitigates age-related slowing (Parbery-Clark et al., 2011; Bidelman et al., 2015; Lu et al., 2022; Bonetti et al., 2024) and with reports linking P300 timing to responsiveness to auditory–cognitive training (Ferguson & Henshaw, 2015; Skoe & Kraus, 2024; Pearson et al., 2025). Importantly, converging evidence from the language domain shows that music training is associated with enhanced processing speed and cognitive control in both auditory and visual modalities, particularly in bilingual musicians, supporting shared temporal and executive mechanisms across music and language (Bugos et al., 2026).

The observed gains in this community-based cohort, despite baseline cognitive and neural disadvantages in the choir group, are consistent with the broader literature. Contextual factors are known to shape cognitive trajectories and vulnerability to age-related disorders, plausibly accounting for the lower initial performance observed in the choir group. Crucially, longitudinal mixed-effects models adjusted for baseline outcome levels, reducing the likelihood that observed changes merely reflect regression to the mean and supporting the interpretation of genuine longitudinal improvement. These findings align with growing evidence that cognitively and socially enriching activities can engage compensatory mechanisms in populations at elevated risk of decline.

### Multidimensional Frailty

At baseline, groups did not differ in Selfy-MPI scores, supporting the interpretation of subsequent changes as longitudinal modulation rather than pre-existing imbalance. At T1, higher frailty was associated with poorer cognitive performance and slower auditory–cortical processing, consistent with evidence that frailty predicts accelerated decline in global cognition and executive function and shares neurobiological substrates with cognitive vulnerability (Zhang et al., 2025; Skalska et al., 2025; Carcelén-Fraile et al., 2025). Within the Comprehensive Geriatric Assessment (CGA)–based framework underlying the SELFY-MPI, frailty is conceptualized as a loss of harmonic interaction among interdependent biological, functional, cognitive, and psychosocial domains, leading to reduced homeostatic stability (Pilotto et al., 2019, 2020). The observed cognitive and neurophysiological associations are therefore coherent with a multisystem view in which central nervous system efficiency represents one embedded component of the broader frailty architecture.

Although the longitudinal mixed model did not reveal a significant Time × Group interaction, exploratory contrasts suggested lower frailty among participants exposed to richer multidomain engagement and higher cumulative activity levels. While interpreted cautiously, this pattern is theoretically consistent with the multidimensional structure of the MPI, which parallels the rationale of multidomain preventive strategies targeting mobility, cognition, and social participation simultaneously. Accordingly, multidomain interventions show superior effects on frailty trajectories compared with single-component approaches (de Souto Barreto et al., 2018; Yu et al., 2020), in line with large prevention frameworks such as FINGER, AGELESS, and SUPERBRAIN (Ngandu et al., 2015; Rosenberg et al., 2018; Deckers et al., 2024; Ponvel et al., 2025; Ong et al., 2025; Moon et al., 2025). The dose–response trend further accords with longitudinal evidence that sustained social and cognitive engagement reduces progression toward frailty (Rogers & Fancourt, 2020; Goto & Yamatsu, 2025), reinforcing the conceptualization of frailty as a dynamic and potentially modifiable multisystem condition (Pilotto et al., 2022).

Interestingly, the neurophysiological findings observed in the present study may provide a biologically plausible mechanism underlying the frailty patterns. The reduction in N2–P3 IPL observed in the choir group suggests faster auditory–cortical processing and improved neural efficiency. Event-related potentials such as the P300 are widely considered markers of attentional allocation and cognitive processing speed in ageing populations and have been proposed as neurophysiological indicators of cognitive vulnerability and cognitive frailty in older adults (Pavarini et al., 2018; Tanaka et al., 2024). Because cognitive processing speed and executive control contribute to several domains included in multidimensional frailty frameworks such as the MPI, improvements in neural processing efficiency may plausibly support broader resilience against frailty-related decline.

### Physicological well-being

The Flourishing Scale primarily captures eudaimonic dimensions (i.e., purpose, meaning, relational connectedness, and perceived contribution) rather than cognitive efficiency per se. Indeed, in early-stage dementia, flourishing relates more strongly to quality of life, mood, and self-efficacy than to measurable cognitive impairment (Clarke et al., 2025).

Longitudinal mixed-effects models showed no significant main effects or Time × Group interaction, indicating overall stability of global well-being across the intervention period. This pattern aligns with meta-analytic evidence that behavioral interventions in generally healthy older adults produce small and heterogeneous effects on well-being, with stronger shifts typically observed in clinically distressed populations (Weiss et al., 2016). Notwithstanding the absence of omnibus longitudinal effects, planned contrasts revealed higher flourishing at follow-up in the Choir group relative to the Passive Control group. This selective difference suggests that structured, socially embedded engagement may confer modest psychosocial advantages compared to inactivity, while diverse forms of active participation yield broadly comparable well-being levels. Consistent with this interpretation, choir-based interventions have been shown to reduce loneliness and enhance social integration even when cognitive or physical indices remain unchanged (Pentikäinen et al., 2021; Johnson et al., 2020). Together, these findings indicate that multidomain engagement may not substantially elevate global flourishing in already well-functioning older adults over a medium-duration period, but may help buffer against psychosocial stagnation associated with disengagement.

### Strengths and limitations

Strengths include the ecologically valid, choir-centered design; the multidomain assessment spanning cognition and auditory neurophysiology; and the formal outcome-level comparison demonstrating the dominance of the N2–P3 effect. Limitations concern the non-randomized allocation with baseline imbalances (addressed statistically but not fully eliminated), attrition that may bias generalizability, and substantial variability in the activities undertaken both within the choir’s multidomain program and across control groups. Such heterogeneity, combined with the relatively short time between the two assessment points, may have diluted some effects that could emerge more clearly in a longer and more standardized trial. Our electrophysiology used a clinical AEP setup optimized for translational use; high-density EEG/MEG could refine source-level inferences in future work.

### Clinical implications and future directions

Clinically, these findings position choir-based multidomain engagement as a scalable, socially embedded strategy to support cognitive efficiency in later life, particularly reasoning-related functions and reserve-linked behaviors, while enhancing the temporal coordination of auditory–cortical processing. The selective shortening of the N2–P3 interval suggests that such engagement strengthens network-level integration between conflict monitoring and cognitive updating rather than uniformly accelerating early sensory stages, consistent with models proposing that musical training refines temporal prediction and attentional dynamics, thereby reducing neural processing costs (Parbery-Clark et al., 2011; Bidelman et al., 2015; Lu et al., 2022; Zhang et al., 2025). Converging neuroimaging evidence indicates that lifelong musical engagement is associated with preserved functional connectivity, cerebellar and auditory structural adaptations, and compensatory prefrontal and reward-system recruitment, all of which may contribute to resilience against age-related decline (Marie et al., 2023; Zhang & Tremblay, 2025; Ma et al., 2025; Espinosa et al., 2025).

The MultiMusic intervention occupies an intermediate position between primary and secondary prevention, targeting community-dwelling older adults without dementia but with potential early vulnerability profiles. Its activity-based multidomain structure converges with large-scale lifestyle prevention programs that have demonstrated the feasibility and efficacy of coordinated cognitive, physical, and social stimulation in modifying age-related trajectories. Yet, it differs from conventional multicomponent programs by embedding coordinated cognitive, sensorimotor, emotional, and social stimulation within a single meaningful activity. Choral singing simultaneously recruits auditory perception, vocal–motor synchronization, executive control, memory, and structured social interaction, i.e., processes central to both cognitive reserve and psychosocial well-being (Pentikäinen et al., 2021; Moisseinen et al., 2024; Raja et al., 2025). In preventive samples with largely intact baseline cognition, electrophysiological indices such as N2–P3 provide sensitive markers of functional adaptation that may precede detectable behavioral change, overcoming ceiling effects common in standard neuropsychological testing (Kujala & Näätänen, 2010).

From a public health perspective, community choirs represent low-cost, non-pharmacological interventions implementable within existing cultural and senior infrastructures, without reliance on specialist medical settings. Economic evaluations suggest that community singing and multidomain lifestyle programs can achieve favorable cost-effectiveness profiles while improving mental health–related quality of life and reducing psychosocial risk factors such as loneliness (Melis et al., 2008; Coulton et al., 2015; Hoang et al., 2022). By engaging individuals who are not yet patients but may already exhibit early frailty or cognitive vulnerability, stimulating activities and programs may contribute to maintaining intrinsic capacity and slowing multidimensional decline (Liang et al., 2024; Sánchez-Sánchez et al., 2022). Building on this feasibility study, the forthcoming RCT should mitigate baseline imbalances, extend follow-up duration to assess persistence of effects, and increase sample size to enable stratified analyses by hearing and cognitive status. Given emerging longitudinal evidence for sustained cognitive and neural benefits of late-life musical engagement (Zendel et al., 2019; Marie et al., 2023; Wang et al., 2025; Moisseinen et al., 2025), a sufficiently powered RCT is warranted to determine efficacy, mechanisms, and boundaries of effect.

## Ackowledgments

We would like to express our sincere gratitude to Giulia Rosangela Calfapietro, choir director of the Libertà neighborhood in Bari (Italy), for her unwavering dedication and support to both the participants and this research. Our thanks also go to ANAS Puglia (Associazione Nazionale di Azione Sociale), the Family Services Center of the Libertà neighborhood in Bari, and the association Musica in Gioco in Adelfia for their invaluable collaboration. Finally, we are deeply grateful to all participants, whose commitment and willingness to be involved made each phase of this study possible. This research was supported by EU funding within the MUR PNRR A novel public private alliance to generate socioeconomic, biomedical and technological solutions for an inclusive Italian ageing society (project n. PE00000015, AGE-IT).

