## Supplementary material for "Promoting Healthy Ageing through Choral Training: A 9-Month Multidomain Intervention (MultiMusic) Enhances Neural Processing Speed in Community-Dwelling Older Adults": Cluster_Analysis_APA_Tables.docx

### 1. Cluster Analysis at T0

#### 1.1. Table 1. Cluster membership counts.

| **Cluster** | **n** | **%** |
| --- | --- | --- |
| 1 | 8 | 8.4 |
| 2 | 23 | 24.2 |
| 3 | 33 | 34.7 |
| 4 | 15 | 15.8 |
| 5 | 16 | 16.8 |

Note. Percentages are based on complete-case subjects (listwise deletion). Clustering distance: 1 - |correlation| among subjects.

#### 1.2. Table 2. Means (SD) of variables by cluster.

Table 2. Means (SD) of variables by cluster at T0 (subject clusters, 1 - |r|).

| **Variable** | **Cluster 1** | **Cluster 2** | **Cluster 3** | **Cluster 4** | **Cluster 5** |
| --- | --- | --- | --- | --- | --- |
| CRI Q | 113.38 (16.06) | 100.87 (16.45) | 110.76 (21.28) | 96.73 (18.70) | 103.62 (19.02) |
| EDT | 0.26 (0.94) | 0.11 (0.85) | 0.19 (1.22) | 0.07 (0.95) | -0.21 (0.84) |
| Intellection left | 100.00 (0.00) | 95.65 (11.61) | 93.33 (12.91) | 97.33 (5.94) | 86.25 (22.17) |
| Intellection right | 100.00 (0.00) | 95.22 (14.42) | 93.64 (12.45) | 97.33 (5.94) | 85.62 (24.49) |
| MDT | -0.79 (0.88) | -1.42 (0.93) | -1.11 (0.91) | -1.24 (1.57) | -1.14 (0.92) |
| MIQ | -2.49 (0.98) | -2.67 (1.10) | -2.48 (1.23) | -2.37 (1.25) | -2.45 (1.27) |
| MPT | -0.17 (0.59) | -0.69 (1.04) | -0.84 (1.02) | -0.95 (1.26) | -0.63 (1.01) |
| Matrix test | 2.21 (0.53) | 2.20 (1.76) | 2.33 (1.26) | 1.15 (2.58) | 1.29 (2.42) |
| MoCa | 23.50 (2.98) | 23.26 (2.83) | 22.79 (2.91) | 21.20 (5.72) | 23.31 (5.93) |
| N2 ampl dx | 425.93 (454.20) | 4.67 (3.43) | 3.84 (2.32) | 4.33 (1.75) | 28.35 (100.45) |
| N2 ampl sx | 124.82 (343.53) | 12.19 (35.67) | 4.23 (2.19) | 3.66 (2.18) | 3.45 (2.00) |
| N2 lat dx | 331.89 (19.31) | 341.54 (22.90) | 340.68 (25.55) | 338.81 (31.86) | 330.62 (17.14) |
| N2 lat sx | 345.29 (18.67) | 343.57 (32.45) | 336.97 (20.34) | 335.08 (27.01) | 334.79 (22.28) |
| P2 N2 dx | 33.70 (21.81) | 36.91 (16.26) | 39.19 (16.37) | 38.86 (11.33) | 34.42 (12.04) |
| P2 N2 sx | 42.50 (17.09) | 33.26 (13.48) | 35.61 (12.66) | 29.61 (11.86) | 30.34 (10.51) |
| P2 lat dx | 301.30 (25.23) | 304.29 (35.02) | 298.28 (33.50) | 299.64 (38.28) | 297.14 (18.69) |
| P2 lat sx | 316.25 (33.74) | 314.44 (40.18) | 300.96 (22.91) | 305.17 (32.07) | 304.25 (19.82) |
| P3 ampl dx | 3.07 (2.73) | 4.80 (3.20) | 4.24 (2.45) | 4.99 (2.95) | 4.03 (1.97) |
| P3 ampl sx | 104.85 (282.91) | 7.19 (8.95) | 4.94 (2.24) | 5.30 (2.52) | 4.30 (2.64) |
| P3 lat dx | 371.77 (34.25) | 382.14 (21.16) | 384.00 (22.66) | 387.75 (16.86) | 374.84 (30.64) |
| P3 lat sx | 386.72 (21.11) | 383.47 (34.59) | 379.18 (21.39) | 382.97 (15.35) | 371.19 (27.86) |
| PTA left | -22.69 (4.62) | -25.09 (9.76) | -25.37 (7.75) | -27.87 (10.31) | -30.59 (12.77) |
| PTA right | -22.16 (3.72) | -25.92 (10.11) | -26.84 (8.68) | -29.78 (8.36) | -31.08 (12.95) |
| RAT | -1.10 (0.95) | -1.11 (0.78) | -0.92 (0.89) | -1.03 (0.73) | -0.90 (0.88) |
| SRT left | -29.62 (5.01) | -32.04 (7.61) | -32.06 (8.07) | -32.87 (9.36) | -35.75 (11.95) |
| SRT right | -29.00 (4.54) | -33.39 (9.16) | -31.88 (7.77) | -33.73 (8.48) | -35.81 (12.31) |
| n2p3 dx | 39.95 (16.76) | 40.45 (15.65) | 43.77 (13.05) | 48.14 (20.28) | 40.60 (17.04) |
| n2p3 sx | 39.79 (15.02) | 40.00 (16.41) | 42.26 (11.02) | 47.89 (23.88) | 35.56 (16.92) |
