## Supplementary material for "Promoting Healthy Ageing through Choral Training: A 9-Month Multidomain Intervention (MultiMusic) Enhances Neural Processing Speed in Community-Dwelling Older Adults": flourishing.docx

T0

| **domain** | **variable** | **n** | **rho** | **p** | **p_fdr** |
| --- | --- | --- | --- | --- | --- |
| **Cognitive** | MoCa | 51 | 34 | 813 | 822 |
| **Cognitive** | MIQ | 51 | -32 | 822 | 822 |
| **Cognitive** | CRI_Q | 51 | -146 | 307 | 822 |
| **Musical** | MDT | 51 | -157 | 272 | 626 |
| **Musical** | MPT | 51 | -29 | 841 | 841 |
| **Musical** | EDT | 51 | 144 | 313 | 626 |
| **Musical** | RAT | 51 | 42 | 771 | 841 |
| **Audiometric** | PTA_right | 51 | 287 | 41 | 246 |
| **Audiometric** | PTA_left | 51 | 196 | 169 | 338 |
| **Audiometric** | SRT_right | 51 | 224 | 114 | 338 |
| **Audiometric** | SRT_left | 51 | 0.17 | 233 | 349 |
| **Audiometric** | Intellection_right | 51 | -138 | 333 | 371 |
| **Audiometric** | Intellection_left | 51 | -128 | 371 | 371 |
| **Neurophys_Latency** | P2_lat_dx | 51 | -118 | 409 | 936 |
| **Neurophys_Latency** | N2_lat_dx | 51 | -163 | 253 | 842 |
| **Neurophys_Latency** | P3_lat_dx | 51 | -216 | 128 | 0.64 |
| **Neurophys_Latency** | P2_N2_dx | 51 | 45 | 753 | 936 |
| **Neurophys_Latency** | P2_lat_sx | 51 | -44 | 758 | 936 |
| **Neurophys_Latency** | N2_lat_sx | 51 | 32 | 824 | 936 |
| **Neurophys_Latency** | P3_lat_sx | 51 | -287 | 0.0413 | 413 |
| **Neurophys_Latency** | P2_N2_sx | 51 | -12 | 936 | 936 |
| **Neurophys_Latency** | n2p3_dx | 51 | -72 | 618 | 936 |
| **Neurophys_Latency** | n2p3_sx | 51 | 17 | 904 | 936 |
| **Neurophys_Amplitude** | P3_ampl_dx | 51 | 84 | 557 | 687 |
| **Neurophys_Amplitude** | P3_ampl_sx | 50 | -73 | 616 | 687 |
| **Neurophys_Amplitude** | N2_ampl_dx | 51 | -58 | 687 | 687 |
| **Neurophys_Amplitude** | N2_ampl_sx | 51 | -83 | 563 | 687 |

T1

| **domain** | **variable** | **n** | **rho** | **p** | **p_fdr** |
| --- | --- | --- | --- | --- | --- |
| **Cognitive** | MoCa | 52 | -115 | 415 | 629 |
| **Cognitive** | MIQ | 52 | 106 | 457 | 629 |
| **Cognitive** | CRI_Q | 52 | -69 | 629 | 629 |
| **Musical** | MDT | 52 | -167 | 235 | 448 |
| **Musical** | MPT | 52 | -108 | 448 | 448 |
| **Musical** | EDT | 52 | -114 | 0.42 | 448 |
| **Musical** | RAT | 52 | -201 | 152 | 448 |
| **Audiometric** | PTA_right | 52 | 0.22 | 116 | 0.44 |
| **Audiometric** | PTA_left | 52 | 204 | 147 | 0.44 |
| **Audiometric** | SRT_right | 52 | 122 | 0.39 | 0.78 |
| **Audiometric** | SRT_left | 52 | 0.09 | 526 | 789 |
| **Audiometric** | Intellection_right | 52 | -31 | 826 | 847 |
| **Audiometric** | Intellection_left | 52 | -27 | 847 | 847 |
| **Neurophys_Latency** | P2_lat_dx | 46 | -265 | 0.0746 | 373 |
| **Neurophys_Latency** | N2_lat_dx | 51 | 1 | 993 | 993 |
| **Neurophys_Latency** | P3_lat_dx | 51 | -131 | 0.36 | 896 |
| **Neurophys_Latency** | P2_N2_dx | 44 | 206 | 181 | 602 |
| **Neurophys_Latency** | P2_lat_sx | 45 | -335 | 0.0244 | 244 |
| **Neurophys_Latency** | N2_lat_sx | 51 | 22 | 876 | 974 |
| **Neurophys_Latency** | P3_lat_sx | 51 | 26 | 858 | 974 |
| **Neurophys_Latency** | P2_N2_sx | 45 | -26 | 867 | 974 |
| **Neurophys_Latency** | n2p3_dx | 51 | 32 | 826 | 974 |
| **Neurophys_Latency** | n2p3_sx | 51 | -109 | 448 | 896 |
| **Neurophys_Amplitude** | P3_ampl_dx | 51 | 125 | 382 | 763 |
| **Neurophys_Amplitude** | P3_ampl_sx | 51 | -35 | 809 | 809 |
| **Neurophys_Amplitude** | N2_ampl_dx | 43 | -192 | 216 | 763 |
| **Neurophys_Amplitude** | N2_ampl_sx | 44 | 83 | 592 | 0.79 |
