## Supplementary material for "Promoting Healthy Ageing through Choral Training: A 9-Month Multidomain Intervention (MultiMusic) Enhances Neural Processing Speed in Community-Dwelling Older Adults": selfy_correlations.docx

| **variable** | **n0** | **r0** | **p0** | **n1** | **r1** | **p1** | **z_change** | **p_change** | **domain** | **p0_fdr** | **p1_fdr** | **pchange_fdr** |
| --- | --- | --- | --- | --- | --- | --- | --- | --- | --- | --- | --- | --- |
| **N2_ampl_dx** | 54 | -0.154919925379391 | 0.263329727736289 | 44 | 0.215487173321358 | 0.160081693490858 | 1.78823942689201 | 0.0737373878108055 | AEP_amplitude | 0.844676420809932 | 0.21344225798781 | 0.237973241842811 |
| **N2_ampl_sx** | 54 | -0.0663649865312862 | 0.633507315607449 | 45 | 0.252791056782015 | 0.0938579681475997 | 1.55904123463413 | 0.118986620921405 | AEP_amplitude | 0.844676420809932 | 0.21344225798781 | 0.237973241842811 |
| **P3_ampl_dx** | 54 | 0.00751592930948787 | 0.956983785582695 | 53 | 0.196435984076165 | 0.158612724664797 | 0.962262109961839 | 0.335917956858736 | AEP_amplitude | 0.956983785582695 | 0.21344225798781 | 0.353007921727263 |
| **P3_ampl_sx** | 53 | 0.0874397418870473 | 0.533556613468297 | 53 | -0.0977771331230077 | 0.486099299759833 | -0.928770719838557 | 0.353007921727263 | AEP_amplitude | 0.844676420809932 | 0.486099299759833 | 0.353007921727263 |
| **N2_lat_dx** | 54 | -0.0280760037944269 | 0.840285920967516 | 53 | 0.130135209848052 | 0.353010680312691 | 0.7987288363402 | 0.424447662324224 | AEP_latency | 0.987887266077969 | 0.538078938998357 | 0.521832470309003 |
| **N2_lat_sx** | 54 | -0.23645264858778 | 0.0851760931824242 | 53 | 0.162357823774366 | 0.245428763697435 | 2.03409929895991 | 0.0419415846499004 | AEP_latency | 0.283920310608081 | 0.490857527394869 | 0.139805282166335 |
| **P2_N2_dx** | 54 | -0.454709087038872 | 0.000551857419028701 | 45 | 0.0617150630627658 | 0.687145714288713 | 2.65114399966638 | 0.0080219631577335 | AEP_latency | 0.00551857419028701 | 0.76349523809857 | 0.0401098157886675 |
| **P2_N2_sx** | 54 | 0.32171495885116 | 0.0176805610696551 | 46 | -0.329546901457407 | 0.0253211899468005 | -3.26455645472003 | 0.00109635612063297 | AEP_latency | 0.0884028053482754 | 0.126605949734002 | 0.0109635612063297 |
| **P2_lat_dx** | 54 | 0.0924110842415349 | 0.506296974653951 | 47 | 0.380146372280813 | 0.00839544033458259 | 1.49476442292921 | 0.134975934343775 | AEP_latency | 0.987887266077969 | 0.0839544033458259 | 0.337439835859437 |
| **P2_lat_sx** | 54 | 0.0519487134723575 | 0.709101380925596 | 46 | 0.256023357975305 | 0.0859058546027163 | 1.01360748799783 | 0.310770083226387 | AEP_latency | 0.987887266077969 | 0.286352848675721 | 0.521832470309003 |
| **P3_lat_dx** | 54 | -6.89766033216197e-05 | 0.999605037653243 | 53 | 0.167412475349777 | 0.230840179674801 | 0.849536203633038 | 0.395582993830371 | AEP_latency | 0.999605037653243 | 0.490857527394869 | 0.521832470309003 |
| **P3_lat_sx** | 54 | -0.0585621615100052 | 0.6740223177402 | 53 | -0.0319977542216113 | 0.820073322648344 | 0.133760127587561 | 0.89359225736276 | AEP_latency | 0.987887266077969 | 0.820073322648344 | 0.89359225736276 |
| **n2p3_dx** | 54 | 0.0378841899141323 | 0.785641705571624 | 53 | -0.105601837445665 | 0.451709016213336 | -0.723049907082021 | 0.469649223278103 | AEP_latency | 0.987887266077969 | 0.56463627026667 | 0.521832470309003 |
| **n2p3_sx** | 54 | -0.0194288880559865 | 0.889098539470172 | 53 | 0.123923111465248 | 0.37665525729885 | 0.72352901953805 | 0.469354931958046 | AEP_latency | 0.987887266077969 | 0.538078938998357 | 0.521832470309003 |
| **Intellection_left** | 54 | -0.194263036557412 | 0.159254532401309 | 54 | -0.172456758887954 | 0.212394565388885 | 0.113952633263003 | 0.909275341950112 | Audiometric | 0.159254532401309 | 0.212394565388885 | 0.916099573180185 |
| **Intellection_right** | 54 | -0.199153294696519 | 0.148813418409535 | 54 | -0.179038031541101 | 0.195193947412385 | 0.105348130029736 | 0.916099573180185 | Audiometric | 0.159254532401309 | 0.212394565388885 | 0.916099573180185 |
| **Matrix_test** | 52 | 0.266122326995815 | 0.0565296494178955 | 52 | 0.43738965851619 | 0.0011852437068761 | 0.971696970245334 | 0.331201324501334 | Audiometric | 0.159254532401309 | 0.00355573112062829 | 0.916099573180185 |
| **CRI_Q** | 54 | -0.187965425479231 | 0.173480292220406 | 54 | -0.272907847417481 | 0.0458678568120049 | -0.453345318669985 | 0.650300104623275 | Cognitive | 0.173480292220406 | 0.0458678568120049 | 0.650300104623275 |
| **MIQ** | 54 | -0.244958769727175 | 0.0742190351609772 | 54 | -0.384359231370477 | 0.00411140206923735 | -0.783326952247393 | 0.43343513990741 | Cognitive | 0.130050591951904 | 0.00616710310385603 | 0.650152709861115 |
| **MoCa** | 54 | -0.235338730890516 | 0.0867003946346027 | 54 | -0.465842992482307 | 0.000385378438761428 | -1.33775119350845 | 0.180977556708742 | Cognitive | 0.130050591951904 | 0.00115613531628428 | 0.542932670126226 |
| **EDT** | 54 | -0.312271413717699 | 0.0215140496565414 | 54 | -0.135845663017841 | 0.327357875582619 | 0.941120369218338 | 0.346643178217809 | Musical | 0.0430280993130827 | 0.732667866299781 | 0.346643178217809 |
| **MDT** | 54 | -0.195435746369651 | 0.156702989180991 | 54 | 0.0221664350443384 | 0.87359211891279 | 1.11171649250588 | 0.266260066404859 | Musical | 0.156702989180991 | 0.87359211891279 | 0.346643178217809 |
| **MPT** | 54 | -0.36793469456266 | 0.00619582442436955 | 54 | -0.125379634583582 | 0.36633393314989 | 1.31288224688997 | 0.189222629931717 | Musical | 0.0247832976974782 | 0.732667866299781 | 0.346643178217809 |
| **RAT** | 54 | -0.284644074827519 | 0.0369674469618151 | 54 | -0.0435948971868648 | 0.754274085970209 | 1.25792279400918 | 0.208419679291448 | Musical | 0.0492899292824201 | 0.87359211891279 | 0.346643178217809 |
