## Supplementary material for "Promoting Healthy Ageing through Choral Training: A 9-Month Multidomain Intervention (MultiMusic) Enhances Neural Processing Speed in Community-Dwelling Older Adults": T0_interdomain_FINAL_.docx

### 1. T0 Inter-domain correlations

#### 1.1. Cognitive <-> Musical

| Var_A | Var_B | r | p | q | Method |
| --- | --- | --- | --- | --- | --- |
| mo_ca | mdt | 0.536 | 0.001 | 0.001 | Spearman |
| mo_ca | mpt | 0.479 | 0.001 | 0.001 | Spearman |
| mo_ca | rat | 0.474 | 0.001 | 0.001 | Spearman |
| mo_ca | edt | 0.405 | 0.002 | 0.004 | Spearman |
| miq | mdt | 0.440 | 0.001 | 0.002 | Spearman |
| miq | mpt | 0.197 | 0.153 | 0.153 | Spearman |
| miq | rat | 0.511 | 0.001 | 0.001 | Spearman |
| miq | edt | 0.379 | 0.005 | 0.007 | Spearman |
| cri_q | mdt | 0.241 | 0.079 | 0.086 | Spearman |
| cri_q | mpt | 0.268 | 0.050 | 0.060 | Spearman |
| cri_q | rat | 0.556 | 0.001 | 0.001 | Spearman |
| cri_q | edt | 0.294 | 0.031 | 0.041 | Spearman |

#### 1.2. Cognitive <-> Audiometric

| Var_A | Var_B | r | p | q | Method |
| --- | --- | --- | --- | --- | --- |
| mo_ca | pta_right (x -1) | 0.389 | 0.004 | 0.058 | Spearman |
| mo_ca | pta_left (x -1) | 0.326 | 0.016 | 0.067 | Spearman |
| mo_ca | srt_right (x -1) | 0.250 | 0.068 | 0.204 | Spearman |
| mo_ca | srt_left (x -1) | 0.284 | 0.038 | 0.132 | Spearman |
| mo_ca | matrix_test (x -1) | 0.350 | 0.011 | 0.058 | Spearman |
| mo_ca | intellection_right | 0.358 | 0.008 | 0.058 | Spearman |
| mo_ca | intellection_left | 0.346 | 0.010 | 0.058 | Spearman |
| miq | pta_right (x -1) | 0.141 | 0.308 | 0.539 | Spearman |
| miq | pta_left (x -1) | 0.144 | 0.299 | 0.539 | Spearman |
| miq | srt_right (x -1) | 0.101 | 0.468 | 0.579 | Spearman |
| miq | srt_left (x -1) | 0.127 | 0.362 | 0.543 | Spearman |
| miq | matrix_test (x -1) | 0.175 | 0.214 | 0.498 | Spearman |
| miq | intellection_right | 0.130 | 0.348 | 0.543 | Spearman |
| miq | intellection_left | 0.065 | 0.640 | 0.708 | Spearman |
| cri_q | pta_right (x -1) | 0.145 | 0.296 | 0.539 | Spearman |
| cri_q | pta_left (x -1) | 0.073 | 0.599 | 0.698 | Spearman |
| cri_q | srt_right (x -1) | 0.046 | 0.741 | 0.778 | Spearman |
| cri_q | srt_left (x -1) | 0.016 | 0.911 | 0.911 | Spearman |
| cri_q | matrix_test (x -1) | 0.231 | 0.099 | 0.259 | Spearman |
| cri_q | intellection_right | 0.110 | 0.427 | 0.579 | Spearman |
| cri_q | intellection_left | 0.101 | 0.468 | 0.579 | Spearman |

#### 1.3. Cognitive <-> Neurophysiologic_amplitude

| Var_A | Var_B | r | p | q | Method |
| --- | --- | --- | --- | --- | --- |
| mo_ca | n2_ampl_dx | 0.189 | 0.171 | 0.684 | Spearman |
| mo_ca | p3_ampl_dx | -0.095 | 0.493 | 0.925 | Spearman |
| mo_ca | n2_ampl_sx | 0.039 | 0.778 | 0.925 | Spearman |
| mo_ca | p3_ampl_sx | -0.085 | 0.545 | 0.925 | Spearman |
| miq | n2_ampl_dx | 0.426 | 0.001 | 0.016 | Spearman |
| miq | p3_ampl_dx | 0.096 | 0.491 | 0.925 | Spearman |
| miq | n2_ampl_sx | -0.066 | 0.636 | 0.925 | Spearman |
| miq | p3_ampl_sx | 0.021 | 0.880 | 0.925 | Spearman |
| cri_q | n2_ampl_dx | 0.211 | 0.126 | 0.684 | Spearman |
| cri_q | p3_ampl_dx | -0.046 | 0.741 | 0.925 | Spearman |
| cri_q | n2_ampl_sx | 0.013 | 0.924 | 0.925 | Spearman |
| cri_q | p3_ampl_sx | -0.013 | 0.925 | 0.925 | Spearman |

#### 1.4. Cognitive <-> Neurophysiologic_latency

| Var_A | Var_B | r | p | q | Method |
| --- | --- | --- | --- | --- | --- |
| mo_ca | p2_lat_dx | 0.106 | 0.444 | 0.667 | Spearman |
| mo_ca | n2_lat_dx | 0.210 | 0.128 | 0.475 | Spearman |
| mo_ca | p3_lat_dx | 0.027 | 0.844 | 0.972 | Spearman |
| mo_ca | p2_n2_dx | -0.034 | 0.809 | 0.971 | Spearman |
| mo_ca | p2_lat_sx | 0.232 | 0.091 | 0.475 | Spearman |
| mo_ca | n2_lat_sx | 0.273 | 0.046 | 0.458 | Spearman |
| mo_ca | p3_lat_sx | 0.159 | 0.252 | 0.590 | Spearman |
| mo_ca | p2_n2_sx | 0.063 | 0.653 | 0.851 | Spearman |
| mo_ca | n2p3_dx | -0.328 | 0.015 | 0.421 | Spearman |
| mo_ca | n2p3_sx | -0.188 | 0.174 | 0.475 | Spearman |
| miq | p2_lat_dx | 0.214 | 0.120 | 0.475 | Spearman |
| miq | n2_lat_dx | 0.207 | 0.132 | 0.475 | Spearman |
| miq | p3_lat_dx | 0.142 | 0.306 | 0.611 | Spearman |
| miq | p2_n2_dx | 0.006 | 0.967 | 0.972 | Spearman |
| miq | p2_lat_sx | 0.147 | 0.290 | 0.611 | Spearman |
| miq | n2_lat_sx | 0.196 | 0.155 | 0.475 | Spearman |
| miq | p3_lat_sx | 0.087 | 0.534 | 0.762 | Spearman |
| miq | p2_n2_sx | 0.005 | 0.972 | 0.972 | Spearman |
| miq | n2p3_dx | -0.189 | 0.171 | 0.475 | Spearman |
| miq | n2p3_sx | -0.119 | 0.393 | 0.660 | Spearman |
| cri_q | p2_lat_dx | 0.116 | 0.406 | 0.660 | Spearman |
| cri_q | n2_lat_dx | 0.112 | 0.418 | 0.660 | Spearman |
| cri_q | p3_lat_dx | -0.008 | 0.955 | 0.972 | Spearman |
| cri_q | p2_n2_dx | -0.077 | 0.579 | 0.789 | Spearman |
| cri_q | p2_lat_sx | 0.157 | 0.256 | 0.590 | Spearman |
| cri_q | n2_lat_sx | 0.115 | 0.408 | 0.660 | Spearman |
| cri_q | p3_lat_sx | -0.034 | 0.808 | 0.971 | Spearman |
| cri_q | p2_n2_sx | 0.006 | 0.964 | 0.972 | Spearman |
| cri_q | n2p3_dx | -0.299 | 0.028 | 0.421 | Spearman |
| cri_q | n2p3_sx | -0.235 | 0.087 | 0.475 | Spearman |

#### 1.5. Musical <-> Audiometric

| Var_A | Var_B | r | p | q | Method |
| --- | --- | --- | --- | --- | --- |
| mdt | pta_right (x -1) | 0.379 | 0.005 | 0.043 | Spearman |
| mdt | pta_left (x -1) | 0.387 | 0.004 | 0.043 | Spearman |
| mdt | srt_right (x -1) | 0.368 | 0.006 | 0.043 | Spearman |
| mdt | srt_left (x -1) | 0.445 | 0.001 | 0.021 | Spearman |
| mdt | matrix_test (x -1) | 0.350 | 0.011 | 0.051 | Spearman |
| mdt | intellection_right | 0.321 | 0.018 | 0.071 | Spearman |
| mdt | intellection_left | 0.351 | 0.009 | 0.051 | Spearman |
| mpt | pta_right (x -1) | 0.218 | 0.113 | 0.197 | Spearman |
| mpt | pta_left (x -1) | 0.236 | 0.085 | 0.161 | Spearman |
| mpt | srt_right (x -1) | 0.262 | 0.056 | 0.149 | Spearman |
| mpt | srt_left (x -1) | 0.262 | 0.055 | 0.149 | Spearman |
| mpt | matrix_test (x -1) | 0.131 | 0.354 | 0.451 | Spearman |
| mpt | intellection_right | 0.213 | 0.122 | 0.201 | Spearman |
| mpt | intellection_left | 0.154 | 0.266 | 0.391 | Spearman |
| rat | pta_right (x -1) | 0.239 | 0.082 | 0.161 | Spearman |
| rat | pta_left (x -1) | 0.244 | 0.075 | 0.161 | Spearman |
| rat | srt_right (x -1) | 0.236 | 0.086 | 0.161 | Spearman |
| rat | srt_left (x -1) | 0.259 | 0.059 | 0.149 | Spearman |
| rat | matrix_test (x -1) | 0.292 | 0.036 | 0.126 | Spearman |
| rat | intellection_right | 0.042 | 0.765 | 0.788 | Spearman |
| rat | intellection_left | 0.075 | 0.591 | 0.662 | Spearman |
| edt | pta_right (x -1) | 0.088 | 0.528 | 0.616 | Spearman |
| edt | pta_left (x -1) | 0.136 | 0.328 | 0.438 | Spearman |
| edt | srt_right (x -1) | 0.051 | 0.715 | 0.770 | Spearman |
| edt | srt_left (x -1) | 0.138 | 0.318 | 0.438 | Spearman |
| edt | matrix_test (x -1) | 0.199 | 0.156 | 0.243 | Spearman |
| edt | intellection_right | 0.038 | 0.788 | 0.788 | Spearman |
| edt | intellection_left | 0.092 | 0.507 | 0.616 | Spearman |

#### 1.6. Musical <-> Neurophysiologic_amplitude

| Var_A | Var_B | r | p | q | Method |
| --- | --- | --- | --- | --- | --- |
| mdt | n2_ampl_dx | 0.265 | 0.053 | 0.422 | Spearman |
| mdt | p3_ampl_dx | 0.095 | 0.495 | 0.663 | Spearman |
| mdt | n2_ampl_sx | 0.045 | 0.746 | 0.796 | Spearman |
| mdt | p3_ampl_sx | 0.143 | 0.307 | 0.546 | Spearman |
| mpt | n2_ampl_dx | 0.178 | 0.198 | 0.529 | Spearman |
| mpt | p3_ampl_dx | 0.047 | 0.738 | 0.796 | Spearman |
| mpt | n2_ampl_sx | 0.195 | 0.157 | 0.529 | Spearman |
| mpt | p3_ampl_sx | -0.021 | 0.882 | 0.882 | Spearman |
| rat | n2_ampl_dx | 0.148 | 0.285 | 0.546 | Spearman |
| rat | p3_ampl_dx | 0.124 | 0.371 | 0.593 | Spearman |
| rat | n2_ampl_sx | 0.291 | 0.033 | 0.422 | Spearman |
| rat | p3_ampl_sx | 0.224 | 0.107 | 0.529 | Spearman |
| edt | n2_ampl_dx | 0.094 | 0.498 | 0.663 | Spearman |
| edt | p3_ampl_dx | 0.163 | 0.238 | 0.545 | Spearman |
| edt | n2_ampl_sx | 0.086 | 0.539 | 0.663 | Spearman |
| edt | p3_ampl_sx | 0.193 | 0.166 | 0.529 | Spearman |

#### 1.7. Musical <-> Neurophysiologic_latency

| Var_A | Var_B | r | p | q | Method |
| --- | --- | --- | --- | --- | --- |
| mdt | p2_lat_dx | 0.042 | 0.760 | 0.981 | Spearman |
| mdt | n2_lat_dx | 0.021 | 0.878 | 0.995 | Spearman |
| mdt | p3_lat_dx | 0.054 | 0.696 | 0.981 | Spearman |
| mdt | p2_n2_dx | -0.003 | 0.985 | 0.995 | Spearman |
| mdt | p2_lat_sx | 0.007 | 0.961 | 0.995 | Spearman |
| mdt | n2_lat_sx | -0.067 | 0.632 | 0.981 | Spearman |
| mdt | p3_lat_sx | -0.107 | 0.442 | 0.981 | Spearman |
| mdt | p2_n2_sx | 0.113 | 0.416 | 0.981 | Spearman |
| mdt | n2p3_dx | 0.047 | 0.734 | 0.981 | Spearman |
| mdt | n2p3_sx | 0.063 | 0.651 | 0.981 | Spearman |
| mpt | p2_lat_dx | -0.063 | 0.650 | 0.981 | Spearman |
| mpt | n2_lat_dx | 0.011 | 0.936 | 0.995 | Spearman |
| mpt | p3_lat_dx | -0.059 | 0.674 | 0.981 | Spearman |
| mpt | p2_n2_dx | 0.008 | 0.957 | 0.995 | Spearman |
| mpt | p2_lat_sx | 0.066 | 0.638 | 0.981 | Spearman |
| mpt | n2_lat_sx | 0.193 | 0.163 | 0.651 | Spearman |
| mpt | p3_lat_sx | -0.001 | 0.994 | 0.995 | Spearman |
| mpt | p2_n2_sx | 0.215 | 0.119 | 0.579 | Spearman |
| mpt | n2p3_dx | -0.067 | 0.629 | 0.981 | Spearman |
| mpt | n2p3_sx | -0.250 | 0.068 | 0.579 | Spearman |
| rat | p2_lat_dx | 0.152 | 0.274 | 0.844 | Spearman |
| rat | n2_lat_dx | 0.246 | 0.073 | 0.579 | Spearman |
| rat | p3_lat_dx | 0.224 | 0.104 | 0.579 | Spearman |
| rat | p2_n2_dx | -0.001 | 0.995 | 0.995 | Spearman |
| rat | p2_lat_sx | 0.151 | 0.274 | 0.844 | Spearman |
| rat | n2_lat_sx | 0.287 | 0.035 | 0.579 | Spearman |
| rat | p3_lat_sx | 0.209 | 0.130 | 0.579 | Spearman |
| rat | p2_n2_sx | 0.064 | 0.645 | 0.981 | Spearman |
| rat | n2p3_dx | -0.186 | 0.179 | 0.651 | Spearman |
| rat | n2p3_sx | -0.101 | 0.468 | 0.981 | Spearman |
| edt | p2_lat_dx | 0.295 | 0.030 | 0.579 | Spearman |
| edt | n2_lat_dx | 0.255 | 0.063 | 0.579 | Spearman |
| edt | p3_lat_dx | 0.234 | 0.088 | 0.579 | Spearman |
| edt | p2_n2_dx | -0.071 | 0.608 | 0.981 | Spearman |
| edt | p2_lat_sx | 0.125 | 0.370 | 0.981 | Spearman |
| edt | n2_lat_sx | 0.139 | 0.315 | 0.901 | Spearman |
| edt | p3_lat_sx | 0.107 | 0.441 | 0.981 | Spearman |
| edt | p2_n2_sx | -0.046 | 0.741 | 0.981 | Spearman |
| edt | n2p3_dx | -0.017 | 0.904 | 0.995 | Spearman |
| edt | n2p3_sx | -0.018 | 0.895 | 0.995 | Spearman |

#### 1.8. Audiometric <-> Neurophysiologic_amplitude

| Var_A | Var_B | r | p | q | Method |
| --- | --- | --- | --- | --- | --- |
| pta_right (x -1) | n2_ampl_dx | 0.015 | 0.913 | 0.979 | Spearman |
| pta_right (x -1) | p3_ampl_dx | -0.120 | 0.388 | 0.838 | Spearman |
| pta_right (x -1) | n2_ampl_sx | 0.097 | 0.487 | 0.838 | Spearman |
| pta_right (x -1) | p3_ampl_sx | -0.118 | 0.398 | 0.838 | Spearman |
| pta_left (x -1) | n2_ampl_dx | -0.086 | 0.538 | 0.838 | Spearman |
| pta_left (x -1) | p3_ampl_dx | -0.145 | 0.297 | 0.838 | Spearman |
| pta_left (x -1) | n2_ampl_sx | -0.004 | 0.979 | 0.979 | Spearman |
| pta_left (x -1) | p3_ampl_sx | -0.151 | 0.279 | 0.838 | Spearman |
| srt_right (x -1) | n2_ampl_dx | -0.041 | 0.768 | 0.979 | Spearman |
| srt_right (x -1) | p3_ampl_dx | -0.211 | 0.126 | 0.838 | Spearman |
| srt_right (x -1) | n2_ampl_sx | -0.023 | 0.868 | 0.979 | Spearman |
| srt_right (x -1) | p3_ampl_sx | -0.217 | 0.118 | 0.838 | Spearman |
| srt_left (x -1) | n2_ampl_dx | -0.011 | 0.934 | 0.979 | Spearman |
| srt_left (x -1) | p3_ampl_dx | -0.131 | 0.344 | 0.838 | Spearman |
| srt_left (x -1) | n2_ampl_sx | 0.006 | 0.963 | 0.979 | Spearman |
| srt_left (x -1) | p3_ampl_sx | -0.141 | 0.313 | 0.838 | Spearman |
| matrix_test (x -1) | n2_ampl_dx | -0.133 | 0.349 | 0.838 | Spearman |
| matrix_test (x -1) | p3_ampl_dx | -0.226 | 0.108 | 0.838 | Spearman |
| matrix_test (x -1) | n2_ampl_sx | -0.093 | 0.512 | 0.838 | Spearman |
| matrix_test (x -1) | p3_ampl_sx | -0.156 | 0.273 | 0.838 | Spearman |
| intellection_right | n2_ampl_dx | 0.065 | 0.642 | 0.899 | Spearman |
| intellection_right | p3_ampl_dx | -0.111 | 0.423 | 0.838 | Spearman |
| intellection_right | n2_ampl_sx | -0.038 | 0.785 | 0.979 | Spearman |
| intellection_right | p3_ampl_sx | -0.141 | 0.316 | 0.838 | Spearman |
| intellection_left | n2_ampl_dx | 0.072 | 0.607 | 0.895 | Spearman |
| intellection_left | p3_ampl_dx | -0.105 | 0.449 | 0.838 | Spearman |
| intellection_left | n2_ampl_sx | -0.021 | 0.880 | 0.979 | Spearman |
| intellection_left | p3_ampl_sx | -0.146 | 0.295 | 0.838 | Spearman |

#### 1.9. Audiometric <-> Neurophysiologic_latency

| Var_A | Var_B | r | p | q | Method |
| --- | --- | --- | --- | --- | --- |
| pta_right (x -1) | p2_lat_dx | 0.068 | 0.625 | 0.975 | Spearman |
| pta_right (x -1) | n2_lat_dx | 0.103 | 0.460 | 0.975 | Spearman |
| pta_right (x -1) | p3_lat_dx | -0.026 | 0.851 | 0.975 | Spearman |
| pta_right (x -1) | p2_n2_dx | -0.034 | 0.808 | 0.975 | Spearman |
| pta_right (x -1) | p2_lat_sx | 0.139 | 0.316 | 0.975 | Spearman |
| pta_right (x -1) | n2_lat_sx | 0.175 | 0.205 | 0.975 | Spearman |
| pta_right (x -1) | p3_lat_sx | 0.116 | 0.402 | 0.975 | Spearman |
| pta_right (x -1) | p2_n2_sx | 0.162 | 0.243 | 0.975 | Spearman |
| pta_right (x -1) | n2p3_dx | -0.049 | 0.724 | 0.975 | Spearman |
| pta_right (x -1) | n2p3_sx | 0.018 | 0.895 | 0.975 | Spearman |
| pta_left (x -1) | p2_lat_dx | 0.065 | 0.640 | 0.975 | Spearman |
| pta_left (x -1) | n2_lat_dx | 0.046 | 0.744 | 0.975 | Spearman |
| pta_left (x -1) | p3_lat_dx | -0.052 | 0.710 | 0.975 | Spearman |
| pta_left (x -1) | p2_n2_dx | -0.039 | 0.782 | 0.975 | Spearman |
| pta_left (x -1) | p2_lat_sx | 0.073 | 0.600 | 0.975 | Spearman |
| pta_left (x -1) | n2_lat_sx | 0.110 | 0.429 | 0.975 | Spearman |
| pta_left (x -1) | p3_lat_sx | 0.043 | 0.755 | 0.975 | Spearman |
| pta_left (x -1) | p2_n2_sx | 0.129 | 0.351 | 0.975 | Spearman |
| pta_left (x -1) | n2p3_dx | -0.013 | 0.923 | 0.975 | Spearman |
| pta_left (x -1) | n2p3_sx | -0.026 | 0.852 | 0.975 | Spearman |
| srt_right (x -1) | p2_lat_dx | -0.002 | 0.986 | 0.986 | Spearman |
| srt_right (x -1) | n2_lat_dx | -0.043 | 0.759 | 0.975 | Spearman |
| srt_right (x -1) | p3_lat_dx | -0.103 | 0.460 | 0.975 | Spearman |
| srt_right (x -1) | p2_n2_dx | -0.052 | 0.710 | 0.975 | Spearman |
| srt_right (x -1) | p2_lat_sx | 0.022 | 0.874 | 0.975 | Spearman |
| srt_right (x -1) | n2_lat_sx | 0.134 | 0.335 | 0.975 | Spearman |
| srt_right (x -1) | p3_lat_sx | 0.034 | 0.809 | 0.975 | Spearman |
| srt_right (x -1) | p2_n2_sx | 0.124 | 0.371 | 0.975 | Spearman |
| srt_right (x -1) | n2p3_dx | -0.016 | 0.911 | 0.975 | Spearman |
| srt_right (x -1) | n2p3_sx | -0.117 | 0.398 | 0.975 | Spearman |
| srt_left (x -1) | p2_lat_dx | -0.038 | 0.786 | 0.975 | Spearman |
| srt_left (x -1) | n2_lat_dx | -0.048 | 0.729 | 0.975 | Spearman |
| srt_left (x -1) | p3_lat_dx | -0.098 | 0.482 | 0.975 | Spearman |
| srt_left (x -1) | p2_n2_dx | -0.016 | 0.911 | 0.975 | Spearman |
| srt_left (x -1) | p2_lat_sx | -0.006 | 0.968 | 0.985 | Spearman |
| srt_left (x -1) | n2_lat_sx | 0.101 | 0.469 | 0.975 | Spearman |
| srt_left (x -1) | p3_lat_sx | 0.014 | 0.919 | 0.975 | Spearman |
| srt_left (x -1) | p2_n2_sx | 0.137 | 0.323 | 0.975 | Spearman |
| srt_left (x -1) | n2p3_dx | 0.019 | 0.893 | 0.975 | Spearman |
| srt_left (x -1) | n2p3_sx | -0.069 | 0.620 | 0.975 | Spearman |
| matrix_test (x -1) | p2_lat_dx | 0.097 | 0.492 | 0.975 | Spearman |
| matrix_test (x -1) | n2_lat_dx | 0.030 | 0.834 | 0.975 | Spearman |
| matrix_test (x -1) | p3_lat_dx | -0.142 | 0.316 | 0.975 | Spearman |
| matrix_test (x -1) | p2_n2_dx | -0.097 | 0.495 | 0.975 | Spearman |
| matrix_test (x -1) | p2_lat_sx | 0.057 | 0.687 | 0.975 | Spearman |
| matrix_test (x -1) | n2_lat_sx | 0.080 | 0.571 | 0.975 | Spearman |
| matrix_test (x -1) | p3_lat_sx | -0.039 | 0.785 | 0.975 | Spearman |
| matrix_test (x -1) | p2_n2_sx | 0.023 | 0.871 | 0.975 | Spearman |
| matrix_test (x -1) | n2p3_dx | -0.194 | 0.168 | 0.975 | Spearman |
| matrix_test (x -1) | n2p3_sx | -0.174 | 0.216 | 0.975 | Spearman |
| intellection_right | p2_lat_dx | -0.031 | 0.825 | 0.975 | Spearman |
| intellection_right | n2_lat_dx | -0.077 | 0.579 | 0.975 | Spearman |
| intellection_right | p3_lat_dx | -0.219 | 0.111 | 0.975 | Spearman |
| intellection_right | p2_n2_dx | -0.034 | 0.805 | 0.975 | Spearman |
| intellection_right | p2_lat_sx | 0.103 | 0.459 | 0.975 | Spearman |
| intellection_right | n2_lat_sx | 0.103 | 0.457 | 0.975 | Spearman |
| intellection_right | p3_lat_sx | 0.026 | 0.853 | 0.975 | Spearman |
| intellection_right | p2_n2_sx | 0.017 | 0.906 | 0.975 | Spearman |
| intellection_right | n2p3_dx | 0.012 | 0.933 | 0.975 | Spearman |
| intellection_right | n2p3_sx | -0.089 | 0.524 | 0.975 | Spearman |
| intellection_left | p2_lat_dx | -0.005 | 0.971 | 0.985 | Spearman |
| intellection_left | n2_lat_dx | -0.043 | 0.760 | 0.975 | Spearman |
| intellection_left | p3_lat_dx | -0.162 | 0.243 | 0.975 | Spearman |
| intellection_left | p2_n2_dx | -0.037 | 0.792 | 0.975 | Spearman |
| intellection_left | p2_lat_sx | 0.062 | 0.658 | 0.975 | Spearman |
| intellection_left | n2_lat_sx | 0.081 | 0.558 | 0.975 | Spearman |
| intellection_left | p3_lat_sx | 0.018 | 0.899 | 0.975 | Spearman |
| intellection_left | p2_n2_sx | 0.108 | 0.435 | 0.975 | Spearman |
| intellection_left | n2p3_dx | 0.038 | 0.784 | 0.975 | Spearman |
| intellection_left | n2p3_sx | -0.079 | 0.568 | 0.975 | Spearman |

#### 1.10. Neurophysiologic_amplitude <-> Neurophysiologic_latency

| Var_A | Var_B | r | p | q | Method |
| --- | --- | --- | --- | --- | --- |
| n2_ampl_dx | p2_lat_dx | -0.109 | 0.435 | 0.678 | Spearman |
| n2_ampl_dx | n2_lat_dx | 0.023 | 0.869 | 0.891 | Spearman |
| n2_ampl_dx | p3_lat_dx | 0.103 | 0.458 | 0.678 | Spearman |
| n2_ampl_dx | p2_n2_dx | 0.218 | 0.113 | 0.329 | Spearman |
| n2_ampl_dx | p2_lat_sx | -0.218 | 0.114 | 0.329 | Spearman |
| n2_ampl_dx | n2_lat_sx | -0.154 | 0.267 | 0.562 | Spearman |
| n2_ampl_dx | p3_lat_sx | -0.133 | 0.339 | 0.657 | Spearman |
| n2_ampl_dx | p2_n2_sx | 0.429 | 0.001 | 0.010 | Spearman |
| n2_ampl_dx | n2p3_dx | 0.197 | 0.154 | 0.385 | Spearman |
| n2_ampl_dx | n2p3_sx | 0.156 | 0.260 | 0.562 | Spearman |
| p3_ampl_dx | p2_lat_dx | -0.182 | 0.188 | 0.442 | Spearman |
| p3_ampl_dx | n2_lat_dx | -0.046 | 0.743 | 0.865 | Spearman |
| p3_ampl_dx | p3_lat_dx | 0.331 | 0.015 | 0.073 | Spearman |
| p3_ampl_dx | p2_n2_dx | 0.215 | 0.118 | 0.329 | Spearman |
| p3_ampl_dx | p2_lat_sx | -0.289 | 0.034 | 0.138 | Spearman |
| p3_ampl_dx | n2_lat_sx | -0.338 | 0.012 | 0.071 | Spearman |
| p3_ampl_dx | p3_lat_sx | -0.104 | 0.455 | 0.678 | Spearman |
| p3_ampl_dx | p2_n2_sx | 0.083 | 0.549 | 0.784 | Spearman |
| p3_ampl_dx | n2p3_dx | 0.607 | 0.001 | 0.001 | Spearman |
| p3_ampl_dx | n2p3_sx | 0.608 | 0.001 | 0.001 | Spearman |
| n2_ampl_sx | p2_lat_dx | -0.071 | 0.611 | 0.803 | Spearman |
| n2_ampl_sx | n2_lat_dx | 0.060 | 0.666 | 0.808 | Spearman |
| n2_ampl_sx | p3_lat_dx | 0.127 | 0.361 | 0.657 | Spearman |
| n2_ampl_sx | p2_n2_dx | 0.110 | 0.427 | 0.678 | Spearman |
| n2_ampl_sx | p2_lat_sx | -0.257 | 0.061 | 0.222 | Spearman |
| n2_ampl_sx | n2_lat_sx | -0.033 | 0.814 | 0.880 | Spearman |
| n2_ampl_sx | p3_lat_sx | -0.008 | 0.954 | 0.954 | Spearman |
| n2_ampl_sx | p2_n2_sx | 0.576 | 0.001 | 0.001 | Spearman |
| n2_ampl_sx | n2p3_dx | 0.128 | 0.358 | 0.657 | Spearman |
| n2_ampl_sx | n2p3_sx | 0.113 | 0.414 | 0.678 | Spearman |
| p3_ampl_sx | p2_lat_dx | 0.069 | 0.622 | 0.803 | Spearman |
| p3_ampl_sx | n2_lat_dx | 0.071 | 0.613 | 0.803 | Spearman |
| p3_ampl_sx | p3_lat_dx | 0.311 | 0.024 | 0.105 | Spearman |
| p3_ampl_sx | p2_n2_dx | -0.040 | 0.779 | 0.865 | Spearman |
| p3_ampl_sx | p2_lat_sx | -0.063 | 0.656 | 0.808 | Spearman |
| p3_ampl_sx | n2_lat_sx | -0.214 | 0.124 | 0.329 | Spearman |
| p3_ampl_sx | p3_lat_sx | 0.043 | 0.758 | 0.865 | Spearman |
| p3_ampl_sx | p2_n2_sx | -0.029 | 0.839 | 0.883 | Spearman |
| p3_ampl_sx | n2p3_dx | 0.381 | 0.005 | 0.033 | Spearman |
| p3_ampl_sx | n2p3_sx | 0.573 | 0.001 | 0.001 | Spearman |
