## Supplementary material for "Promoting Healthy Ageing through Choral Training: A 9-Month Multidomain Intervention (MultiMusic) Enhances Neural Processing Speed in Community-Dwelling Older Adults": T0_intradomain_FINAL_.docx

### 1. T0 Intra-domain correlations

#### 1.1. Cognitive

| Var_A | Var_B | r | p | q | Method |
| --- | --- | --- | --- | --- | --- |
| mo_ca | miq | 0.476 | 0.001 | 0.001 | Spearman |
| mo_ca | cri_q | 0.650 | 0.001 | 0.001 | Spearman |
| miq | cri_q | 0.477 | 0.001 | 0.001 | Spearman |

#### 1.2. Musical

| Var_A | Var_B | r | p | q | Method |
| --- | --- | --- | --- | --- | --- |
| mdt | mpt | 0.193 | 0.162 | 0.162 | Spearman |
| mdt | rat | 0.400 | 0.003 | 0.008 | Spearman |
| mdt | edt | 0.208 | 0.132 | 0.158 | Spearman |
| mpt | rat | 0.343 | 0.011 | 0.022 | Spearman |
| mpt | edt | 0.232 | 0.091 | 0.136 | Pearson |
| rat | edt | 0.420 | 0.002 | 0.008 | Spearman |

#### 1.3. Audiometric

| Var_A | Var_B | r | p | q | Method |
| --- | --- | --- | --- | --- | --- |
| pta_right (x -1) | pta_left (x -1) | 0.939 | 0.001 | 0.001 | Spearman |
| pta_right (x -1) | srt_right (x -1) | 0.786 | 0.001 | 0.001 | Spearman |
| pta_right (x -1) | srt_left (x -1) | 0.784 | 0.001 | 0.001 | Spearman |
| pta_right (x -1) | matrix_test (x -1) | 0.675 | 0.001 | 0.001 | Spearman |
| pta_right (x -1) | intellection_right | 0.604 | 0.001 | 0.001 | Spearman |
| pta_right (x -1) | intellection_left | 0.616 | 0.001 | 0.001 | Spearman |
| pta_left (x -1) | srt_right (x -1) | 0.815 | 0.001 | 0.001 | Spearman |
| pta_left (x -1) | srt_left (x -1) | 0.830 | 0.001 | 0.001 | Spearman |
| pta_left (x -1) | matrix_test (x -1) | 0.725 | 0.001 | 0.001 | Spearman |
| pta_left (x -1) | intellection_right | 0.597 | 0.001 | 0.001 | Spearman |
| pta_left (x -1) | intellection_left | 0.616 | 0.001 | 0.001 | Spearman |
| srt_right (x -1) | srt_left (x -1) | 0.969 | 0.001 | 0.001 | Spearman |
| srt_right (x -1) | matrix_test (x -1) | 0.645 | 0.001 | 0.001 | Spearman |
| srt_right (x -1) | intellection_right | 0.657 | 0.001 | 0.001 | Spearman |
| srt_right (x -1) | intellection_left | 0.657 | 0.001 | 0.001 | Spearman |
| srt_left (x -1) | matrix_test (x -1) | 0.632 | 0.001 | 0.001 | Spearman |
| srt_left (x -1) | intellection_right | 0.653 | 0.001 | 0.001 | Spearman |
| srt_left (x -1) | intellection_left | 0.685 | 0.001 | 0.001 | Spearman |
| matrix_test (x -1) | intellection_right | 0.518 | 0.001 | 0.001 | Spearman |
| matrix_test (x -1) | intellection_left | 0.533 | 0.001 | 0.001 | Spearman |
| intellection_right | intellection_left | 0.916 | 0.001 | 0.001 | Spearman |

#### 1.4. Neurophysiologic_amplitude

| Var_A | Var_B | r | p | q | Method |
| --- | --- | --- | --- | --- | --- |
| n2_ampl_dx | p3_ampl_dx | 0.448 | 0.001 | 0.002 | Spearman |
| n2_ampl_dx | n2_ampl_sx | 0.325 | 0.016 | 0.016 | Spearman |
| n2_ampl_dx | p3_ampl_sx | 0.341 | 0.013 | 0.015 | Spearman |
| p3_ampl_dx | n2_ampl_sx | 0.351 | 0.009 | 0.014 | Spearman |
| p3_ampl_dx | p3_ampl_sx | 0.720 | 0.001 | 0.001 | Spearman |
| n2_ampl_sx | p3_ampl_sx | 0.423 | 0.002 | 0.003 | Spearman |

#### 1.5. Neurophysiologic_latency

| Var_A | Var_B | r | p | q | Method |
| --- | --- | --- | --- | --- | --- |
| p2_lat_dx | n2_lat_dx | 0.762 | 0.001 | 0.001 | Spearman |
| p2_lat_dx | p3_lat_dx | 0.587 | 0.001 | 0.001 | Spearman |
| p2_lat_dx | p2_n2_dx | -0.618 | 0.001 | 0.001 | Spearman |
| p2_lat_dx | p2_lat_sx | 0.664 | 0.001 | 0.001 | Spearman |
| p2_lat_dx | n2_lat_sx | 0.622 | 0.001 | 0.001 | Spearman |
| p2_lat_dx | p3_lat_sx | 0.640 | 0.001 | 0.001 | Spearman |
| p2_lat_dx | p2_n2_sx | -0.103 | 0.460 | 0.576 | Spearman |
| p2_lat_dx | n2p3_dx | -0.439 | 0.001 | 0.002 | Spearman |
| p2_lat_dx | n2p3_sx | -0.063 | 0.649 | 0.769 | Spearman |
| n2_lat_dx | p3_lat_dx | 0.815 | 0.001 | 0.001 | Spearman |
| n2_lat_dx | p2_n2_dx | -0.181 | 0.191 | 0.268 | Spearman |
| n2_lat_dx | p2_lat_sx | 0.557 | 0.001 | 0.001 | Spearman |
| n2_lat_dx | n2_lat_sx | 0.724 | 0.001 | 0.001 | Spearman |
| n2_lat_dx | p3_lat_sx | 0.775 | 0.001 | 0.001 | Spearman |
| n2_lat_dx | p2_n2_sx | -0.048 | 0.731 | 0.802 | Spearman |
| n2_lat_dx | n2p3_dx | -0.467 | 0.001 | 0.001 | Spearman |
| n2_lat_dx | n2p3_sx | -0.026 | 0.854 | 0.894 | Spearman |
| p3_lat_dx | p2_n2_dx | -0.103 | 0.457 | 0.576 | Spearman |
| p3_lat_dx | p2_lat_sx | 0.301 | 0.027 | 0.049 | Spearman |
| p3_lat_dx | n2_lat_sx | 0.433 | 0.001 | 0.002 | Spearman |
| p3_lat_dx | p3_lat_sx | 0.620 | 0.001 | 0.001 | Spearman |
| p3_lat_dx | p2_n2_sx | -0.038 | 0.787 | 0.843 | Spearman |
| p3_lat_dx | n2p3_dx | 0.004 | 0.974 | 0.974 | Spearman |
| p3_lat_dx | n2p3_sx | 0.289 | 0.034 | 0.059 | Spearman |
| p2_n2_dx | p2_lat_sx | -0.308 | 0.023 | 0.044 | Spearman |
| p2_n2_dx | n2_lat_sx | -0.221 | 0.109 | 0.163 | Spearman |
| p2_n2_dx | p3_lat_sx | -0.273 | 0.046 | 0.077 | Spearman |
| p2_n2_dx | p2_n2_sx | 0.055 | 0.694 | 0.801 | Spearman |
| p2_n2_dx | n2p3_dx | 0.214 | 0.120 | 0.174 | Spearman |
| p2_n2_dx | n2p3_sx | -0.050 | 0.722 | 0.802 | Spearman |
| p2_lat_sx | n2_lat_sx | 0.802 | 0.001 | 0.001 | Spearman |
| p2_lat_sx | p3_lat_sx | 0.689 | 0.001 | 0.001 | Spearman |
| p2_lat_sx | p2_n2_sx | -0.455 | 0.001 | 0.001 | Spearman |
| p2_lat_sx | n2p3_dx | -0.439 | 0.001 | 0.002 | Spearman |
| p2_lat_sx | n2p3_sx | -0.270 | 0.048 | 0.077 | Spearman |
| n2_lat_sx | p3_lat_sx | 0.856 | 0.001 | 0.001 | Spearman |
| n2_lat_sx | p2_n2_sx | -0.174 | 0.209 | 0.286 | Spearman |
| n2_lat_sx | n2p3_dx | -0.529 | 0.001 | 0.001 | Spearman |
| n2_lat_sx | n2p3_sx | -0.334 | 0.013 | 0.026 | Spearman |
| p3_lat_sx | p2_n2_sx | -0.249 | 0.070 | 0.108 | Spearman |
| p3_lat_sx | n2p3_dx | -0.343 | 0.011 | 0.023 | Spearman |
| p3_lat_sx | n2p3_sx | 0.127 | 0.359 | 0.475 | Spearman |
| p2_n2_sx | n2p3_dx | 0.063 | 0.649 | 0.769 | Spearman |
| p2_n2_sx | n2p3_sx | -0.005 | 0.972 | 0.974 | Spearman |
| n2p3_dx | n2p3_sx | 0.515 | 0.001 | 0.001 | Spearman |
