## Supplementary material for "Promoting Healthy Ageing through Choral Training: A 9-Month Multidomain Intervention (MultiMusic) Enhances Neural Processing Speed in Community-Dwelling Older Adults": Zou_T0_pair_test.docx

### 1. Zou tests at T0 for bilateral variables vs specified targets

Targets: Musical (MDT, MPT, EDT, RAT), Cognitive (MoCA, CRI_Q, MIQ), Audiometric (Matrix_test).

| **bilateral** | **X** | | **Y** | | **r_dxY** | | **r_sxY** | | **r_dx_sx** | | **diff_r** | | **abs_diff** | | **Zou 95% CI** | | **q_FDR** |
| --- | --- | --- | --- | --- | --- | --- | --- | --- | --- | --- | --- | --- | --- | --- | --- | --- | --- |
| PTA | PTA_right/left | | MDT | | 0.397 | | 0.425 | | 0.965 | | -0.027 | | 0.027 | | [-0.11, 0.05] | | 0.5000 |
| PTA | PTA_right/left | | MPT | | 0.199 | | 0.214 | | 0.965 | | -0.015 | | 0.015 | | [-0.09, 0.06] | | 0.5000 |
| PTA | PTA_right/left | | EDT | | 0.077 | | 0.119 | | 0.965 | | -0.042 | | 0.042 | | [-0.11, 0.03] | | 0.5000 |
| PTA | PTA_right/left | | RAT | | 0.239 | | 0.244 | | 0.939 | | -0.006 | | 0.006 | | [-0.10, 0.09] | | 0.5000 |
| PTA | PTA_right/left | | MoCa | | 0.389 | | 0.326 | | 0.939 | | 0.062 | | 0.062 | | [-0.03, 0.17] | | 0.5000 |
| PTA | PTA_right/left | | CRI_Q | | 0.145 | | 0.073 | | 0.939 | | 0.072 | | 0.072 | | [-0.02, 0.17] | | 0.5000 |
| PTA | PTA_right/left | | MIQ | | 0.141 | | 0.144 | | 0.939 | | -0.002 | | 0.002 | | [-0.10, 0.09] | | 0.5000 |
| PTA | PTA_right/left | | Matrix_test | | 0.675 | | 0.725 | | 0.939 | | -0.050 | | 0.050 | | [-0.15, 0.03] | | 0.5000 |
| SRT | SRT_right/left | | MDT | | 0.368 | | 0.445 | | 0.969 | | -0.077 | | 0.077 | | [-0.16, -0.01] | | 0.1996 |
| SRT | SRT_right/left | | MPT | | 0.262 | | 0.262 | | 0.969 | | 0.000 | | 0.000 | | [-0.07, 0.07] | | 0.5000 |
| SRT | SRT_right/left | | EDT | | 0.051 | | 0.138 | | 0.969 | | -0.088 | | 0.088 | | [-0.16, -0.02] | | 0.1996 |
| SRT | SRT_right/left | | RAT | | 0.236 | | 0.259 | | 0.969 | | -0.023 | | 0.023 | | [-0.10, 0.05] | | 0.5000 |
| SRT | SRT_right/left | | MoCa | | 0.250 | | 0.284 | | 0.969 | | -0.033 | | 0.033 | | [-0.11, 0.04] | | 0.5000 |
| SRT | SRT_right/left | | CRI_Q | | 0.046 | | 0.016 | | 0.969 | | 0.030 | | 0.030 | | [-0.04, 0.10] | | 0.5000 |
| SRT | SRT_right/left | | MIQ | | 0.101 | | 0.127 | | 0.969 | | -0.026 | | 0.026 | | [-0.09, 0.04] | | 0.5000 |
| SRT | SRT_right/left | | Matrix_test | | 0.645 | | 0.632 | | 0.968 | | 0.013 | | 0.013 | | [-0.06, 0.10] | | 0.5000 |
| Intellection | Intellection_right/left | | MDT | | 0.321 | | 0.351 | | 0.916 | | -0.030 | | 0.030 | | [-0.14, 0.08] | | 0.5000 |
| Intellection | Intellection_right/left | | MPT | | 0.213 | | 0.154 | | 0.916 | | 0.059 | | 0.059 | | [-0.05, 0.17] | | 0.5000 |
| Intellection | Intellection_right/left | | EDT | | 0.038 | | 0.092 | | 0.916 | | -0.055 | | 0.055 | | [-0.16, 0.06] | | 0.5000 |
| Intellection | Intellection_right/left | | RAT | | 0.042 | | 0.075 | | 0.916 | | -0.033 | | 0.033 | | [-0.14, 0.08] | | 0.5000 |
| Intellection | Intellection_right/left | | MoCa | | 0.358 | | 0.346 | | 0.916 | | 0.012 | | 0.012 | | [-0.10, 0.13] | | 0.5000 |
| Intellection | Intellection_right/left | | CRI_Q | | 0.110 | | 0.101 | | 0.916 | | 0.010 | | 0.010 | | [-0.10, 0.12] | | 0.5000 |
| Intellection | Intellection_right/left | | MIQ | | 0.130 | | 0.065 | | 0.916 | | 0.065 | | 0.065 | | [-0.05, 0.17] | | 0.5000 |
| Intellection | Intellection_right/left | | Matrix_test | | 0.518 | | 0.533 | | 0.916 | | -0.015 | | 0.015 | | [-0.13, 0.09] | | 0.5000 |
| P2_lat | P2_lat_ right/left | | MDT | | 0.082 | | -0.042 | | 0.627 | | 0.124 | | 0.124 | | [-0.11, 0.35] | | 0.5000 |
| P2_lat | P2_lat_ right/left | | MPT | | 0.045 | | 0.060 | | 0.627 | | -0.014 | | 0.014 | | [-0.24, 0.22] | | 0.5000 |
| P2_lat | P2_lat_ right/left | | EDT | | 0.309 | | 0.084 | | 0.627 | | 0.225 | | 0.225 | | [-0.00, 0.45] | | 0.5000 |
| P2_lat | P2_lat_ right/left | | RAT | | 0.152 | | 0.151 | | 0.664 | | 0.000 | | 0.000 | | [-0.22, 0.22] | | 0.5000 |
| P2_lat | P2_lat_ right/left | | MoCa | | 0.106 | | 0.232 | | 0.664 | | -0.126 | | 0.126 | | [-0.34, 0.09] | | 0.5000 |
| P2_lat | P2_lat_ right/left | | CRI_Q | | 0.116 | | 0.157 | | 0.664 | | -0.042 | | 0.042 | | [-0.26, 0.18] | | 0.5000 |
| P2_lat | P2_lat_ right/left | | MIQ | | 0.214 | | 0.147 | | 0.664 | | 0.067 | | 0.067 | | [-0.15, 0.28] | | 0.5000 |
| P2_lat | P2_lat_ right/left | | Matrix_test | | 0.097 | | 0.057 | | 0.662 | | 0.040 | | 0.040 | | [-0.18, 0.26] | | 0.5000 |
| N2_lat | N2_lat_ right/left | | MDT | | 0.044 | | -0.074 | | 0.720 | | 0.118 | | 0.118 | | [-0.08, 0.31] | | 0.5000 |
| N2_lat | N2_lat_right/left | | MPT | | 0.059 | | 0.153 | | 0.720 | | -0.094 | | 0.094 | | [-0.29, 0.11] | | 0.5000 |
| N2_lat | N2_lat_ right/left | | EDT | | 0.288 | | 0.099 | | 0.720 | | 0.189 | | 0.189 | | [-0.01, 0.38] | | 0.5000 |
| N2_lat | N2_lat_ right/left | | RAT | | 0.246 | | 0.287 | | 0.724 | | -0.041 | | 0.041 | | [-0.24, 0.15] | | 0.5000 |
| N2_lat | N2_lat_right/left | | MoCa | | 0.210 | | 0.273 | | 0.724 | | -0.063 | | 0.063 | | [-0.26, 0.13] | | 0.5000 |
| N2_lat | N2_lat_ right/left | | CRI_Q | | 0.112 | | 0.115 | | 0.724 | | -0.002 | | 0.002 | | [-0.20, 0.20] | | 0.5000 |
| N2_lat | N2_lat_ right/left | | MIQ | | 0.207 | | 0.196 | | 0.724 | | 0.011 | | 0.011 | | [-0.19, 0.21] | | 0.5000 |
| N2_lat | N2_lat_ right/left | | Matrix_test | | 0.030 | | 0.080 | | 0.724 | | -0.051 | | 0.051 | | [-0.25, 0.15] | | 0.5000 |
| P3_lat | P3_lat_ right/left | | MDT | | 0.081 | | -0.051 | | 0.599 | | 0.132 | | 0.132 | | [-0.11, 0.37] | | 0.5000 |
| P3_lat | P3_lat_ right/left | | MPT | | 0.014 | | 0.003 | | 0.599 | | 0.011 | | 0.011 | | [-0.23, 0.25] | | 0.5000 |
| P3_lat | P3_lat_ right/left | | EDT | | 0.225 | | 0.076 | | 0.599 | | 0.148 | | 0.148 | | [-0.09, 0.38] | | 0.5000 |
| P3_lat | P3_lat_ right/left | RAT | | 0.224 | | 0.209 | | 0.620 | | 0.015 | | 0.015 | | [-0.21, 0.24] | | 0.5000 | |
| P3_lat | P3_lat_ right/left | MoCa | | 0.027 | | 0.159 | | 0.620 | | -0.131 | | 0.131 | | [-0.36, 0.10] | | 0.5000 | |
| P3_lat | P3_lat_ right/left | CRI_Q | | -0.008 | | -0.034 | | 0.620 | | 0.026 | | 0.026 | | [-0.21, 0.26] | | 0.5000 | |
| P3_lat | P3_lat_ right/left | MIQ | | 0.142 | | 0.087 | | 0.620 | | 0.055 | | 0.055 | | [-0.18, 0.29] | | 0.5000 | |
| P3_lat | P3_lat_ right/left | Matrix_test | | -0.142 | | -0.039 | | 0.597 | | -0.103 | | 0.103 | | [-0.34, 0.14] | | 0.5000 | |
| P2_N2 | P2_N2_ right/left | MDT | | -0.003 | | 0.113 | | 0.055 | | -0.116 | | 0.116 | | [-0.48, 0.26] | | 0.5000 | |
| P2_N2 | P2_N2_ right/left | MPT | | 0.008 | | 0.215 | | 0.055 | | -0.207 | | 0.207 | | [-0.56, 0.16] | | 0.5000 | |
| P2_N2 | P2_N2_ right/left | EDT | | -0.071 | | -0.046 | | 0.055 | | -0.025 | | 0.025 | | [-0.39, 0.34] | | 0.5000 | |
| P2_N2 | P2_N2_ right/left | RAT | | -0.001 | | 0.064 | | 0.055 | | -0.065 | | 0.065 | | [-0.43, 0.31] | | 0.5000 | |
| P2_N2 | P2_N2_ right/left | MoCa | | -0.034 | | 0.063 | | 0.055 | | -0.096 | | 0.096 | | [-0.46, 0.28] | | 0.5000 | |
| P2_N2 | P2_N2_ right/left | CRI_Q | | -0.077 | | 0.006 | | 0.055 | | -0.083 | | 0.083 | | [-0.45, 0.29] | | 0.5000 | |
| P2_N2 | P2_N2_ right/left | MIQ | | 0.006 | | 0.005 | | 0.055 | | 0.001 | | 0.001 | | [-0.37, 0.37] | | 0.5000 | |
| P2_N2 | P2_N2_ right/left | Matrix_test | | -0.097 | | 0.023 | | 0.024 | | -0.120 | | 0.120 | | [-0.49, 0.27] | | 0.5000 | |
| N2_ampl | N2_ampl_right/left | MDT | | 0.265 | | 0.045 | | 0.325 | | 0.220 | | 0.220 | | [-0.09, 0.51] | | 0.5000 | |
| N2_ampl | N2_ampl_right/left | MPT | | 0.178 | | 0.195 | | 0.325 | | -0.018 | | 0.018 | | [-0.32, 0.29] | | 0.5000 | |
| N2_ampl | N2_ampl_right/left | EDT | | 0.094 | | 0.086 | | 0.325 | | 0.009 | | 0.009 | | [-0.30, 0.32] | | 0.5000 | |
| N2_ampl | N2_ampl_right/left | RAT | | 0.148 | | 0.291 | | 0.325 | | -0.143 | | 0.143 | | [-0.44, 0.16] | | 0.5000 | |
| N2_ampl | N2_ampl_right/left | MoCa | | 0.189 | | 0.039 | | 0.325 | | 0.150 | | 0.150 | | [-0.16, 0.45] | | 0.5000 | |
| N2_ampl | N2_ampl_right/left | CRI_Q | | 0.211 | | 0.013 | | 0.325 | | 0.197 | | 0.197 | | [-0.12, 0.50] | | 0.5000 | |
| N2_ampl | N2_ampl_right/left | MIQ | | 0.426 | | -0.066 | | 0.325 | | 0.492 | | 0.492 | | [0.19, 0.76] | | 0.3992 | |
| N2_ampl | N2_ampl_right/left | Matrix_test | | -0.133 | | -0.093 | | 0.316 | | -0.040 | | 0.040 | | [-0.35, 0.28] | | 0.5000 | |
| P3_ampl | P3_ampl_right/left | MDT | | 0.070 | | 0.143 | | 0.720 | | -0.073 | | 0.073 | | [-0.27, 0.13] | | 0.5000 | |
| P3_ampl | P3_ampl_ right/left | MPT | | 0.060 | | -0.021 | | 0.720 | | 0.081 | | 0.081 | | [-0.12, 0.28] | | 0.5000 | |
| P3_ampl | P3_ampl_ right/left | EDT | | 0.196 | | 0.193 | | 0.720 | | 0.003 | | 0.003 | | [-0.20, 0.20] | | 0.5000 | |
| P3_ampl | P3_ampl_ right/left | RAT | | 0.140 | | 0.224 | | 0.720 | | -0.084 | | 0.084 | | [-0.28, 0.12] | | 0.5000 | |
| P3_ampl | P3_ampl_ right/left | MoCa | | -0.099 | | -0.085 | | 0.720 | | -0.014 | | 0.014 | | [-0.22, 0.19] | | 0.5000 | |
| P3_ampl | P3_ampl_ right/left | CRI_Q | | -0.038 | | -0.013 | | 0.720 | | -0.025 | | 0.025 | | [-0.23, 0.18] | | 0.5000 | |
| P3_ampl | P3_ampl_ right/left | MIQ | | 0.080 | | 0.021 | | 0.720 | | 0.059 | | 0.059 | | [-0.15, 0.26] | | 0.5000 | |
| P3_ampl | P3_ampl_ right/left | Matrix_test | | -0.212 | | -0.156 | | 0.712 | | -0.055 | | 0.055 | | [-0.26, 0.15] | | 0.5000 | |
| n2p3 | n2p3_ right/left | MDT | | 0.047 | | 0.063 | | 0.515 | | -0.016 | | 0.016 | | [-0.28, 0.25] | | 0.5000 | |
| n2p3 | n2p3_ right/left | MPT | | -0.067 | | -0.250 | | 0.515 | | 0.183 | | 0.183 | | [-0.08, 0.44] | | 0.5000 | |
| n2p3 | n2p3_ right/left | EDT | | -0.017 | | -0.018 | | 0.515 | | 0.002 | | 0.002 | | [-0.26, 0.27] | | 0.5000 | |
| n2p3 | n2p3_ right/left | RAT | | -0.186 | | -0.101 | | 0.515 | | -0.085 | | 0.085 | | [-0.34, 0.18] | | 0.5000 | |
| n2p3 | n2p3_ right/left | MoCa | | -0.328 | | -0.188 | | 0.515 | | -0.141 | | 0.141 | | [-0.39, 0.12] | | 0.5000 | |
| n2p3 | n2p3_ right/left | CRI_Q | | -0.299 | | -0.235 | | 0.515 | | -0.064 | | 0.064 | | [-0.32, 0.19] | | 0.5000 | |
| n2p3 | n2p3_ right/left | MIQ | | -0.189 | | -0.119 | | 0.515 | | -0.071 | | 0.071 | | [-0.33, 0.19] | | 0.5000 | |
| n2p3 | n2p3_ right/left | Matrix_test | | -0.194 | | -0.174 | | 0.476 | | -0.020 | | 0.020 | | [-0.29, 0.26] | | 0.5000 | |
