## Supplementary material for "Promoting Healthy Ageing through Choral Training: A 9-Month Multidomain Intervention (MultiMusic) Enhances Neural Processing Speed in Community-Dwelling Older Adults": Latency_effect_analysis.docx

To formally compare the strength of longitudinal effects across neurophysiological outcomes, we fitted a joint mixed-effects model including Outcome (N2 latency vs. N2–P3 interpeak latency) as an additional factor and tested the three-way interaction Outcome × Time × Group. This interaction was significant, F(2, 373.6) = 9.73, p < .001, indicating that the magnitude of the Time × Group effect differed between the two latency measures. To quantify this difference, we subsequently estimated separate mixed-effects models for N2 and N2–P3 latencies using an identical fixed-effects structure (Time × Group, with PC1_P300_latency_T0 as covariate and Side included as a factor). Effect size estimates (partial η² for the Time × Group interaction) showed that the interaction accounted for approximately 15% of the variance in N2–P3 latency (η²p ≈ .15), compared with about 5% in N2 latency (η²p ≈ .05). Follow-up hemisphere-specific models further demonstrated that the Time × Group effect on N2–P3 latency was robust and comparable across both hemispheres (η²p ≈ .23–.24), whereas the corresponding effect for N2 latency was weaker and less spatially consistent (η²p ≈ .08 on the right; ≈ .02 on the left). (**Figure 5a** and **5b**).

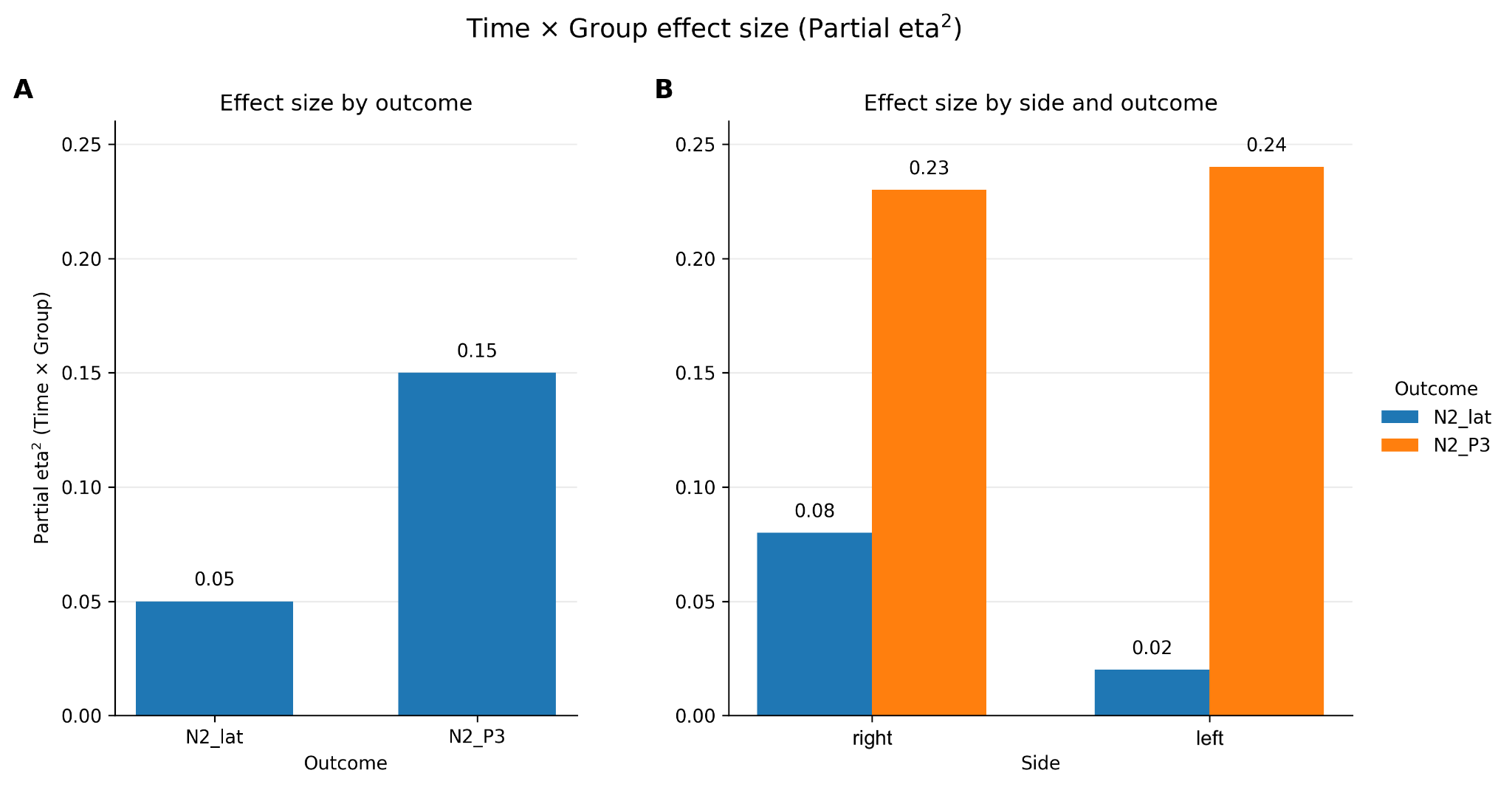

*Figure 5. Time × Group interaction effect sizes (partial η²) for N2 latency and N2–P3 inter-peak latency (IPL). Panel A shows the overall Time × Group interaction effect size for each outcome, indicating a larger effect for the N2–P3 IPL compared to N2 latency alone. Panel B displays the same interaction effect sizes separately for right (dx) and left (sx) hemispheric recordings. The interaction effect for N2–P3 is consistently large across both sides, whereas the effect for N2 latency is smaller and more variable. Values above bars indicate partial η².*
