## Supplementary material for "Promoting Healthy Ageing through Choral Training: A 9-Month Multidomain Intervention (MultiMusic) Enhances Neural Processing Speed in Community-Dwelling Older Adults": R_outputs_hearing.docx

1) Matrix

```
Linear mixed model fit by maximum likelihood . t-tests use Satterthwaite's
  method [lmerModLmerTest]
Formula: Matrix ~ PC1_Hearing_T0 + (1 | ID)

      AIC       BIC    logLik -2*log(L)  df.resid
    259.9     270.5    -126.0     251.9       100

Scaled residuals:
     Min       1Q   Median       3Q      Max
-2.38955 -0.26421  0.01695  0.25086  2.39951

Random effects:
 Groups   Name        Variance Std.Dev.
 ID       (Intercept) 1.5730   1.2542
 Residual             0.1328   0.3644
Number of obs: 104, groups:  ID, 52

Fixed effects:
                 Estimate Std. Error         df t value Pr(>|t|)
(Intercept)    -2.2524296  0.1830320 51.9999917 -12.306  < 2e-16 ***
PC1_Hearing_T0  0.0015083  0.0002077 51.9999916   7.263 1.89e-09 ***
---
Signif. codes:  0 ‘***’ 0.001 ‘**’ 0.01 ‘*’ 0.05 ‘.’ 0.1 ‘ ’ 1

Correlation of Fixed Effects:
            (Intr)
PC1_Hrng_T0 -0.243
```

2) PTA

```
Linear mixed model fit by maximum likelihood . t-tests use Satterthwaite's
  method [lmerModLmerTest]
Formula: PTA ~ PC1_Hearing_T0 + Time + Side + (1 | ID)
   Data: pta_long

      AIC       BIC    logLik -2*log(L)  df.resid
   1114.1    1134.3    -551.0    1102.1       210

Scaled residuals:
    Min      1Q  Median      3Q     Max
-2.2124 -0.5840  0.0494  0.4984  3.8742

Random effects:
 Groups   Name        Variance Std.Dev.
 ID       (Intercept) 29.509   5.432
 Residual              4.126   2.031
Number of obs: 216, groups:  ID, 54

Fixed effects:
                Estimate Std. Error        df t value Pr(>|t|)
(Intercept)    2.399e+01  7.986e-01 6.102e+01  30.039  < 2e-16 ***
PC1_Hearing_T0 8.863e-03  8.953e-04 5.400e+01   9.899 9.78e-14 ***
TimeT1         1.544e+00  2.764e-01 1.620e+02   5.585 9.67e-08 ***
SideRight      9.877e-01  2.764e-01 1.620e+02   3.573 0.000465 ***
---
Signif. codes:  0 ‘***’ 0.001 ‘**’ 0.01 ‘*’ 0.05 ‘.’ 0.1 ‘ ’ 1

Correlation of Fixed Effects:
            (Intr) PC1_H_ TimeT1
PC1_Hrng_T0 -0.231
TimeT1      -0.173  0.000
SideRight   -0.173  0.000  0.000


Type III Analysis of Variance Table with Satterthwaite's method
               Sum Sq Mean Sq NumDF DenDF F value    Pr(>F)
PC1_Hearing_T0 404.32  404.32     1    54  97.990 9.784e-14 ***
Time           128.70  128.70     1   162  31.191 9.671e-08 ***
Side            52.68   52.68     1   162  12.767  0.000465 ***
---
Signif. codes:  0 ‘***’ 0.001 ‘**’ 0.01 ‘*’ 0.05 ‘.’ 0.1 ‘ ’ 1
```

3) SRT

```
Linear mixed model fit by maximum likelihood . t-tests use Satterthwaite's
  method [lmerModLmerTest]
Formula: SRT ~ PC1_Hearing_T0 + Time + (1 | ID)

      AIC       BIC    logLik -2*log(L)  df.resid
   1164.7    1181.6    -577.4    1154.7       211

Scaled residuals:
    Min      1Q  Median      3Q     Max
-3.6350 -0.3654  0.0180  0.3104  4.1278

Random effects:
 Groups   Name        Variance Std.Dev.
 ID       (Intercept) 25.760   5.075
 Residual              5.932   2.436
Number of obs: 216, groups:  ID, 54

Fixed effects:
                Estimate Std. Error        df t value Pr(>|t|)
(Intercept)    3.077e+01  7.499e-01 5.964e+01  41.038  < 2e-16 ***
PC1_Hearing_T0 7.979e-03  8.456e-04 5.400e+01   9.436 5.13e-13 ***
TimeT1         1.093e+00  3.314e-01 1.620e+02   3.296   0.0012 **
---
Signif. codes:  0 ‘***’ 0.001 ‘**’ 0.01 ‘*’ 0.05 ‘.’ 0.1 ‘ ’ 1

Correlation of Fixed Effects:
            (Intr) PC1_H_
PC1_Hrng_T0 -0.232
TimeT1      -0.221  0.000
ANOVA tipo III (Satterthwaite):


Type III Analysis of Variance Table with Satterthwaite's method
               Sum Sq Mean Sq NumDF DenDF F value   Pr(>F)
PC1_Hearing_T0 528.16  528.16     1    54  89.031 5.13e-13 ***
Time            64.46   64.46     1   162  10.866 0.001203 **
---
Signif. codes:  0 ‘***’ 0.001 ‘**’ 0.01 ‘*’ 0.05 ‘.’ 0.1 ‘ ’ 1
```

4) Intellection

```
Linear mixed model fit by maximum likelihood . t-tests use Satterthwaite's
  method [lmerModLmerTest]
Formula: Intellection ~ PC1_Hearing_T0 + (1 | ID)

      AIC       BIC    logLik -2*log(L)  df.resid
   1402.0    1415.5    -697.0    1394.0       212

Scaled residuals:
    Min      1Q  Median      3Q     Max
-4.9578  0.0797  0.0798  0.0800  5.7610

Random effects:
 Groups   Name        Variance Std.Dev.
 ID       (Intercept) 40.98    6.401
 Residual             21.76    4.665
Number of obs: 216, groups:  ID, 54

Fixed effects:
                Estimate Std. Error        df t value Pr(>|t|)
(Intercept)    96.815665   0.954595 53.999981  101.42   <2e-16 ***
PC1_Hearing_T0 -0.013984   0.001104 53.999981  -12.67   <2e-16 ***
---
Signif. codes:  0 ‘***’ 0.001 ‘**’ 0.01 ‘*’ 0.05 ‘.’ 0.1 ‘ ’ 1

Correlation of Fixed Effects:
            (Intr)
PC1_Hrng_T0 -0.238
```
