## Supplementary material for "Promoting Healthy Ageing through Choral Training: A 9-Month Multidomain Intervention (MultiMusic) Enhances Neural Processing Speed in Community-Dwelling Older Adults": R_outputs_cognitive.docx

1) MIQ

```
Linear mixed model fit by maximum likelihood . t-tests use Satterthwaite's
  method [lmerModLmerTest]
Formula: MIQ ~ MIQ_T0 + Time * Groups + (1 | ID)

      AIC       BIC    logLik -2*log(L)  df.resid
    236.2     260.3    -109.1     218.2        99

Scaled residuals:
    Min      1Q  Median      3Q     Max
-2.5105 -0.6015 -0.1127  0.4810  2.9908

Random effects:
 Groups   Name        Variance Std.Dev.
 ID       (Intercept) 0.0000   0.0000
 Residual             0.4415   0.6645
Number of obs: 108, groups:  ID, 54

Fixed effects:
                              Estimate Std. Error        df t value Pr(>|t|)
(Intercept)                   -0.90317    0.22896 108.00000  -3.945 0.000142
MIQ_T0                         0.71634    0.05471 108.00000  13.093  < 2e-16
TimeT1                         0.42415    0.21012 108.00000   2.019 0.046009
GroupsControl_Active           0.28407    0.22272 108.00000   1.275 0.204896
GroupsControl_Passive          0.23755    0.22753 108.00000   1.044 0.298805
TimeT1:GroupsControl_Active   -0.80065    0.30530 108.00000  -2.622 0.009990
TimeT1:GroupsControl_Passive  -0.06696    0.31518 108.00000  -0.212 0.832152

(Intercept)                  ***
MIQ_T0                       ***
TimeT1                       *
GroupsControl_Active
GroupsControl_Passive
TimeT1:GroupsControl_Active  **
TimeT1:GroupsControl_Passive
---
Signif. codes:  0 ‘***’ 0.001 ‘**’ 0.01 ‘*’ 0.05 ‘.’ 0.1 ‘ ’ 1

Correlation of Fixed Effects:
            (Intr) MIQ_T0 TimeT1 GrpC_A GrpC_P TT1:GC_A
MIQ_T0       0.761
TimeT1      -0.459  0.000
GrpsCntrl_A -0.620 -0.246  0.472
GrpsCntrl_P -0.577 -0.201  0.462  0.485
TmT1:GrpC_A  0.316  0.000 -0.688 -0.685 -0.318
TmT1:GrpC_P  0.306  0.000 -0.667 -0.314 -0.693  0.459

2. ANOVA di tipo III (Satterthwaite)

Summary
Type III Analysis of Variance Table with Satterthwaite's method
            Sum Sq Mean Sq NumDF DenDF  F value  Pr(>F)
MIQ_T0      75.690  75.690     1   108 171.4322 < 2e-16 ***
Time         0.488   0.488     1   108   1.1044 0.29564
Groups       1.768   0.884     2   108   2.0024 0.13999
Time:Groups  3.586   1.793     2   108   4.0605 0.01994 *
---
Signif. codes:  0 ‘***’ 0.001 ‘**’ 0.01 ‘*’ 0.05 ‘.’ 0.1 ‘ ’ 1
3. EMMEANS Time × Groups

Summary
 Time Groups          emmean    SE  df lower.CL upper.CL
 T0   Choir            -2.77 0.157 115    -3.08    -2.46
 T1   Choir            -2.34 0.157 115    -2.65    -2.03
 T0   Control_Active   -2.48 0.164 115    -2.81    -2.16
 T1   Control_Active   -2.86 0.164 115    -3.18    -2.54
 T0   Control_Passive  -2.53 0.172 115    -2.87    -2.19
 T1   Control_Passive  -2.17 0.172 115    -2.51    -1.83

Degrees-of-freedom method: kenward-roger
Confidence level used: 0.95
4. Contrasti post-hoc
4.1. Time (T1 vs T0) entro ciascun gruppo (Bonferroni)

Summary
Groups = Choir:
 contrast estimate    SE   df t.ratio p.value
 T1 - T0     0.424 0.217 58.3   1.952  0.0557

Groups = Control_Active:
 contrast estimate    SE   df t.ratio p.value
 T1 - T0    -0.377 0.229 58.3  -1.644  0.1056

Groups = Control_Passive:
 contrast estimate    SE   df t.ratio p.value
 T1 - T0     0.357 0.243 58.3   1.470  0.1468

Degrees-of-freedom method: kenward-roger
4.2. Groups entro ciascun Time (Bonferroni)

Summary
Time = T0:
 contrast                         estimate    SE  df t.ratio p.value
 Control_Active - Choir             0.2841 0.230 115   1.233  0.6598
 Control_Passive - Choir            0.2375 0.235 115   1.010  0.9444
 Control_Passive - Control_Active  -0.0465 0.236 115  -0.197  1.0000

Time = T1:
 contrast                         estimate    SE  df t.ratio p.value
 Control_Active - Choir            -0.5166 0.230 115  -2.243  0.0805
 Control_Passive - Choir            0.1706 0.235 115   0.725  1.0000
 Control_Passive - Control_Active   0.6872 0.236 115   2.908  0.0131

Degrees-of-freedom method: kenward-roger
P value adjustment: bonferroni method for 3 tests
```

2) MoCa

```
Linear mixed model fit by maximum likelihood . t-tests use Satterthwaite's
  method [lmerModLmerTest]
Formula: MoCa ~ PC1_Cognitive_T0 + Time + Groups + (1 | ID)

      AIC       BIC    logLik -2*log(L)  df.resid
    506.1     524.9    -246.1     492.1       101

Scaled residuals:
     Min       1Q   Median       3Q      Max
-2.47505 -0.58031  0.00731  0.48462  2.03716

Random effects:
 Groups   Name        Variance Std.Dev.
 ID       (Intercept) 2.133    1.461
 Residual             3.839    1.959
Number of obs: 108, groups:  ID, 54

Fixed effects:
                        Estimate Std. Error         df t value Pr(>|t|)
(Intercept)           21.8649320  0.5170053 70.1837930  42.292  < 2e-16 ***
PC1_Cognitive_T0      -0.0033005  0.0003455 53.9999998  -9.552 3.38e-13 ***
TimeT1                 2.6296296  0.3770657 54.0000002   6.974 4.54e-09 ***
GroupsControl_Active  -0.7682892  0.7291972 53.9999998  -1.054    0.297
GroupsControl_Passive -0.3811706  0.7387428 53.9999998  -0.516    0.608
---
Signif. codes:  0 ‘***’ 0.001 ‘**’ 0.01 ‘*’ 0.05 ‘.’ 0.1 ‘ ’ 1

Correlation of Fixed Effects:
            (Intr) PC1_C_ TimeT1 GrpC_A
PC1_Cgnt_T0 -0.330
TimeT1      -0.365  0.000
GrpsCntrl_A -0.683  0.442  0.000
GrpsCntrl_P -0.673  0.450  0.000  0.650

Analysis of Deviance Table (Type III Wald F tests with Kenward-Roger df)

Response: MoCa
                         F Df Df.res    Pr(>F)
(Intercept)      1668.6446  1 64.247 < 2.2e-16 ***
PC1_Cognitive_T0   84.4743  1 50.000 2.539e-12 ***
Time               47.7350  1 53.000 6.318e-09 ***
Groups              0.5172  2 50.000    0.5994
---
Signif. codes:  0 ‘***’ 0.001 ‘**’ 0.01 ‘*’ 0.05 ‘.’ 0.1 ‘ ’ 1
```
