## Supplementary material for "Promoting Healthy Ageing through Choral Training: A 9-Month Multidomain Intervention (MultiMusic) Enhances Neural Processing Speed in Community-Dwelling Older Adults": flourishing_contrasts.docx

```
1) Final mixed model with groups

Linear mixed model fit by maximum likelihood . t-tests use Satterthwaite's
  method [lmerModLmerTest]
Formula: Flourishing ~ Flourishing_T0 + (1 | ID)

      AIC       BIC    logLik -2*log(L)  df.resid
    588.7     599.2    -290.3     580.7        98

Scaled residuals:
    Min      1Q  Median      3Q     Max
-4.2565 -0.3450  0.0540  0.4101  2.4024

Random effects:
 Groups   Name        Variance Std.Dev.
 ID       (Intercept)  0.00    0.000
 Residual             17.38    4.169
Number of obs: 102, groups:  ID, 51

Fixed effects:
               Estimate Std. Error       df t value Pr(>|t|)
(Intercept)     11.3661     3.5337 102.0000   3.217  0.00174 **
Flourishing_T0   0.7579     0.0747 102.0000  10.145  < 2e-16 ***
---
Signif. codes:  0 ‘***’ 0.001 ‘**’ 0.01 ‘*’ 0.05 ‘.’ 0.1 ‘ ’ 1

Correlation of Fixed Effects:
            (Intr)
Florshng_T0 -0.993


2) Mixed model with Time * Groups interaction

Linear mixed model fit by maximum likelihood . t-tests use Satterthwaite's
  method [lmerModLmerTest]
Formula: Flourishing ~ Time * Groups + Flourishing_T0 + (1 | ID)

      AIC       BIC    logLik -2*log(L)  df.resid
    592.2     615.9    -287.1     574.2        93

Scaled residuals:
    Min      1Q  Median      3Q     Max
-3.8232 -0.3379  0.0500  0.4079  2.6146

Random effects:
 Groups   Name        Variance Std.Dev.
 ID       (Intercept)  0.00    0.000
 Residual             16.31    4.039
Number of obs: 102, groups:  ID, 51

Fixed effects:
                              Estimate Std. Error        df t value Pr(>|t|)
(Intercept)                   12.66758    3.66119 102.00000   3.460  0.00079
TimeT1                         0.94737    1.31031 102.00000   0.723  0.47133
GroupsControl_Active          -0.49324    1.33553 102.00000  -0.369  0.71265
GroupsControl_Passive         -0.54329    1.43059 102.00000  -0.380  0.70491
Flourishing_T0                 0.73724    0.07347 102.00000  10.035  < 2e-16
TimeT1:GroupsControl_Active   -0.39181    1.87862 102.00000  -0.209  0.83520
TimeT1:GroupsControl_Passive  -3.01880    2.01172 102.00000  -1.501  0.13655

(Intercept)                  ***
TimeT1
GroupsControl_Active
GroupsControl_Passive
Flourishing_T0               ***
TimeT1:GroupsControl_Active
TimeT1:GroupsControl_Passive
---
Signif. codes:  0 ‘***’ 0.001 ‘**’ 0.01 ‘*’ 0.05 ‘.’ 0.1 ‘ ’ 1

Correlation of Fixed Effects:
            (Intr) TimeT1 GrpC_A GrpC_P Flr_T0 TT1:GC_A
TimeT1      -0.179
GrpsCntrl_A -0.275  0.491
GrpsCntrl_P -0.267  0.458  0.460
Florshng_T0 -0.967  0.000  0.103  0.106
TmT1:GrpC_A  0.125 -0.697 -0.703 -0.319  0.000
TmT1:GrpC_P  0.117 -0.651 -0.320 -0.703  0.000  0.454

Type III Analysis of Variance Table with Satterthwaite's method
                Sum Sq Mean Sq NumDF DenDF  F value Pr(>F)
Time              0.90    0.90     1   102   0.0552 0.8148
Groups           67.37   33.69     2   102   2.0653 0.1321
Flourishing_T0 1642.41 1642.41     1   102 100.6947 <2e-16 ***
Time:Groups      41.33   20.67     2   102   1.2670 0.2861
---
Signif. codes:  0 ‘***’ 0.001 ‘**’ 0.01 ‘*’ 0.05 ‘.’ 0.1 ‘ ’ 1

emmeans Time * Groups

Time Groups          emmean    SE  df lower.CL upper.CL
 T0   Choir             47.3 0.965 109     45.4     49.2
 T1   Choir             48.3 0.965 109     46.3     50.2
 T0   Control_Active    46.8 0.988 110     44.9     48.8
 T1   Control_Active    47.4 0.988 110     45.4     49.3
 T0   Control_Passive   46.8 1.120 110     44.5     49.0
 T1   Control_Passive   44.7 1.120 110     42.5     46.9

Degrees-of-freedom method: kenward-roger
Confidence level used: 0.95

contrast(flour_emm_interact,
         interaction = "pairwise",
         by = "Time",
         adjust = "bonferroni")

Time = T0:
 Groups_pairwise                  estimate   SE  df t.ratio p.value
 Choir - Control_Active              0.493 1.38 109   0.356  1.0000
 Choir - Control_Passive             0.543 1.48 109   0.367  1.0000
 Control_Active - Control_Passive    0.050 1.49 110   0.034  1.0000

Time = T1:
 Groups_pairwise                  estimate   SE  df t.ratio p.value
 Choir - Control_Active              0.885 1.38 109   0.640  1.0000
 Choir - Control_Passive             3.562 1.48 109   2.403  0.0538
 Control_Active - Control_Passive    2.677 1.49 110   1.795  0.2262


3) Final mixed model with Activities_level

Linear mixed model fit by maximum likelihood . t-tests use Satterthwaite's
  method [lmerModLmerTest]
Formula: Flourishing ~ activities_level + Flourishing_T0 + (1 | ID)

      AIC       BIC    logLik -2*log(L)  df.resid
    586.3     599.5    -288.2     576.3        97

Scaled residuals:
    Min      1Q  Median      3Q     Max
-4.1545 -0.3374  0.0674  0.3823  2.5179

Random effects:
 Groups   Name        Variance Std.Dev.
 ID       (Intercept)  0.00    0.000
 Residual             16.65    4.081
Number of obs: 102, groups:  ID, 51

Fixed effects:
                     Estimate Std. Error        df t value Pr(>|t|)
(Intercept)          13.00571    3.54597 102.00000   3.668 0.000391 ***
activities_levelLow  -1.71706    0.81580 102.00000  -2.105 0.037770 *
Flourishing_T0        0.79052    0.07607 102.00000  10.391 2.05e-17 ***
---
Signif. codes:  0 ‘***’ 0.001 ‘**’ 0.01 ‘*’ 0.05 ‘.’ 0.1 ‘ ’ 1

Type III Analysis of Variance Table with Satterthwaite's method
                  Sum Sq Mean Sq NumDF DenDF F value  Pr(>F)
activities_level   73.77   73.77     1   102    4.43 0.03777 *
Flourishing_T0   1703.35 1703.35     1   102  102.28 < 2e-16 ***
---
Signif. codes:  0 ‘***’ 0.001 ‘**’ 0.01 ‘*’ 0.05 ‘.’ 0.1 ‘ ’ 1


4) Mixed model with Time * Activities_level interaction

Linear mixed model fit by maximum likelihood . t-tests use Satterthwaite's
  method [lmerModLmerTest]
Formula: Flourishing ~ Time * activities_level + Flourishing_T0 + (1 |      ID)

      AIC       BIC    logLik -2*log(L)  df.resid
    587.1     605.5    -286.6     573.1        95

Scaled residuals:
    Min      1Q  Median      3Q     Max
-4.0559 -0.3479  0.0320  0.3534  2.7223

Random effects:
 Groups   Name        Variance Std.Dev.
 ID       (Intercept)  0.00    0.000
 Residual             16.14    4.017
Number of obs: 102, groups:  ID, 51

Fixed effects:
                            Estimate Std. Error        df t value Pr(>|t|)
(Intercept)                 12.22310    3.54051 102.00000   3.452  0.00081 ***
TimeT1                      4.98291    1.86023 102.00000   2.679  0.00863 **
activities_levelLow         9.00477    4.34958 102.00000   2.070  0.04109 *
Flourishing_T0              0.76824    0.07485 102.00000  10.277 3.30e-17 ***
TimeT1:activities_levelLow -3.61337    2.00200 102.00000  -1.805  0.07394 .
---
Signif. codes:  0 ‘***’ 0.001 ‘**’ 0.01 ‘*’ 0.05 ‘.’ 0.1 ‘ ’ 1

Type III Analysis of Variance Table with Satterthwaite's method
                       Sum Sq Mean Sq NumDF DenDF  F value  Pr(>F)
Time                     0.38    0.38     1   102   0.0233 0.87910
activities_level        73.77   73.77     1   102   4.5716 0.03489 *
Flourishing_T0        1703.35 1703.35     1   102 105.5529 < 2e-16 ***
Time:activities_level   52.61   52.61     1   102   3.2602 0.07393 .
---
Signif. codes:  0 ‘***’ 0.001 ‘**’ 0.01 ‘*’ 0.05 ‘.’ 0.1 ‘ ’ 1

EMMEANS Time * activities_level

 Time activities_level emmean    SE  df lower.CL upper.CL
 T0   High               47.1 0.860 107     45.4     48.8
 T1   High               48.7 0.860 107     47.0     50.4
 T0   Low                46.9 0.779 107     45.3     48.4
 T1   Low                45.5 0.779 107     44.0     47.1

Degrees-of-freedom method: kenward-roger
Confidence level used: 0.95

Contrasts: T0 vs T1 Within Groups (Bonferroni)

activities_level = High:
 Time_pairwise estimate   SE   df t.ratio p.value
 T0 - T1          -1.57 1.21 54.2  -1.289  0.2030

activities_level = Low:
 Time_pairwise estimate   SE   df t.ratio p.value
 T0 - T1           1.32 1.10 54.2   1.200  0.2353

Degrees-of-freedom method: kenward-roger

Contrasts: differences Within Time (Bonferroni)
Time = T0:
 activities_level_pairwise estimate   SE  df t.ratio p.value
 High - Low                   0.274 1.16 107   0.236  0.8142

Time = T1:
 activities_level_pairwise estimate   SE  df t.ratio p.value
 High - Low                   3.160 1.16 107   2.720  0.0076

Degrees-of-freedom method: kenward-roger
```
