## Supplementary material for "Promoting Healthy Ageing through Choral Training: A 9-Month Multidomain Intervention (MultiMusic) Enhances Neural Processing Speed in Community-Dwelling Older Adults": full_outputs.docx

```
===== MDT =====
Final model formula: mdt ~ 1 + (1 | id)

Type III ANOVA (final model):
Analysis of Deviance Table (Type III Wald chisquare tests)

Response: mdt
             Chisq Df Pr(>Chisq)
(Intercept) 94.741  1  < 2.2e-16 ***
---
Signif. codes:  0 ‘***’ 0.001 ‘**’ 0.01 ‘*’ 0.05 ‘.’ 0.1 ‘ ’ 1

Type III ANOVA (richest significant step model):
Analysis of Deviance Table (Type III Wald chisquare tests)

Response: mdt
                 Chisq Df Pr(>Chisq)
(Intercept)    29.9824  1  4.360e-08 ***
time            0.0072  1    0.93241
groups          5.6051  2    0.06066 .
pc1_musical_t0 53.3335  1  2.815e-13 ***
time:groups     1.3301  2    0.51426
---
Signif. codes:  0 ‘***’ 0.001 ‘**’ 0.01 ‘*’ 0.05 ‘.’ 0.1 ‘ ’ 1

===== MPT =====
Final model formula: mpt ~ 1 + (1 | id)

Type III ANOVA (final model):
Analysis of Deviance Table (Type III Wald chisquare tests)

Response: mpt
             Chisq Df Pr(>Chisq)
(Intercept) 37.755  1  8.021e-10 ***
---
Signif. codes:  0 ‘***’ 0.001 ‘**’ 0.01 ‘*’ 0.05 ‘.’ 0.1 ‘ ’ 1

Type III ANOVA (richest significant step model):
Analysis of Deviance Table (Type III Wald chisquare tests)

Response: mpt
                 Chisq Df Pr(>Chisq)
(Intercept)    13.4741  1  0.0002419 ***
time            0.7589  1  0.3836851
groups          0.3846  2  0.8250504
pc1_musical_t0 22.5857  1   2.01e-06 ***
time:groups     4.4199  2  0.1097066
---
Signif. codes:  0 ‘***’ 0.001 ‘**’ 0.01 ‘*’ 0.05 ‘.’ 0.1 ‘ ’ 1


Final model formula: edt ~ 1 + (1 | id)

Type III ANOVA (final model):
Analysis of Deviance Table (Type III Wald chisquare tests)

Response: edt
             Chisq Df Pr(>Chisq)
(Intercept) 1.1049  1     0.2932

Type III ANOVA (richest significant step model):
Analysis of Deviance Table (Type III Wald chisquare tests)

Response: edt
                 Chisq Df Pr(>Chisq)
(Intercept)     0.0006  1     0.9811
time            0.5485  1     0.4589
groups          0.4314  2     0.8060
pc1_musical_t0 26.6365  1  2.456e-07 ***
time:groups     0.9803  2     0.6125
---
Signif. codes:  0 ‘***’ 0.001 ‘**’ 0.01 ‘*’ 0.05 ‘.’ 0.1 ‘ ’ 1

===== RAT =====
Final model formula: rat ~ 1 + (1 | id)

Type III ANOVA (final model):
Analysis of Deviance Table (Type III Wald chisquare tests)

Response: rat
             Chisq Df Pr(>Chisq)
(Intercept) 95.833  1  < 2.2e-16 ***
---
Signif. codes:  0 ‘***’ 0.001 ‘**’ 0.01 ‘*’ 0.05 ‘.’ 0.1 ‘ ’ 1

Type III ANOVA (richest significant step model):
Analysis of Deviance Table (Type III Wald chisquare tests)

Response: rat
                  Chisq Df Pr(>Chisq)
(Intercept)    110.2422  1  < 2.2e-16 ***
time             8.4052  1   0.003741 **
groups           7.1874  2   0.027496 *
pc1_musical_t0  82.9560  1  < 2.2e-16 ***
time:groups      4.9692  2   0.083361 .
---
Signif. codes:  0 ‘***’ 0.001 ‘**’ 0.01 ‘*’ 0.05 ‘.’ 0.1 ‘ ’ 1
```
