## Supplementary material for "Promoting Healthy Ageing through Choral Training: A 9-Month Multidomain Intervention (MultiMusic) Enhances Neural Processing Speed in Community-Dwelling Older Adults": R_outputs_musical.docx

1) MDT

```
Linear mixed model fit by maximum likelihood . t-tests use Satterthwaite's
  method [lmerModLmerTest]
Formula: MDT ~ PC1_Musical_T0 + (1 | ID)

      AIC       BIC    logLik -2*log(L)  df.resid
    270.8     281.5    -131.4     262.8       104

Scaled residuals:
     Min       1Q   Median       3Q      Max
-2.64288 -0.63053  0.04434  0.61938  1.77313

Random effects:
 Groups   Name        Variance Std.Dev.
 ID       (Intercept) 0.2489   0.4989
 Residual             0.4632   0.6806
Number of obs: 108, groups:  ID, 54

Fixed effects:
                 Estimate Std. Error         df t value Pr(>|t|)
(Intercept)    -1.2175561  0.0944138 53.9999981 -12.896  < 2e-16 ***
PC1_Musical_T0 -0.0006479  0.0001061 53.9999981  -6.106 1.15e-07 ***
---
Signif. codes:  0 ‘***’ 0.001 ‘**’ 0.01 ‘*’ 0.05 ‘.’ 0.1 ‘ ’ 1

Correlation of Fixed Effects:
            (Intr)
PC1_Mscl_T0 0.041
```

2) MPT

```
Linear mixed model fit by maximum likelihood . t-tests use Satterthwaite's
  method [lmerModLmerTest]
Formula: MPT ~ PC1_Musical_T0 + (1 | ID)

      AIC       BIC    logLik -2*log(L)  df.resid
    288.4     299.1    -140.2     280.4       104

Scaled residuals:
    Min      1Q  Median      3Q     Max
-4.1661 -0.4438  0.1218  0.5792  1.5839

Random effects:
 Groups   Name        Variance Std.Dev.
 ID       (Intercept) 0.1914   0.4375
 Residual             0.6167   0.7853
Number of obs: 108, groups:  ID, 54

Fixed effects:
                 Estimate Std. Error         df t value Pr(>|t|)
(Intercept)    -0.7261391  0.0962871 54.0000012  -7.541 5.44e-10 ***
PC1_Musical_T0 -0.0005217  0.0001082 54.0000012  -4.821 1.20e-05 ***
---
Signif. codes:  0 ‘***’ 0.001 ‘**’ 0.01 ‘*’ 0.05 ‘.’ 0.1 ‘ ’ 1

Correlation of Fixed Effects:
            (Intr)
PC1_Mscl_T0 0.041
```

3) EDT

```
Linear mixed model fit by maximum likelihood . t-tests use Satterthwaite's
  method [lmerModLmerTest]
Formula: EDT ~ PC1_Musical_T0 + (1 | ID)

      AIC       BIC    logLik -2*log(L)  df.resid
    299.5     310.3    -145.8     291.5       104

Scaled residuals:
    Min      1Q  Median      3Q     Max
-2.2060 -0.5498 -0.1399  0.7466  1.7830

Random effects:
 Groups   Name        Variance Std.Dev.
 ID       (Intercept) 0.0000   0.0000
 Residual             0.8707   0.9331
Number of obs: 108, groups:  ID, 54

Fixed effects:
                 Estimate Std. Error         df t value Pr(>|t|)
(Intercept)     8.805e-02  8.986e-02  1.080e+02   0.980    0.329
PC1_Musical_T0 -4.711e-04  1.010e-04  1.080e+02  -4.664 8.91e-06 ***
---
Signif. codes:  0 ‘***’ 0.001 ‘**’ 0.01 ‘*’ 0.05 ‘.’ 0.1 ‘ ’ 1

Correlation of Fixed Effects:
            (Intr)
PC1_Mscl_T0 0.041
```

4) RAT

```
Linear mixed model fit by maximum likelihood . t-tests use Satterthwaite's
  method [lmerModLmerTest]
Formula: RAT ~ PC1_Musical_T0 + (1 | ID)

      AIC       BIC    logLik -2*log(L)  df.resid
    209.7     220.4    -100.8     201.7       104

Scaled residuals:
     Min       1Q   Median       3Q      Max
-2.14901 -0.58562  0.03593  0.57175  1.66449

Random effects:
 Groups   Name        Variance Std.Dev.
 ID       (Intercept) 0.1308   0.3617
 Residual             0.2701   0.5197
Number of obs: 108, groups:  ID, 54

Fixed effects:
                 Estimate Std. Error         df t value Pr(>|t|)
(Intercept)    -9.977e-01  7.023e-02  5.400e+01 -14.208  < 2e-16 ***
PC1_Musical_T0 -5.858e-04  7.892e-05  5.400e+01  -7.423 8.48e-10 ***
---
Signif. codes:  0 ‘***’ 0.001 ‘**’ 0.01 ‘*’ 0.05 ‘.’ 0.1 ‘ ’ 1
```
