## Supplementary material for "Promoting Healthy Ageing through Choral Training: A 9-Month Multidomain Intervention (MultiMusic) Enhances Neural Processing Speed in Community-Dwelling Older Adults": R_outputs_amplitude.docx

1) Final model N2P2 Amplitude

```
Linear mixed model fit by ML (lmerTest)
Formula: P2N2_ampl ~ PC1_P300_amplitude_T0 + (1 | ID)

AIC = 964.17, BIC = 977.60, logLik = -478.09, df.resid = 208

Random effects:
 ID (Intercept) Var = 2.041 (SD = 1.429)
 Residual              4.039 (SD = 2.010)

Fixed effects:
                          Estimate   SE        df   t       p
(Intercept)              4.033e+00 0.2401  53.0  16.80  < .001 ***
PC1_P300_amplitude_T0    1.128e-03 0.000281 53.0  4.01  0.000191 ***
```

2) Final model N2P3 Amplitude

```
Linear mixed model fit by maximum likelihood . t-tests use Satterthwaite's
  method [lmerModLmerTest]
Formula: N2P3_ampl ~ PC1_P300_amplitude_T0 + Time + (1 | ID)

      AIC       BIC    logLik -2*log(L)  df.resid
    857.5     873.8    -423.7     847.5       188

Scaled residuals:
     Min       1Q   Median       3Q      Max
-2.39338 -0.64270 -0.04893  0.63923  2.16365

Random effects:
 Groups   Name        Variance Std.Dev.
 ID       (Intercept) 2.671    1.634
 Residual             3.243    1.801
Number of obs: 193, groups:  ID, 53

Fixed effects:
                        Estimate Std. Error         df t value Pr(>|t|)
(Intercept)            5.042e+00  2.864e-01  7.371e+01  17.601  < 2e-16 ***
PC1_P300_amplitude_T0  1.231e-03  3.006e-04  4.813e+01   4.094 0.000161 ***
TimeT1                -1.110e+00  2.672e-01  1.462e+02  -4.156 5.48e-05 ***
---
Signif. codes:  0 ‘***’ 0.001 ‘**’ 0.01 ‘*’ 0.05 ‘.’ 0.1 ‘ ’ 1

Correlation of Fixed Effects:
            (Intr) PC1_P3
PC1_P300__T  0.034
TimeT1      -0.414  0.002

Type III Analysis of Variance Table with Satterthwaite's method
                      Sum Sq Mean Sq NumDF   DenDF F value    Pr(>F)
PC1_P300_amplitude_T0 54.350  54.350     1  48.132  16.761 0.0001612 ***
Time                  56.016  56.016     1 146.182  17.275 5.484e-05 ***
---
Signif. codes:  0 ‘***’ 0.001 ‘**’ 0.01 ‘*’ 0.05 ‘.’ 0.1 ‘ ’ 1
```
