## Supplementary material for "Promoting Healthy Ageing through Choral Training: A 9-Month Multidomain Intervention (MultiMusic) Enhances Neural Processing Speed in Community-Dwelling Older Adults": R_outputs_latency.docx

```
1)Final model N2P3 interpeak latency

Linear mixed model fit by maximum likelihood . t-tests use Satterthwaite's
  method [lmerModLmerTest]
Formula: n2p3 ~ N2P3_T0 + Time * Groups + (1 | ID)

      AIC       BIC    logLik -2*log(L)  df.resid
   3366.3    3402.8   -1674.1    3348.3       419

Scaled residuals:
     Min       1Q   Median       3Q      Max
-3.03496 -0.56991  0.00136  0.58285  3.05351

Random effects:
 Groups   Name        Variance Std.Dev.
 ID       (Intercept)  17.13    4.139
 Residual             133.88   11.571
Number of obs: 428, groups:  ID, 54

Fixed effects:
                              Estimate Std. Error        df t value
(Intercept)                   17.99356    4.25964  59.45612   4.224
N2P3_T0                        0.67455    0.07147  54.15640   9.438
TimeT1                       -21.12113    1.82948 374.45930 -11.545
GroupsControl_Active          -5.48293    2.60594  96.71023  -2.104
GroupsControl_Passive         -3.77275    2.52566 105.69090  -1.494
TimeT1:GroupsControl_Active   20.20290    2.68570 379.38007   7.522
TimeT1:GroupsControl_Passive  17.20425    2.74421 374.45930   6.269
                                         Pr(>|t|)
(Intercept)                     0.000083525859592 ***
N2P3_T0                         0.000000000000494 ***
TimeT1                       < 0.0000000000000002 ***
GroupsControl_Active                        0.038 *
GroupsControl_Passive                       0.138
TimeT1:GroupsControl_Active     0.000000000000393 ***
TimeT1:GroupsControl_Passive    0.000000000999644 ***
---
Signif. codes:  0 ‘***’ 0.001 ‘**’ 0.01 ‘*’ 0.05 ‘.’ 0.1 ‘ ’ 1

Correlation of Fixed Effects:
            (Intr) N2P3_T TimeT1 GrpC_A GrpC_P TT1:GC_A
N2P3_T0     -0.928
TimeT1      -0.215  0.000
GrpsCntrl_A -0.657  0.462  0.351
GrpsCntrl_P -0.539  0.328  0.362  0.536
TmT1:GrpC_A  0.137  0.010 -0.681 -0.500 -0.243
TmT1:GrpC_P  0.143  0.000 -0.667 -0.234 -0.543  0.454

Type III Analysis of Variance Table with Satterthwaite's method
             Sum Sq Mean Sq NumDF  DenDF F value              Pr(>F)
N2P3_T0     11925.4 11925.4     1  54.16  89.076 0.00000000000049440 ***
Time         7914.4  7914.4     1 377.59  59.116 0.00000000000013016 ***
Groups        808.1   404.1     2  54.44   3.018             0.05715 .
Time:Groups  8948.8  4474.4     2 377.56  33.421 0.00000000000004328 ***
---
Signif. codes:  0 ‘***’ 0.001 ‘**’ 0.01 ‘*’ 0.05 ‘.’ 0.1 ‘ ’ 1

emmeans Time x Groups
Time Groups          emmean   SE  df lower.CL upper.CL
 T0   Choir             49.2 1.77 110     45.7     52.7
 T1   Choir             28.0 1.77 110     24.5     31.6
 T0   Control_Active    43.7 1.82 114     40.1     47.3
 T1   Control_Active    42.8 1.87 120     39.1     46.5
 T0   Control_Passive   45.4 1.84 123     41.8     49.0
 T1   Control_Passive   41.5 1.84 123     37.8     45.1

Degrees-of-freedom method: kenward-roger
Confidence level used: 0.95

Post-hoc tests Within Groups

contrast        estimate   SE  df t.ratio p.value
 Choir_T1vsT0     -21.121 1.84 377 -11.498 <0.0001
 CtrlAct_T1vsT0    -0.918 1.98 387  -0.465  1.0000
 CtrlPass_T1vsT0   -3.917 2.05 377  -1.907  0.1717

Degrees-of-freedom method: kenward-roger
P value adjustment: bonferroni method for 3 tests

Post-hoc tests Between Groups

Time = T0:
 contrast                         estimate   SE  df t.ratio p.value
 Choir - Control_Active               5.48 2.68 103   2.042  0.1310
 Choir - Control_Passive              3.77 2.60 113   1.452  0.4481
 Control_Active - Control_Passive    -1.71 2.54 122  -0.673  1.0000

Time = T1:
 contrast                         estimate   SE  df t.ratio p.value
 Choir - Control_Active             -14.72 2.73 107  -5.400 <0.0001
 Choir - Control_Passive            -13.43 2.60 113  -5.168 <0.0001
 Control_Active - Control_Passive     1.29 2.58 125   0.500  1.0000

Degrees-of-freedom method: kenward-roger
P value adjustment: bonferroni method for 3 tests


2)Final model P2 LATENCY

Linear mixed model fit by maximum likelihood . t-tests use Satterthwaite's
  method [lmerModLmerTest]
Formula: P2_lat ~ PC1_P300_latency_T0 + Time + Groups + Side + (1 | ID)

      AIC       BIC    logLik -2*log(L)  df.resid
   2027.8    2054.6   -1005.9    2011.8       204

Scaled residuals:
    Min      1Q  Median      3Q     Max
-3.1097 -0.6023 -0.0416  0.6475  2.0905

Random effects:
 Groups   Name        Variance Std.Dev.
 ID       (Intercept) 119.9    10.95
 Residual             677.0    26.02
Number of obs: 212, groups:  ID, 53

Fixed effects:
                        Estimate Std. Error         df t value
(Intercept)           296.875587   4.640826 101.113293  63.970
PC1_P300_latency_T0     0.015841   0.002827  53.000000   5.603
TimeT1                  0.839375   7.973235 159.000000   0.105
Groups2                11.030389   6.446104  53.000000   1.713
Groups3                12.935065   6.332918  53.000000   2.042
Sidesx                -12.475274   7.432194 159.000000  -1.678
                       Pr(>|t|)
(Intercept)           < 0.0000000000000002 ***
PC1_P300_latency_T0            0.0000007695 ***
TimeT1                         0.91610
Groups2                        0.09242 .
Groups3                        0.04603 *
Sidesx                         0.09565 .
---
Signif. codes:  0 ‘***’ 0.001 ‘**’ 0.01 ‘*’ 0.05 ‘.’ 0.1 ‘ ’ 1
Type III Analysis of Variance Table with Satterthwaite's method
                     Sum Sq Mean Sq NumDF DenDF F value       Pr(>F)
PC1_P300_latency_T0 21252.7 21252.7     1    53 31.3929 0.0000007695 ***
Time                    7.5     7.5     1   159  0.0111      0.91610
Groups                842.3   421.1     2    53  0.6221      0.54070
Side                 1911.1  1911.1     1   159  2.8230      0.09489 .
---
Signif. codes:  0 ‘***’ 0.001 ‘**’ 0.01 ‘*’ 0.05 ‘.’ 0.1 ‘ ’ 1


3)Final N2 mixed model

Linear mixed model fit by maximum likelihood . t-tests use Satterthwaite's
  method [lmerModLmerTest]
Formula: N2_lat ~ PC1_P300_latency_T0 + Time * Groups + Side + (1 | ID)
   Data: n2_long

      AIC       BIC    logLik -2*log(L)  df.resid
   1926.6    1960.2    -953.3    1906.6       202

Scaled residuals:
    Min      1Q  Median      3Q     Max
-2.1931 -0.6909 -0.1172  0.6320  3.1230

Random effects:
 Groups   Name        Variance Std.Dev.
 ID       (Intercept)  82.32    9.073
 Residual             406.33   20.158
Number of obs: 212, groups:  ID, 53

Fixed effects:
                               Estimate Std. Error         df t value
(Intercept)                  330.522849   4.078243 134.293133  81.045
PC1_P300_latency_T0            0.013193   0.002255  53.000000   5.851
TimeT1                         8.447500   4.507359 159.000000   1.874
GroupsControl_Active          10.973078   5.827848 108.098691   1.883
GroupsControl_Passive         16.628414   5.683547 115.386711   2.926
Sidesx                         0.709481   2.768849 159.000000   0.256
TimeT1:GroupsControl_Active  -15.846912   6.649646 159.000000  -2.383
TimeT1:GroupsControl_Passive -15.830156   6.761038 159.000000  -2.341
                                         Pr(>|t|)
(Intercept)                  < 0.0000000000000002 ***
PC1_P300_latency_T0                   0.000000312 ***
TimeT1                                    0.06274 .
GroupsControl_Active                      0.06241 .
GroupsControl_Passive                     0.00414 **
Sidesx                                    0.79810
TimeT1:GroupsControl_Active              0.01821 *
TimeT1:GroupsControl_Passive             0.02057 *
---
Signif. codes:  0 ‘***’ 0.001 ‘**’ 0.01 ‘*’ 0.05 ‘.’ 0.1 ‘ ’ 1

Type III Analysis of Variance Table with Satterthwaite's method
                     Sum Sq Mean Sq NumDF DenDF F value       Pr(>F)
PC1_P300_latency_T0 13909.7 13909.7     1    53 34.2330 0.0000003124 ***
Time                  234.2   234.2     1   159  0.5764      0.44884
Groups               1504.6   752.3     2    53  1.8515      0.16702
Side                   26.7    26.7     1   159  0.0657      0.79810
Time:Groups          3124.0  1562.0     2   159  3.8442      0.02342 *
---
Signif. codes:  0 ‘***’ 0.001 ‘**’ 0.01 ‘*’ 0.05 ‘.’ 0.1 ‘ ’ 1

Emmeans Time x Groups

Time Groups          emmean   SE  df lower.CL upper.CL
 T0   Choir              331 3.96 122      323      339
 T1   Choir              339 3.96 122      332      347
 T0   Control_Active     342 4.36 119      333      351
 T1   Control_Active     335 4.36 119      326      343
 T0   Control_Passive    348 4.36 124      339      356
 T1   Control_Passive    340 4.36 124      332      349

Results are averaged over the levels of: Side
Degrees-of-freedom method: kenward-roger
Confidence level used: 0.95
Post hoc tests Within Groups
Groups = Choir:
 contrast estimate   SE  df t.ratio p.value
 T0 - T1     -8.45 4.57 163  -1.850  0.0661

Groups = Control_Active:
 contrast estimate   SE  df t.ratio p.value
 T0 - T1      7.40 4.95 163   1.494  0.1370

Groups = Control_Passive:
 contrast estimate   SE  df t.ratio p.value
 T0 - T1      7.38 5.10 163   1.446  0.1500

Results are averaged over the levels of: Side
Degrees-of-freedom method: kenward-roger

Post-hoc tests Between Groups

Time = T0:
 contrast                         estimate   SE  df t.ratio p.value
 Choir - Control_Active            -10.973 6.01 116  -1.826  0.2113
 Choir - Control_Passive           -16.628 5.86 124  -2.839  0.0159
 Control_Active - Control_Passive   -5.655 6.21 120  -0.911  1.0000

Time = T1:
 contrast                         estimate   SE  df t.ratio p.value
 Choir - Control_Active              4.874 6.01 116   0.811  1.0000
 Choir - Control_Passive            -0.798 5.86 124  -0.136  1.0000
 Control_Active - Control_Passive   -5.672 6.21 120  -0.914  1.0000

Results are averaged over the levels of: Side
Degrees-of-freedom method: kenward-roger
P value adjustment: bonferroni method for 3 tests
```

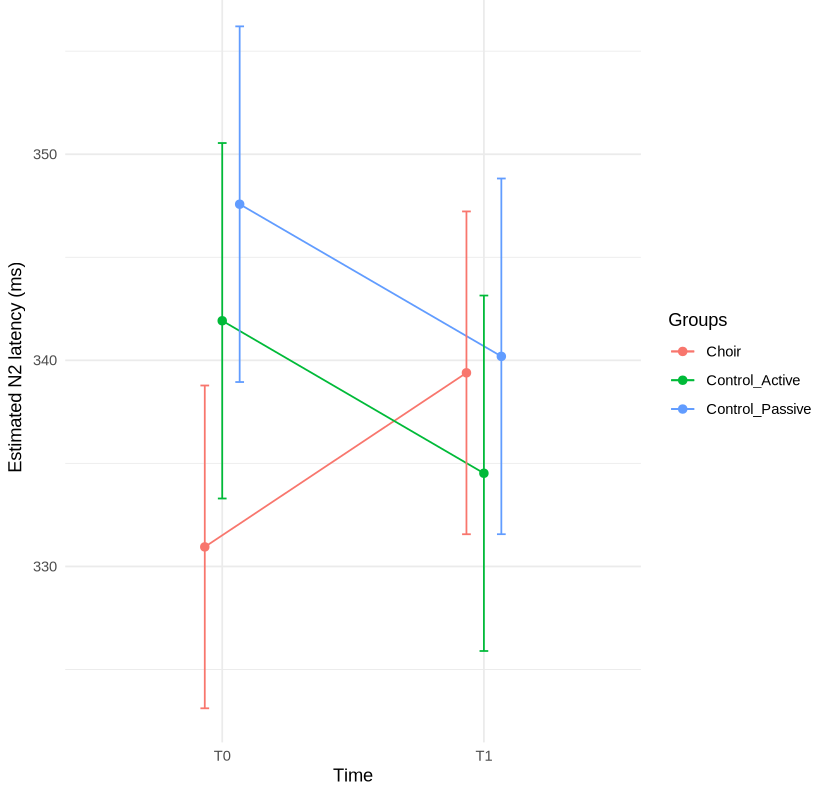

```
4)Joint mixed model N2P3 - N2 latency

Linear mixed model fit by maximum likelihood . t-tests use Satterthwaite's
  method [lmerModLmerTest]
Formula: Latency ~ PC1_P300_latency_T0 + Outcome * Time * Groups + Side +
    (1 | ID)


      AIC       BIC    logLik -2*log(L)  df.resid
   3779.5    3844.3   -1873.7    3747.5       410

Scaled residuals:
    Min      1Q  Median      3Q     Max
-2.6558 -0.6541 -0.1127  0.6271  3.5389

Random effects:
 Groups   Name        Variance Std.Dev.
 ID       (Intercept)   8.801   2.967
 Residual             379.061  19.469
Number of obs: 426, groups:  ID, 54

Fixed effects:
                                             Estimate  Std. Error          df
(Intercept)                                328.414553    3.307724  358.029335
PC1_P300_latency_T0                          0.005215    0.001245   53.294765
OutcomeN2_P3                              -271.702125    4.353512  372.608805
TimeT1                                       8.447500    4.353512  372.608805
GroupsControl_Active                        17.025966    4.739499  324.452264
GroupsControl_Passive                       18.155275    4.729596  345.070399
Sidesx                                       0.224695    1.886600  372.608805
OutcomeN2_P3:TimeT1                        -29.568625    6.156795  372.608805
OutcomeN2_P3:GroupsControl_Active          -37.676640    6.376505  375.669419
OutcomeN2_P3:GroupsControl_Passive         -30.745844    6.530267  372.608805
TimeT1:GroupsControl_Active                -15.846912    6.422677  372.608805
TimeT1:GroupsControl_Passive               -15.830156    6.530267  372.608805
OutcomeN2_P3:TimeT1:GroupsControl_Active    35.500773    9.050447  374.134667
OutcomeN2_P3:TimeT1:GroupsControl_Passive   33.034406    9.235193  372.608805
                                          t value             Pr(>|t|)
(Intercept)                                99.287 < 0.0000000000000002 ***
PC1_P300_latency_T0                         4.188             0.000106 ***
OutcomeN2_P3                              -62.410 < 0.0000000000000002 ***
TimeT1                                      1.940             0.053087 .
GroupsControl_Active                        3.592             0.000378 ***
GroupsControl_Passive                       3.839             0.000147 ***
Sidesx                                      0.119             0.905260
OutcomeN2_P3:TimeT1                        -4.803        0.00000227285 ***
OutcomeN2_P3:GroupsControl_Active          -5.909        0.00000000774 ***
OutcomeN2_P3:GroupsControl_Passive         -4.708        0.00000353069 ***
TimeT1:GroupsControl_Active                -2.467             0.014061 *
TimeT1:GroupsControl_Passive               -2.424             0.015821 *
OutcomeN2_P3:TimeT1:GroupsControl_Active    3.923             0.000104 ***
OutcomeN2_P3:TimeT1:GroupsControl_Passive   3.577             0.000393 ***
---
Signif. codes:  0 ‘***’ 0.001 ‘**’ 0.01 ‘*’ 0.05 ‘.’ 0.1 ‘ ’ 1

===== anova(mod_joint_N2_N2P3, type = 3, ddf = "Satterthwaite") =====
Type III Analysis of Variance Table with Satterthwaite's method
                     Sum Sq Mean Sq NumDF  DenDF    F value
PC1_P300_latency_T0    6648    6648     1  53.29    17.5380
Outcome             9364695 9364695     1 373.58 24704.9662
Time                   3162    3162     1 373.58     8.3411
Groups                  961     480     2  53.65     1.2676
Side                      5       5     1 372.61     0.0142
Outcome:Time           1193    1193     1 373.58     3.1468
Outcome:Groups         7911    3956     2 373.57    10.4350
Time:Groups              68      34     2 373.57     0.0896
Outcome:Time:Groups    7375    3688     2 373.57     9.7283
                                   Pr(>F)
PC1_P300_latency_T0             0.0001062 ***
Outcome             < 0.00000000000000022 ***
Time                            0.0041014 **
Groups                          0.2898003
Side                            0.9052600
Outcome:Time                    0.0768915 .
Outcome:Groups                 0.00003892 ***
Time:Groups                     0.9143378
Outcome:Time:Groups            0.00007610 ***
---
Signif. codes:  0 ‘***’ 0.001 ‘**’ 0.01 ‘*’ 0.05 ‘.’ 0.1 ‘ ’ 1

Effect size estimated by Side
Outcome  Side   F_TG   NumDF  DenDF   p_TG      eta2p_Time_Group
N2_lat   dx     2.29   2      53      0.112    0.079
N2_lat   sx     1.17   2      106     0.315    0.022

N2_P3    dx     8.09   2      53.9    0.00085  0.231
N2_P3    sx     8.18   2      52.4    0.00081  0.238


5) Final model P3 latency

Linear mixed model fit by maximum likelihood
Formula: P3_lat ~ PC1_P300_latency_T0 + (1 | ID)

AIC = 2478.1,  BIC = 2491.2,  logLik = -1235.1,  df.resid = 190

Random effects:
 Groups   Name        Variance Std.Dev.
 ID       (Intercept)   671.5   25.91
 Residual             19184.6  138.51
Number of obs: 194, groups: ID, 53

Fixed effects:
                     Estimate Std. Error   df t value Pr(>|t|)
(Intercept)         389.52      10.59   56.37  36.78  < 2e-16 ***
PC1_P300_latency_T0   0.0105     0.0121 58.28   0.87   0.39
Type III Analysis of Variance Table
                    Sum Sq Mean Sq NumDF  DenDF F value Pr(>F)
PC1_P300_latency_T0  14403   14403     1 58.283  0.7508 0.3898


6) Final model P2N2 latency

Linear mixed model fit by ML
Formula: P2_N2 ~ PC1_P300_latency_T0 + Intensity + Time + Side + (1 | ID)

AIC = 1627.7, BIC = 1650.8, logLik = -806.9, df.resid = 192
Random effects:
 ID (Intercept) Var = 16.14 (SD = 4.02)
 Residual              180.25 (SD = 13.43)
 N obs = 199, ID = 53

Fixed effects:
                          Estimate  SE        df   t       p
(Intercept)               38.60    2.20   111.75  17.58  < .001 ***
PC1_P300_latency_T0      -0.00467  0.00129 51.86  -3.63  0.00064 ***
Intensity                -1.06     0.45   50.54  -2.36  0.0222 *
TimeT1                   -2.76     1.91  155.63  -1.44  0.151
SideRight                 3.68     1.90  148.34   1.94  0.0548 .


ANOVA type III (Satterthwaite)
                     F       p
PC1_P300_latency_T0 13.21  0.00064 ***
Intensity            5.57  0.0222 *
Time                 2.08  0.1512
Side                 3.75  0.0548 .
```
