## Supplementary material for "Promoting Healthy Ageing through Choral Training: A 9-Month Multidomain Intervention (MultiMusic) Enhances Neural Processing Speed in Community-Dwelling Older Adults": Selfy_MPI.docx

1) Final Mixed model with Groups

```
Linear mixed model fit by maximum likelihood . t-tests use Satterthwaite's
  method [lmerModLmerTest]
Formula: Selfy ~ Time + Groups + (1 | ID)

      AIC       BIC    logLik -2*log(L)  df.resid
   -193.7    -177.6     102.8    -205.7       102

Scaled residuals:
    Min      1Q  Median      3Q     Max
-1.8355 -0.4283 -0.0143  0.4416  3.3222

Random effects:
 Groups   Name        Variance Std.Dev.
 ID       (Intercept) 0.008211 0.09061
 Residual             0.003765 0.06136
Number of obs: 108, groups:  ID, 54

Fixed effects:
                       Estimate Std. Error        df t value Pr(>|t|)
(Intercept)            0.209500   0.023228 61.424595   9.019 7.46e-13 ***
TimeT1                -0.020000   0.011808 54.000000  -1.694   0.0961 .
GroupsControl_Active  -0.001167   0.032641 54.000000  -0.036   0.9716
GroupsControl_Passive -0.087625   0.033697 54.000000  -2.600   0.0120 *
---
Signif. codes:
0 ‘***’ 0.001 ‘**’ 0.01 ‘*’ 0.05 ‘.’ 0.1 ‘ ’ 1

Correlation of Fixed Effects:
            (Intr) TimeT1 GrpC_A
TimeT1      -0.254
GrpsCntrl_A -0.666  0.000
GrpsCntrl_P -0.645  0.000  0.459
Analysis of Deviance Table (Type III Wald chisquare tests)

Response: Selfy
              Chisq Df Pr(>Chisq)
(Intercept) 81.3498  1    < 2e-16 ***
Time         2.8687  1    0.09032 .
Groups       8.4587  2    0.01456 *
---
Signif. codes:
0 ‘***’ 0.001 ‘**’ 0.01 ‘*’ 0.05 ‘.’ 0.1 ‘ ’ 1

EMMs for Groups


 Groups          emmean     SE   df lower.CL upper.CL
 Choir            0.200 0.0231 57.2   0.1532    0.246
 Control_Active   0.198 0.0244 57.2   0.1495    0.247
 Control_Passive  0.112 0.0258 57.2   0.0601    0.164

Results are averaged over the levels of: Time
Degrees-of-freedom method: kenward-roger
Confidence level used: 0.95


Post-hoc for Groups (Bonferroni, ML)
 contrast                         estimate     SE   df t.ratio p.value
 Choir - Control_Active            0.00117 0.0336 57.2   0.035  1.0000
 Choir - Control_Passive           0.08762 0.0347 57.2   2.527  0.0429
 Control_Active - Control_Passive  0.08646 0.0355 57.2   2.434  0.0542

Results are averaged over the levels of: Time
Degrees-of-freedom method: kenward-roger
P value adjustment: bonferroni method for 3 tests


   2) Mixed model with Time * Groups interaction

Linear mixed model fit by maximum likelihood .
t-tests use Satterthwaite's method [lmerModLmerTest]

Formula: Selfy ~ Selfy_T0 + Time * Groups + (1 | ID)

      AIC       BIC    logLik -2*log(L)  df.resid
   -303.0    -278.9     160.5    -321.0        99

Scaled residuals:
    Min      1Q  Median      3Q     Max
-3.3308 -0.4170  0.0519  0.3535  4.0657

Random effects:
 Groups   Name        Variance Std.Dev.
 ID       (Intercept) 0.000000 0.00000
 Residual             0.002997 0.05475
Number of obs: 108, groups:  ID, 54

Fixed effects:
                              Estimate Std. Error        df t value Pr(>|t|)
(Intercept)                    0.04405    0.01562 108.00000   2.821  0.0057 **
Selfy_T0                       0.79746    0.04458 108.00000  17.889  <2e-16 ***
TimeT1                        -0.03600    0.01731 108.00000  -2.079  0.0399 *
GroupsControl_Active          -0.00467    0.01782 108.00000  -0.262  0.7937
GroupsControl_Passive         -0.01823    0.01880 108.00000  -0.970  0.3343
TimeT1:GroupsControl_Active    0.04378    0.02515 108.00000   1.740  0.0846 .
TimeT1:GroupsControl_Passive   0.00475    0.02597 108.00000   0.183  0.8552
---
Signif. codes:
0 ‘***’ 0.001 ‘**’ 0.01 ‘*’ 0.05 ‘.’ 0.1 ‘ ’ 1

Type III Analysis of Variance Table with Satterthwaite's method

             Sum Sq Mean Sq NumDF DenDF  F value  Pr(>F)
Selfy_T0    0.95916 0.95916     1   108 320.0204 < 2e-16 ***
Time        0.01052 0.01052     1   108   3.5110 0.06367 .
Groups      0.01785 0.00893     2   108   2.9779 0.05509 .
Time:Groups 0.01052 0.00526     2   108   1.7545 0.17789
---
Signif. codes:
0 ‘***’ 0.001 ‘**’ 0.01 ‘*’ 0.05 ‘.’ 0.1 ‘ ’ 1


EMMs Time × Groups

Time Groups          emmean     SE  df lower.CL upper.CL
 T0   Choir            0.190 0.0128 115    0.165    0.215
 T1   Choir            0.154 0.0128 115    0.129    0.179

 T0   Control_Active   0.185 0.0134 115    0.159    0.212
 T1   Control_Active   0.193 0.0134 115    0.167    0.220

 T0   Control_Passive  0.172 0.0144 115    0.143    0.200
 T1   Control_Passive  0.141 0.0144 115    0.112    0.169

Degrees-of-freedom method: kenward-roger
Confidence level used: 0.95

Time contrasts within Groups

Groups = Choir:
 Time_pairwise estimate     SE   df t.ratio p.value
 T0 - T1        0.03600 0.0179 58.3   2.011  0.0490

Groups = Control_Active:
 Time_pairwise estimate     SE   df t.ratio p.value
 T0 - T1       -0.00778 0.0189 58.3  -0.412  0.6817

Groups = Control_Passive:
 Time_pairwise estimate     SE   df t.ratio p.value
 T0 - T1        0.03125 0.0200 58.3   1.561  0.1239

Degrees-of-freedom method: kenward-roger


3) Final mixed model for number of activities

Linear mixed model fit by maximum likelihood
Formula: Selfy_num ~ Selfy_T0 + Time + activities_level + (1 | ID)

      AIC       BIC    logLik -2*log(L)  df.resid
   -300.3    -284.2     156.1    -312.3       102

Random effects:
 Groups   Name        Variance Std.Dev.
 ID       (Intercept) 0.000000 0.000
 Residual             0.003249 0.057

Number of obs: 108, groups:  ID, 54

Fixed effects:
                      Estimate Std. Error         df t value Pr(>|t|)
(Intercept)           0.028563   0.013609 108.000000   2.099   0.0382 *
Selfy_T0              0.823543   0.044976 108.000000  18.315  < 2e-16 ***
TimeT1               -0.019998   0.010976 108.000000  -1.822   0.0711 .
activities_levelLow  -0.006448   0.010712 108.000000  -0.602   0.5482

Signif. codes: 0 ‘***’ 0.001 ‘**’ 0.01 ‘*’ 0.05 ‘.’ 0.1


Type III Analysis of Variance Table with Satterthwaite's method

                  Sum Sq Mean Sq NumDF DenDF  F value  Pr(>F)
Selfy_T0         1.08931 1.08931     1   108 335.2812 < 2e-16 ***
Time             0.01080 0.01080     1   108   3.3242 0.07103 .
activities_level 0.00118 0.00118     1   108   0.3629 0.54818

4) Model with Time * Activities_level interaction

Linear mixed model fit by maximum likelihood . t-tests use Satterthwaite's
  method [lmerModLmerTest]
Formula: Selfy_num ~ Selfy_T0 + Time * activities_level + (1 | ID)

      AIC       BIC    logLik -2*log(L)  df.resid
   -300.1    -281.3     157.0    -314.1       101

Scaled residuals:
    Min      1Q  Median      3Q     Max
-3.3999 -0.3150  0.0065  0.3954  3.9565

Random effects:
 Groups   Name        Variance Std.Dev.
 ID       (Intercept) 0.000000 0.00000
 Residual             0.003196 0.05653
Number of obs: 108, groups:  ID, 54

Fixed effects:
                             Estimate Std. Error         df t value Pr(>|t|)
(Intercept)                  0.036688   0.014805 108.000000   2.478   0.0148 *
Selfy_T0                     0.823543   0.044609 108.000000  18.461   <2e-16 ***
TimeT1                      -0.036250   0.016320 108.000000  -2.221   0.0284 *
activities_levelLow         -0.007867   0.015610 108.000000  -0.504   0.6153
TimeT1:activities_levelLow   0.029250   0.021896 108.000000   1.336   0.1844

(Intercept)                *
Selfy_T0                   ***
TimeT1                     *
activities_levelLow
TimeT1:activities_levelLow
---
Signif. codes:  0 ‘***’ 0.001 ‘**’ 0.01 ‘*’ 0.05 ‘.’ 0.1 ‘ ’ 1

Correlation of Fixed Effects:
            (Intr) Slf_T0 TimeT1 actv_L
Selfy_T0    -0.626
TimeT1      -0.551  0.000
actvts_lvlL -0.656  0.127  0.523
TmT1:ctvt_L  0.411  0.000 -0.745 -0.701

Type III Analysis of Variance Table with Satterthwaite's method
                       Sum Sq Mean Sq NumDF DenDF  F value  Pr(>F)
Selfy_T0              1.08931 1.08931     1   108 340.8214 < 2e-16 ***
Time                  0.01247 0.01247     1   108   3.9017 0.05079 .
activities_level      0.00118 0.00118     1   108   0.3689 0.54490
Time:activities_level 0.00570 0.00570     1   108   1.7846 0.18440
---
Signif. codes:  0 ‘***’ 0.001 ‘**’ 0.01 ‘*’ 0.05 ‘.’ 0.1 ‘ ’ 1

EMMEANS for  Time × activities_level

 Time activities_level emmean     SE  df lower.CL upper.CL
 T0   High              0.188 0.0119 113    0.164    0.211
 T1   High              0.151 0.0119 113    0.128    0.175
 T0   Low               0.180 0.0106 113    0.159    0.201
 T1   Low               0.173 0.0106 113    0.152    0.194

Degrees-of-freedom method: kenward-roger
Confidence level used: 0.95

contrast(emm_interact,
         interaction = "pairwise",
         by = "activities_level",
         adjust = "bonferroni")
r


activities_level = High:
 Time_pairwise estimate     SE   df t.ratio p.value
 T0 - T1         0.0362 0.0167 57.2   2.169  0.0342

activities_level = Low:
 Time_pairwise estimate     SE   df t.ratio p.value
 T0 - T1         0.0070 0.0149 57.2   0.468  0.6413

Degrees-of-freedom method: kenward-roger
```
