## Supplementary material for "Promoting Healthy Ageing through Choral Training: A 9-Month Multidomain Intervention (MultiMusic) Enhances Neural Processing Speed in Community-Dwelling Older Adults": APA_Tables_T0_neurophysiologic_.docx

Neurophysiologic domain - Latency

| **Variable** | **F** | **df** | **p** | **q (FDR)** |
| --- | --- | --- | --- | --- |
| p2_lat_dx | 2.540 | 2, 51 | 0.088 | 0.143 |
| n2_lat_dx | 3.750 | 2, 51 | 0.030 | 0.060 |
| p3_lat_dx | 0.540 | 2, 51 | 0.588 | 0.624 |
| p2_n2_dx | 2.410 | 2, 51 | 0.100 | 0.143 |
| p2_lat_sx | 5.370 | 2, 51 | 0.008 | 0.026 |
| n2_lat_sx | 3.770 | 2, 51 | 0.030 | 0.060 |
| p3_lat_sx | 0.610 | 2, 51 | 0.546 | 0.624 |
| p2_n2_sx | 2.210 | 2, 51 | 0.120 | 0.150 |
| n2p3_dx | 4.160 | 2, 51 | 0.021 | 0.035 |
| n2p3_sx | 4.240 | 2, 51 | 0.020 | 0.035 |

Neurophysiologic domain - Amplitude

| **Variable** | **F** | **df** | **p** | **q (FDR)** |
| --- | --- | --- | --- | --- |
| n2_ampl_dx | 3.330 | 2, 51 | 0.044 | 0.175 |
| p3_ampl_dx | 1.080 | 2, 51 | 0.346 | 0.439 |
| n2_ampl_sx | 1.040 | 2, 51 | 0.362 | 0.439 |
| p3_ampl_sx | 0.840 | 2, 50 | 0.439 | 0.439 |
