## Supplementary material for "Promoting Healthy Ageing through Choral Training: A 9-Month Multidomain Intervention (MultiMusic) Enhances Neural Processing Speed in Community-Dwelling Older Adults": T0_ANOVAS.docx

### 1. T0 One-way ANOVA results by domain (BH-FDR within domain)

At T0, 7 measures survived the Benjamini–Hochberg correction applied within domains. The Choir scored lower than both control groups on CRI-q (pFDR < .001), MoCA (pFDR 0.019), and MIQ (pFDR 0.029). In the musical domain, the Choir showed lower rhythmic aptitude on RAT (pFDR 0.018). Electrophysiologically, the Choir showed longer left-hemisphere P2 latency (pFDR 0.036; higher latency = worse) and higher interpeak N2–P3 latency values bilaterally (right: pFDR 0.002; left: pFDR 0.036).

#### 1.1. COGNITIVE DOMAIN

| Measure | N | F (df1, df2) | p | pFDR |
| --- | --- | --- | --- | --- |
| CRI_Q | 54 | F(2, 51) = 20.10 | < .001 | < .001 |
| MIQ | 54 | F(2, 51) = 3.80 | 0.029 | 0.029 |
| MoCa | 54 | F(2, 51) = 4.74 | 0.013 | 0.019 |

#### 1.2. MUSICAL DOMAIN

| Measure | N | F (df1, df2) | p | pFDR |
| --- | --- | --- | --- | --- |
| EDT | 54 | F(2, 51) = 1.29 | 0.283 | 0.377 |
| MDT | 54 | F(2, 51) = 0.14 | 0.869 | 0.869 |
| MPT | 54 | F(2, 51) = 1.59 | 0.214 | 0.377 |
| RAT | 54 | F(2, 51) = 6.00 | 0.005 | 0.018 |

#### 1.3. AUDIOMETRIC DOMAIN

| Measure | N | F (df1, df2) | p | pFDR |
| --- | --- | --- | --- | --- |
| Intellection_left | 54 | F(2, 51) = 0.68 | 0.514 | 0.679 |
| Intellection_right | 54 | F(2, 51) = 0.48 | 0.622 | 0.679 |
| Matrix_test | 52 | F(2, 49) = 0.51 | 0.605 | 0.679 |
| PTA_left | 54 | F(2, 51) = 0.94 | 0.397 | 0.679 |
| PTA_right | 54 | F(2, 51) = 0.39 | 0.679 | 0.679 |
| SRT_left | 54 | F(2, 51) = 1.40 | 0.255 | 0.679 |
| SRT_right | 54 | F(2, 51) = 0.94 | 0.397 | 0.679 |
