## Supplementary figures and images for "Promoting Healthy Ageing through Choral Training: A 9-Month Multidomain Intervention (MultiMusic) Enhances Neural Processing Speed in Community-Dwelling Older Adults"

### new_cluster.png

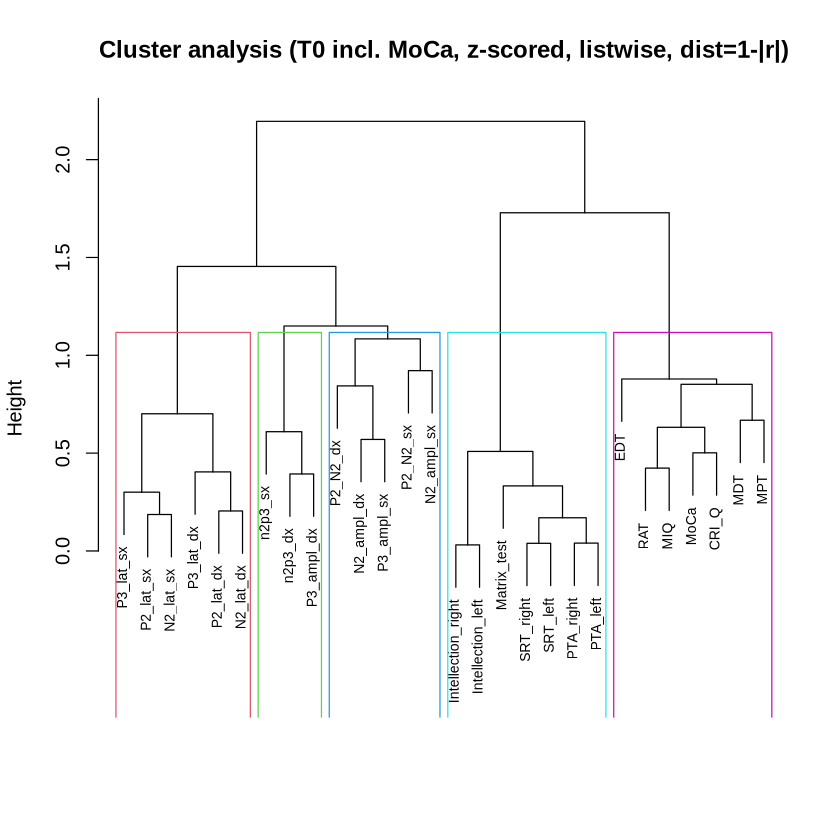

### T0_interdomain_significant_barplot_ENG.png

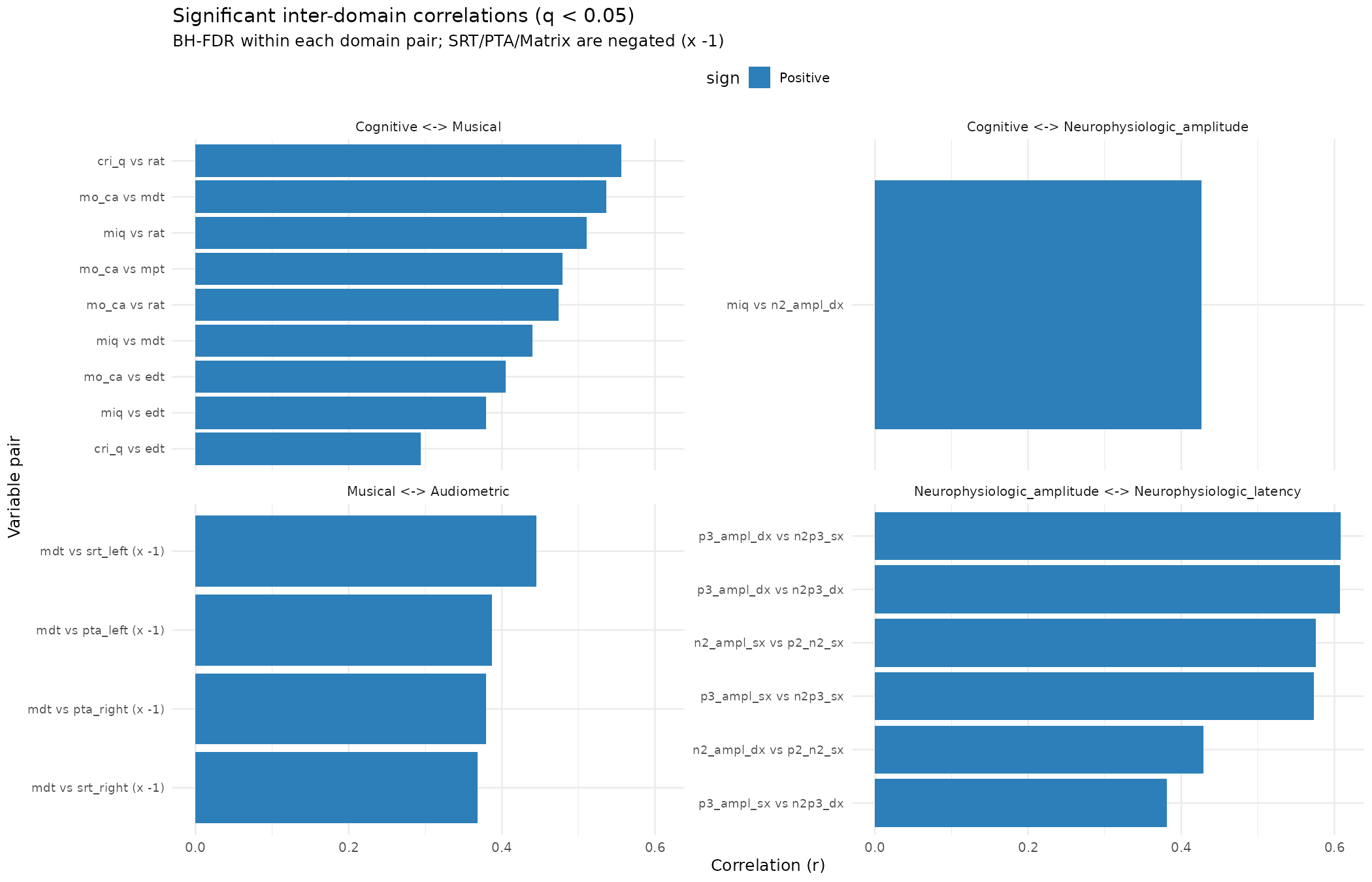

### t1_correlations.png

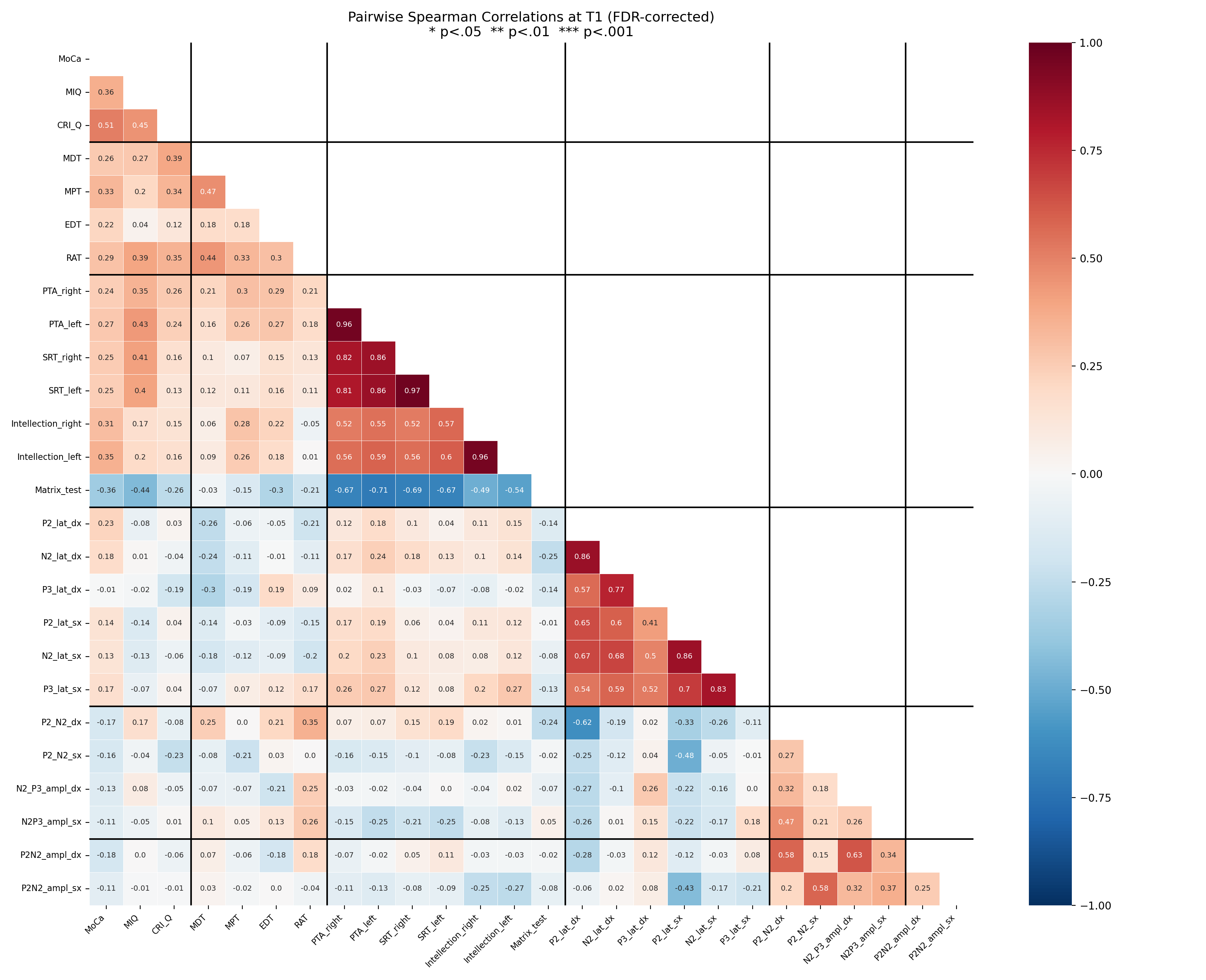
